# TLR7 inhibition limits cardiac ischemic injury by disrupting ITGAM-dependent immune–endothelial interaction

**DOI:** 10.64898/2026.01.17.698774

**Authors:** Yijia Li, Yang Yang, Chanhee Park, Boyang Ren, Ruoxing Li, Amol Shetty, Brittney Williams, Ziyi Li, Lin Zou, Wei Chao

## Abstract

Percutaneous coronary intervention (PCI) limits ischemic myocardial infarction but also triggers ischemia-reperfusion (I/R) injury in part driven by innate immune activation. Here, we identify Toll-like receptor 7 (TLR7), an endosomal sensor of single-stranded RNA, as a mediator of post-ischemic inflammation and myocardial damage. Pharmacological inhibition of TLR7 with enpatoran reduced myocardial inflammation and infarct size and improved cardiac function in a mouse model of I/R injury when administered before, during, or shortly after ischemia. Single-nucleus RNA sequencing revealed coordinated post-I/R expansion of myeloid cells and distinct inflammatory endothelial subsets enriched for leukocyte-interaction programs, with marked upregulation of *Itgam* in cardiac leukocytes and endothelial cells and in circulating monocytes. Circulating *ITGAM^+^* monocytes were similarly increased in patients with ST-segment elevation myocardial infarction 24 hours after coronary stenting. Mechanistically, TLR7 activation induced *Itgam* expression in endothelial cells and leukocytes and promoted their adhesion via ITGAM-ICAM1 interaction under physiological shear stress, whereas ITGAM neutralization disrupted this interaction, reduced immune cell infiltration, and limited ischemic injury. These findings define a TLR7–ITGAM signaling axis as a key driver of endothelial–leukocyte crosstalk in myocardial I/R injury and support TLR7 inhibition as a promising therapeutic strategy to mitigate acute myocardial infarction.

## INTRODUCTION

Cardiovascular disease, such as acute coronary syndrome, remains the leading cause of death and leads to immense health and economic burdens worldwide(*1*). In the U.S., about 805,000 people experience a heart attack every year. Of these, 605,000 are a first heart attack(*2*). Current reperfusion therapy, such as percutaneous coronary intervention (PCI) (*e.g.,* stenting), remains the most effective strategy to limit myocardial infarct (MI) size and preserve cardiac function(*3, 4*). However, reperfusion itself can paradoxically cause damage to myocardium, termed ischemia-reperfusion (I/R) injury(*4*), which is associated with subsequent lethal cardiovascular events, such as recurrent MI, arrhythmia, and cardiac arrest in 5-6% of patients who have undergone PCI(*3, 4*). Therefore, new cardiac protective strategies for lethal reperfusion injury are urgently needed.

Myocardial I/R injury is a complex process driven by multiple mechanisms(*4, 5*), including injurious innate immune activity in response to cell-released damage-associated molecular patterns (DAMPs)(*6–8*), such as extracellular (ex) nucleic acids. Studies have shown that myocardial I/R triggers the release of cellular nucleic acids into the cardiac interstitial space(*9*) and numerous miRNAs into blood circulation(*10*). Tissue RNA isolated from human and rodent hearts or plasma miRNAs identified via RNA array in mice with ischemic MI prove to be inflammatory and act via Toll-like receptor 7(TLR7)(*10–12*). Administration of RNase(*13, 14*) or a specific nucleic acid-binding nanoprobe(*9*) before and during ischemia reduces cell-free RNA and confers cardioprotection against I/R injury with reduced inflammation, smaller infarct size, and improved left ventricular function. Notably, a similar paradigm is seen in autoimmune diseases, including systemic lupus erythematosus (SLE), rheumatoid arthritis, and psoriasis, in which self ex-RNA activates TLR7–mediated signaling(*15, 16*). Ex-RNA has been targeted for potential treatment of SLE in a recent clinical trial(*17*). Patients with these autoimmune disorders exhibit a significantly higher risk of ischemic MI, likely attributed to chronic inflammation, endothelial dysfunction, and unstable atherosclerotic plaques(*18, 19*). Taken together, these observations support the hypothesis that ex-RNA-driven innate immune signaling mediates myocardial inflammation and contributes to cardiac injury during I/R.

Expressed across immune and non-immune cells, mouse TLR7, or human TLR8, is a single-stranded RNA (ssRNA) sensor located in endosomes that recognizes certain extracellular miRNAs(*20–22*) and viral RNAs(*23, 24*). TLR7 activation elicits inflammatory cytokines, chemokines, and type I interferons. In the heart, local injection of miR-146a-5p directly triggers TLR7-dependent myocardial inflammation and cardiomyocyte dysfunction(*12*), and TLR7 is involved in adverse LV remodeling after MI in mice(*25*). TLR7 signaling is also reportedly associated with autoimmune myocarditis(*26*), conduction abnormalities(*27*). Yet the mechanistic role of TLR7 in cardiac I/R injury remains unknown.

In this study, we tested the therapeutic potential of the selective TLR7/8 antagonist enpatoran in a mouse model of I/R injury using dosing regimens that model clinically relevant reperfusion scenarios encountered in patients with acute myocardial infarction. To elucidate underlying mechanisms, we applied single-nucleus RNA sequencing (snRNA-seq) to define changes in myocardial cellular compositions and cell–type–specific transcriptional programs following I/R and TLR7 inhibition. This approach uncovered coordinated expansion of myeloid cells and distinct inflammatory endothelial subsets enriched for leukocyte-interaction pathways, with prominent upregulation of *Itgam* in cardiac leukocytes and endothelial cells. Complementary in vitro and in vivo studies demonstrated that TLR7 activation promotes leukocyte–endothelial adhesion through induction of the ITGAM–ICAM1 axis under physiological shear stress, whereas ITGAM neutralization attenuated immune cell infiltration and reduced ischemic myocardial injury. Together, these findings establish TLR7 as a druggable upstream regulator of immune–endothelial crosstalk in myocardial I/R injury and support TLR7 inhibition as a translational strategy to limit cardiac ischemic injury.

## RESULTS

### Plasma small RNA-seq identifies an increase in proinflammatory miRNAs after myocardial I/R injury

In a mouse model of cardiac I/R injury(*9, 13*), there was a substantial increase in plasma RNA at 24 hours (**Fig. 1A**). RNA-seq revealed that miRNAs were the predominant small RNA species in plasma, accounting for 82% of total RNA reads in control animals (**Fig. 1B**). Among the 1,915 annotated mouse miRNAs in miRBase, 508 miRNAs were detected with ≥10 raw reads. Of these, 7 miRNAs, miR-29c-5p, miR-3061-3p, miR-741-3p, miR-8116, miR-1843-5p, miR-5113, and miR-8097, were significantly upregulated by >1.5-fold (P < 0.05) in the I/R mice as compared with sham controls (**Fig. 1C**). Notably, four of these miRNAs (miR-29c-5p, miR-3061-3p, miR-741-3p, and miR-8116) harbored at least one of the five UU-rich motifs predictive for pro-inflammatory activity(*21*) (**Fig. 1C, Table S1**), three of them elicited a dose-dependent MIP-2 production in bone marrow-derived macrophages (BMDMs), an effect that was completely blocked by the TLR7 antagonist enpatoran (**Fig. 1D**).

**Figure 1.**
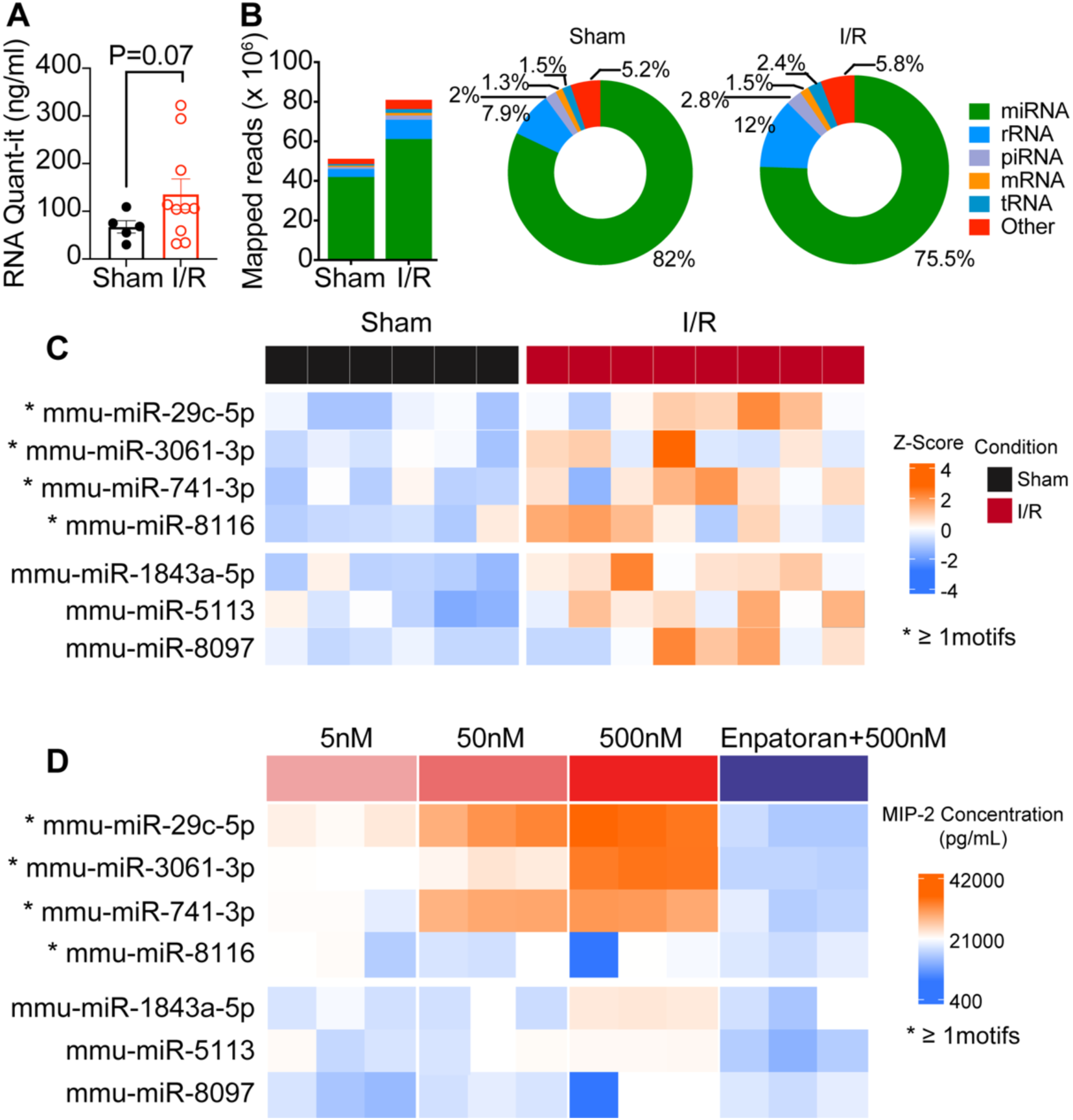
Plasma small RNA-seq and identification of proinflammatory miRNAs in a mouse model of MI. A, Plasma RNA quantifications 24 hours after Sham or I/R procedure (n=5-10). **B,** RNA-seq analysis with mapped read counts and relative percentage of different RNA biotypes in the plasma (n=6-8). **C,** Upregulated plasma miRNAs between I/R vs. Sham identified by RNA-seq. Four miRNAs contain at least one of the five proinflammatory motifs (.uuc; u..uu.; .a. uu.; g.uu.; uu. u.). **D,** MIP-2 production in bone marrow-derived macrophages (BMDM) treated with miRNA mimics. I/R, ischemia-reperfusion**. \*** Indicates miRNAs containing proinflammatory motifs in their sequences (see Table S1).

### Enpatoran, a selective TLR7/8 antagonist, protects the heart from I/R injury

Given the known contribution of ex-RNA and multiple miRNAs to innate inflammation and myocardial I/R injury, we reasoned that targeting the downstream RNA sensor TLR7 would be more efficacious than individual plasma miRNAs. To test this, we examined the effect of enpatoran, a selective TLR7/8 antagonist recently developed by Merck KGaA for treatment of autoimmune diseases(*28*), in a mouse model of cardiac I/R injury. Each mouse received two intraperitoneal injections of enpatoran (5 mg/kg) or saline, two hours apart. Four dosing regimens centered around 45 min of ischemia were adopted (**Fig. 2A**):(1) *Prevention (PP)*: 1^st^ dose given 1 hour before ischemia (45 min); (2) *Treatment (TP)*: 1^st^ dose at 35 minutes during ischemia; (3) *Early rescue (ERP)*: 1^st^ dose at 30 minutes after ischemia; and (4) *Late rescue (LRP)*: 1^st^ dose at 2 hours after ischemia. Enpatoran markedly reduced MI size in the groups of PP, TP, and ERP protocols, achieving 35–76% reductions in myocardial infarction/area-at-risk (MI/AAR), whereas the LRP protocol provided no protection (**Fig. 2B-E**). Area-at-risk was similar across groups (**Fig. 2C**), indicating that ischemic areas were unchanged in the different treatment groups. Plasma cTnI levels were reduced by ∼70% with enpatoran (**Fig. 2F**), further validating the robust cardioprotection by TLR7 inhibition. Baseline echocardiographic parameters were comparable across groups (**Table S2**). I/R injury caused ventricular dilation and impaired systolic and diastolic function, accompanied by reduced cardiac output (CO), LV ejection fraction (LVEF), global longitudinal strain (GLS), and radial strain (GRS), and increased left atrial (LA) size and mitral valve E wave velocity and reversed longitudinal strain rate ratio (E/e′ sr) (**Fig. 2G-O**). Enpatoran administered in the PP, TP, or ERP regimens significantly improved cardiac structure and function, restoring CO, LVEF, GLS, and GRS and reducing LA dilation and E/e′ sr (**Fig. 2G-O**). In contrast, the LRP regimen failed to improve cardiac performance, consistent with the absence of infarct-sparing effect. Of note, enpatoran had no adverse effects on cardiac function in sham mice, and echocardiographic parameters were comparable between the treatment groups (**Table S3**).

**Figure 2.**
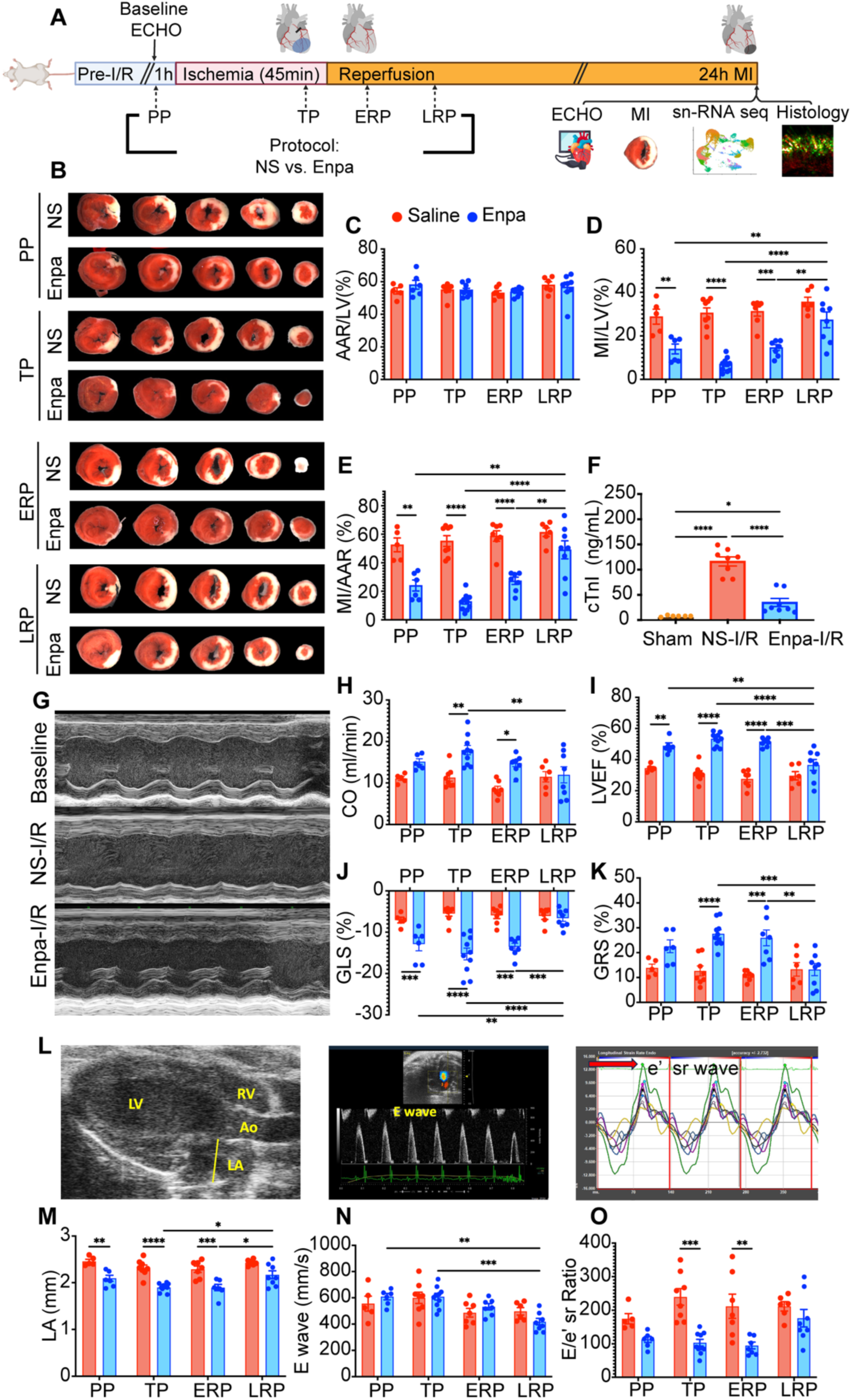
Enpatoran reduces infarct size and preserves cardiac function after I/R. **A**, Study diagram for intraperitoneal administration of enpatoran in cardiac I/R model and experimental endpoints (n = 6-10 mice per group). For each protocol (PP, TP, ERP, or LRP), two doses of enpatoran (5 mg/kg) were administered 2 hours apart; the diagram shows the 1^st^ dose. **B**, Representative TTC-stained images of MI. **C-E**, AAR/LV, MI/LV, and MI/AAR 24h after I/R. **F**, Plasma cTnI following I/R (PP protocol). **G**, M-mode images of echocardiography. **H-K**, Cardiac output (CO) (**H**), left ventricular ejection fraction (LVEF) (**I**), GLS (**J**), and GRS (**K**). **L**, B-mode images showing LA diameter (left panel), E waves (middle panel), and e’ sr waves (right panel). **M-O**, Quantification of LA diameter (**M**), E wave velocity (**N**), and E/e’ sr (**O**). Data shown are means ± SEM. One-way and two-way ANOVA were used to test statistical significance. *P < 0.05, **P < 0.01, ***P < 0.001, ****P < 0.0001. Enpa, enpatoran; NS, normal saline; I/R, ischemia-reperfusion; PP, prevention protocol; TP, treatment protocol; ERP, early rescue protocol; LRP, late rescue protocol; ECHO, echocardiography; AAR, area-at-risk; MI, myocardial infarction; RV, right ventricle; LA, left atrium; Ao, aorta; E wave, mitral valve E wave velocity; E/e’ sr, mitral valve E wave velocity and reversed longitudinal strain rate ratio; GLS, global longitudinal strain; GRS, global radial strain; cTnI, Cardiac troponin I.

To determine whether enpatoran-induced cardioprotection is mediated via TLR7 inhibition, rather than off-target effects, age-matched WT and TLR7-deficient (TLR7 KO) mice underwent I/R injury and received enpatoran or saline following the TP protocol (**Fig. S1A**). Like enpatoran-treated WT mice, TLR7 KO mice had significantly reduced MI/LV and MI/AAR ratios compared with WT mice treated with saline. However, enpatoran provided no additional MI-sparing effect in TLR7 KO mice (**Fig. S1B–C**), suggesting that its cardioprotective effects are TLR7-dependent. Baseline echocardiographic parameters were comparable across groups (**Table S4**). After I/R, NS-treated WT mice developed severe systolic and diastolic dysfunction, including reduced CO, LVEF, and GLS, and increased LA diameter and E/e′ sr (**Fig. S1D–H**). In contrast, NS-TLR7 KO and Enpa-TLR7 KO mice showed similarly preserved cardiac function, further confirming that enpatoran acts through TLR7 inhibition. Of note, comparable cardioprotection was observed in female mice treated with enpatoran (**Fig. S2, Table S5**). Specifically, both enpatoran treatment and TLR7 deficiency significantly reduced infarct size and preserved systolic and diastolic cardiac function after I/R in female mice, as assessed by echocardiographic parameters. Collectively, these results demonstrate that TLR7 inhibition by enpatoran confers a robust, time-sensitive cardioprotection against ischemic MI in a TLR7-dependent manner.

### snRNA-seq reveals expansion of myeloid and endothelial cells with distinct transcriptomic signatures after I/R injury

Given the central role of innate immune activation in cardiac I/R injury, we first analyzed the temporal dynamics of cardiac immune cell infiltration and cytokine expression between 24 hours to 14 days in our mouse model of I/R injury. Flow cytometry revealed a rapid innate immune response, with Ly6G⁺ neutrophils peaking at 24 hours and Ly6C^high^ monocytes peaking between 24 and 72 hours, whereas Ly6C^low^ monocytes and F4/80⁺ macrophages remained relatively stable over time (**Fig. S3A–B**). Consistently, IL-6– and TNFα–expressing neutrophils and Ly6C^high^ monocytes peaked at 24 and 72 hours, respectively, while cytokine-positive Ly6C^low^ monocytes exhibited a delayed increase (**Fig. S3C–D**).

Next, we performed snRNA-seq on left ventricular tissues collected 24 h after surgery from NS-Sham, NS-I/R, and enpatoran-treated I/R mice. High-quality nuclear preparations yielded 18,133, 20,432, and 18,201 nuclei per group, respectively (**Table S6**). Uniform Manifold Approximation and Projection (UMAP) revealed distinct cardiac cell populations across all groups (**Fig. 3A**). Compared with sham controls, I/R injury resulted in a marked reduction in cardiomyocytes and a pronounced increase in infiltrating immune cells, including neutrophils and monocytes. Although the overall abundance of endothelial cells (ECs) was largely unchanged, an EC subcluster (EC8) transcriptionally proximal to infiltrating neutrophils and monocytes was significantly expanded following I/R. Enpatoran treatment substantially preserved cardiomyocyte abundance and significantly reduced neutrophil and monocyte infiltration compared with NS-treated I/R mice (**Fig. 3A, Table S7**).

**Figure 3.**
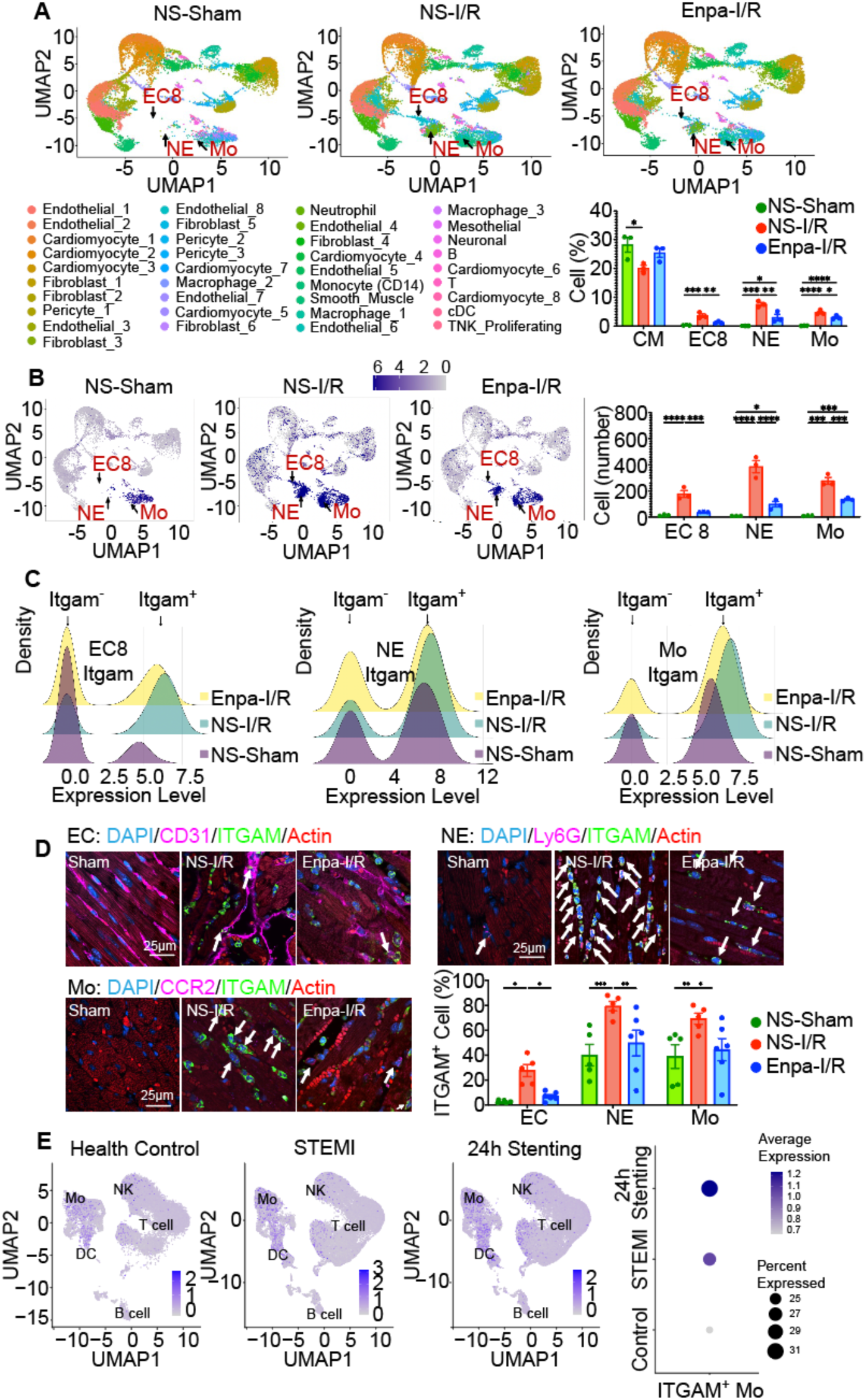
snRNA-seq reveals expansion of myeloid and endothelial cells with distinct transcriptomic signatures in ischemic MI. A, UMAP visualizing cardiac cell clusters (upper) and the percentage of CM, EC8, NE, and Mo among total cells (lower right) from three groups of male mice (n=3 per group). **B,** UMAP of Itgam^+^ cells with *Itgam* expression levels shown by a light grey-to-dark purple gradient (log2-normalized expression; *left*). Quantification of Itgam^+^ EC8, NE, and Mo was shown on the *right*. **C**, Density-Expression plot of *Itgam* in EC8 (*left*), NE (*middle*), and Mo (*right*). **D**, Representative immunofluorescent images of sarcomeric actin (*red*), CD31/Ly6G/CCR2 (*magenta*), and Itgam (*green*) of heart sections. Nuclei were stained with blue DAPI. The bar graph illustrates quantification of Itgam^+^ EC, NE, and Mo (n=5-6 mice per group). **E,** UMAP of human blood mononuclear cells showing *ITGAM* expression. scRNA-seq analysis of monocyte *ITGAM* expression from healthy controls and STEMI patients before and after coronary reperfusion. Dot plots represent the percentage of ITGAM^+^ monocytes among total mononuclear cells analyzed (n=38 patients per group). Data shown are means ± SEM. A one-way ANOVA was used to test for statistical significance. *P < 0.05, **P < 0.01, ***P < 0.001, ****P < 0.0001. NS, normal saline; Enpa, enpatoran; I/R, ischemia-reperfusion; CM, cardiomyocyte; EC, endothelial cell; NE, neutrophil; Mo, monocyte; UMAP, uniform manifold approximation and projection; DAPI, 4’, 6-diamidino-2-phenylindole; STEMI, ST-segment elevation myocardial infarction; IF, Immunofluorescence.

KEGG pathway analysis identified robust activation of innate immune and proinflammatory pathways across myeloid and endothelial compartments after I/R, including cytokine–cytokine receptor interaction, chemokine signaling, hematopoietic cell lineage, and leukocyte transendothelial migration, many of which were attenuated by enpatoran treatment (**Table S8**). Similar to infiltrating immune cells, multiple endothelial clusters (EC4, EC5, and EC8) exhibited upregulation of proinflammatory immune-related pathways following I/R. Notably, the leukocyte transendothelial migration pathway was significantly induced in the EC clusters after I/R and was substantially attenuated in response to enpatoran treatment (EC4, *P* < 0.001; EC5, *P* = 0.210; EC8, *P* < 0.001). Moreover, EC4 was characterized by enrichment of pathways associated with endothelial activation, cytoskeletal remodeling, cell–cell and cell–matrix interactions, and leukocyte trafficking, consistent with a proinflammatory, barrier-regulatory endothelial phenotype. Many inflammation-related pathways in EC4 were downregulated following enpatoran treatment. Similarly, EC5 displayed upregulation of inflammatory and immune signaling, endothelial structural remodeling, and barrier regulation, most of which were also attenuated by enpatoran. Collectively, these findings demonstrate that myocardial I/R induces a coordinated expansion of myeloid cells and robust transcriptional reprogramming of immune cells and endothelial cells, which are significantly reversed by TLR7 inhibition.

Importantly, snRNA-seq analysis also revealed distinct cell–type–specific expression patterns of Tlr (Tlr1-13) family genes in the heart. Under Sham conditions, *Tlr7* emerged as one of the most highly expressed TLRs in the heart, second only to Tlr4 (**Fig. S4A**). While *Tlr4* expression was predominantly detected in cardiomyocytes, *Tlr7* expression was enriched mainly in CD14⁺ monocytes, macrophage subsets, and EC8, with minimal expression in neutrophils. Following I/R injury, overall cardiac expression of *Tlr4 and Tlr7* increased; however, their cell–type–specific expression patterns differed. *Tlr4* upregulation was most prominent in cardiomyocytes, monocytes, and neutrophils. In contrast, *Tlr7* expression was selectively increased in proliferating T/NK cells and macrophage subsets, while its expression decreased in EC8 and CD14⁺ monocytes (**Fig. S4B–C**). Pharmacologic TLR7 inhibition with enpatoran resulted in distinct modulatory effects on *Tlr4* and *Tlr7* expression across cardiac cell populations. Enpatoran treatment markedly attenuated *Tlr4* expression in monocytes and neutrophils, whereas *Tlr7* expression was significantly reduced in proliferating T/NK cells and macrophage subsets, accompanied by a relative restoration of *Tlr7* expression in EC8 endothelial cells and CD14⁺ monocytes (**Fig. S4B–C**).

### *Itgam* expression is upregulated in cardiac cells and in circulating monocytes after I/R

Among many genes associated with these proinflammatory pathways, *Itgam* exhibited an impressive upregulation following I/R injury and downregulation to enpatoran treatment in monocytes (I/R vs. Sham: 4.0-fold; Enpa-I/R vs. NS-I/R: 0.59-fold), neutrophils (Enpa-I/R vs. NS-I/R: 0.59-fold), and EC8 (I/R vs. Sham: 16-fold; Enpa-I/R vs. NS-I/R: 0.37-fold). ITGAM is a critical subunit of the Mac-1 integrin complex (CD11b/CD18, or known as Itgam/Itgb2) predominantly expressed in neutrophils and monocytes but also in endothelial cells(*29–31*) and involved in various immune functions, including leukocyte adhesion, migration, phagocytosis, and several immunological disorders(*32*), mainly through the binding of the I-domain of ITGAM to ligands like ICAM-1(*33*). The abundance of Itgam^+^ cells was significantly increased in the EC8, monocyte, and neutrophil clusters (**Fig. 3B**). Density-Expression plot further revealed an increased proportion of Itgam^+^ cells within these cell clusters, accompanied by elevated overall *Itgam* expression in the I/R group (**Fig. 3C**). These transcriptional changes were corroborated at the protein level by immunofluorescence analysis, which revealed increased *Itgam* expression in myocardial CD31⁺ endothelial cells, Ly6G⁺ neutrophils, and CCR2⁺ monocytes following I/R injury (**Fig. 3D and Fig. S5–S7**). In sham-operated hearts, endothelial cells exhibited an intact, well-organized vascular architecture with minimal immune cell infiltration or *Itgam* expression. In contrast, I/R injury disrupted endothelial organization and myocardial structure, with markedly increased ITGAM⁺ endothelial cells and enhanced infiltration of ITGAM⁺ neutrophils and monocytes. Enpatoran treatment reversed these effects, significantly reducing ITGAM expression in cardiac endothelial cells, neutrophils, and monocytes (**Fig. 3B–D**).

To determine whether cardiac I/R induces systemic leukocyte expansion and ITGAM expression, we analyzed leukocytes from peripheral blood, spleen, and bone marrow. Circulating neutrophil numbers increased rapidly within 2 hours after I/R, while monocyte numbers modestly decreased at 4 hours. (**Fig. S8A-B3**). Importantly, ITGAM expression in circulating monocytes was significantly increased between 2-4 hours after I/R, which was reversed by enpatoran treatment (**Fig. S8B**). No significant alterations were observed in the spleen or bone marrow, with the exception of a modest increase in splenic neutrophils and a concomitant decrease in splenic monocytes after I/R (**Fig. S9**).

To assess *ITGAM* expression in human leukocytes, we re-analyzed a previously published single-cell RNA-seq dataset from patients with acute MI(*34*). The proportion of monocytes within peripheral blood mononuclear cells (PBMCs) was significantly increased in patients with STEMI and further expanded 24 hours after stent placement compared with healthy controls. As anticipated, TLR8 was highly expressed compared to *TLR7* in human monocytes; both receptors share functional similarities in endosomal RNA sensing and innate immune activation. Importantly, like mouse blood monocytes, monocytes from STEMI patients were increased in % *ITGAM+* numbers as well as level of *ITGAM* expression, which was further increased after coronary stenting (**Fig. 3E**). Together, these results indicate that myocardial I/R induces TLR7-dependent ITGAM upregulation across cardiac endothelial cells, neutrophils, and monocytes, as well as in circulating monocytes

### TLR7 activation promotes *Itgam* expression and endothelial–immune cell crosstalk

To determine whether TLR7 activation is sufficient to induce *Itgam* expression, bone marrow–derived macrophages (BMDMs) and human coronary artery endothelial cells (HCAECs) were treated with the TLR7/8 agonists R837, R848, or miR-146a-5p, an inflammatory miRNA that binds to and activates TLR7(*20*) (**Fig. 4A**). R837 increased *Itgam* expression in BMDMs at 6 and 18 hours, accompanied by an increase in IL-6 production (**Fig. 4B–C**). Confocal immunofluorescent microscopy demonstrated marked increases in ITGAM protein expression in both BMDMs and HCAECs following TLR7/8 stimulation (**Fig. 4D–G**). To assess the functional consequences of TLR7/8 activation, we used a microfluidic adhesion assay to quantify human EC (HCAECs)–monocyte (THP-1) adhesion under physiological shear stress. R848 treatment of either HCAECs or THP-1 monocytes increased endothelial–monocyte adhesion to a similar extent, whereas concurrent stimulation of both cell types resulted in a significantly greater increase in monocyte adhesion (**Fig. S10**) (**videos S1–S2**). Notably, R848-induced endothelial–monocyte adhesion was markedly attenuated by enpatoran treatment, confirming a TLR7/8-mediated process (**Fig. 5A–B**). To determine the contribution of ITGAM to R848-induced endothelial–monocyte crosstalk, an ITGAM-neutralizing antibody was applied to either HCAECs or THP-1 monocytes after R848 stimulation. Blocking ITGAM on THP-1 monocytes reduced adhesion to endothelial cells by 85.8% (**videos S3**), whereas ITGAM blockade on HCAECs produced a more modest reduction of 51.9% (**Fig. 5C–D, videos S4**). Concurrent ITGAM blockade in both cell types resulted in the greatest reduction in endothelial–monocyte adhesion (**videos S5**), suggesting that both EC and monocyte ITGAM play significant roles, but monocyte-expressed ITGAM plays a more dominant role, in mediating TLR7-induced monocyte–endothelial interactions. Similar results were observed in human neutrophil–EC interaction (**Fig. 5E–F)**. Together, these results suggest that TLR7 activation enhances endothelial–Mo/NE functional crosstalk through an ITGAM-dependent mechanism.

**Figure 4.**
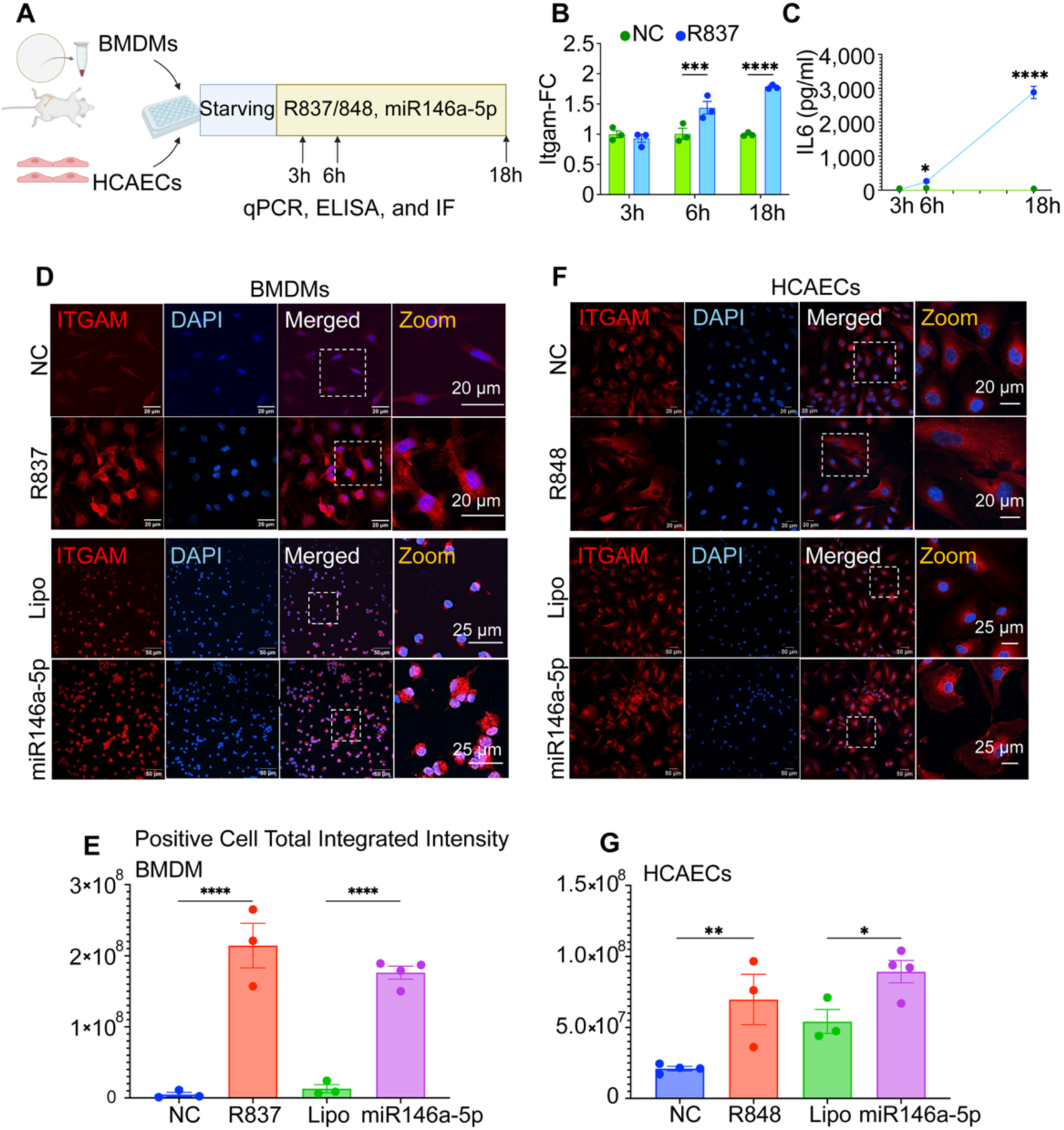
TLR7 activation increases *Itgam* expression in macrophages and endothelial cells. A, Experimental diagram of cell treatment with TLR7 agonists. **B,** Expression level of *Itgam* in BMDMs was examined by qRT-PCR. Relative expression of *Itgam* in the R837-treatment group was calculated with the non-treated control (NC). **C**, IL-6 production by BMDMs as measured by ELISA. **D & F**, The representative images of ITGAM expression in BMDMs (**D**) and HCAECs (**F**). **E & G,** Quantification of the ITGAM^+^ cell total integrated intensity. Data shown are means ± SEM. A one-way ANOVA was used to test for statistical significance. *P < 0.05, **P < 0.01, ***P < 0.001, ****P < 0.0001. NC, nontreatment control; DAPI, 4′,6-diamidino-2-phenylindole; BMDMs, bone-marrow-derived macrophage; HCAEC, human coronary artery endothelial cell; FC, fold-change; lipo, lipofectamine; integrated intensity, the sum of pixel intensities within a designated area of a cell.

**Figure 5.**
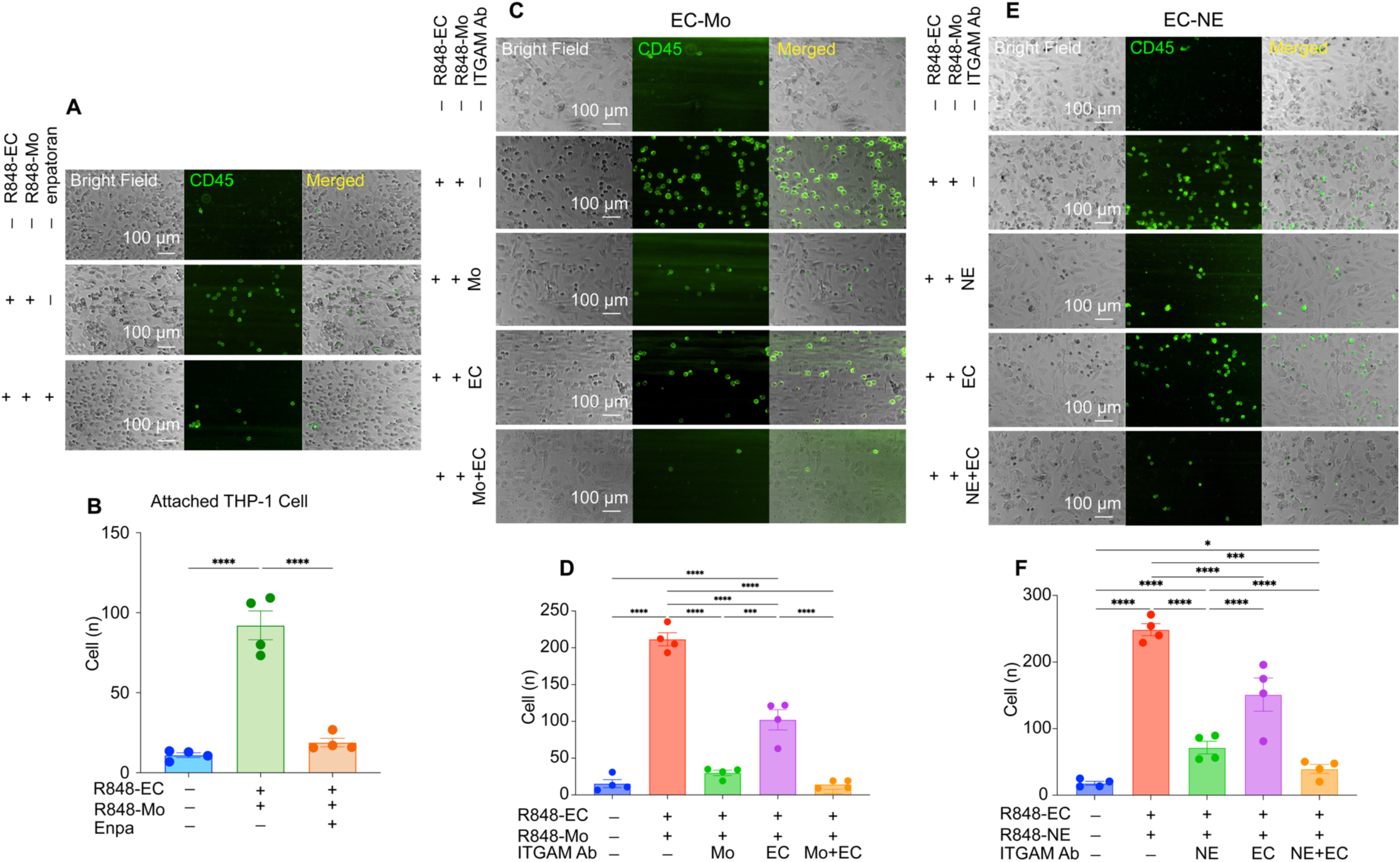
TLR7 activation increases leukocyte-endothelial adhesion. A, C, **E**, Microfluidic adhesion assay of THP-1 (Mo) or human fresh neutrophil adhesion to HCAEC under different treatment conditions as indicated (R848/enpatoran: 3h for Mo, 18h for EC; ITGAM Ab: 1h). **B, D, & F**, Quantification of THP-1 or human fresh neutrophil adhesion under different treatment conditions. CD45⁺ immune cells are shown in green. Adherent CD45⁺ cells appear as discrete green puncta, while continuous green streaks reflect perfused monocytes under flow conditions. Data shown are means ± SEM. A one-way ANOVA was used to test for statistical significance. *P < 0.05, ***P < 0.001, ****P < 0.0001. HCAEC, Human coronary artery endothelial cell; NE, neutrophil; Mo, monocyte; ITGAM Ab, ITGAM neutralizing antibody.

### ITGAM neutralization attenuates cardiac leukocyte infiltration and confers protection against myocardial I/R injury

To determine whether ITGAM contributes to post-ischemia inflammation and injury after I/R, mice were treated with an ITGAM-neutralizing antibody (ITGAM Ab) or an isotype control antibody (100 μg per mouse, i.p.) 1 hour before ischemia (**Fig. 6A**). Mice receiving ITGAM Ab exhibited significantly smaller infarct as evidenced by reduced MI/LV and MI/AAR ratios (**Fig. 6B–C**) and improved systolic (better LVEF, CO, and GLS) and diastolic (smaller LA dilation and a lower E/e′ sr ratio) functions as compared with the controls (**Fig. 6D–H, and Table S9).** ITGAM Ab treatment also markedly reduced infiltration of neutrophils and monocytes after I/R injury (**Fig. 6I–L**). These results support the notion that ITGAM expression, driven by I/R and mediated by TLR7 signaling, plays an important role in post-ischemia myocardial inflammation and injury.

**Figure 6.**
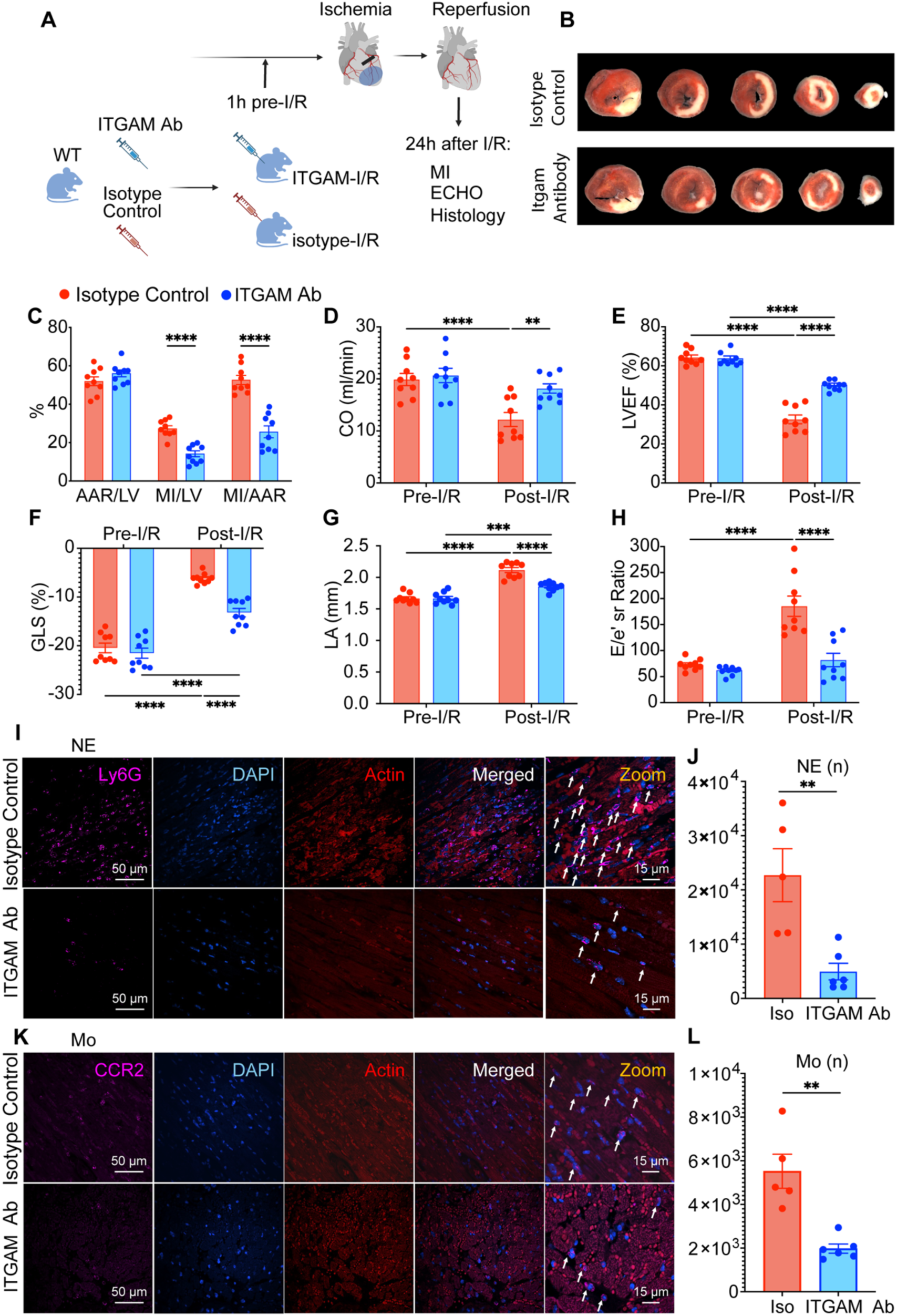
ITGAM neutralization attenuates leukocyte infiltration and limits ischemic MI. **A**, Study diagram for intraperitoneal administration of ITGAM neutralizing antibody or control isotype in a mouse model of MI and various experimental endpoints (n = 9 mice per group). **B**, Representative images of MI areas after triphenyl tetrazolium chloride (TTC) staining. **C**, Quantification of AAR/LV, MI/LV, and MI/AAR 24 hours after I/R. **D-E**, Cardiac output (CO) (**D**) and left ventricular ejection fraction (LVEF) (**E**). **F**, Echocardiography speckle tracking imaging showing global longitudinal strain (GLS). **G-H**, Echocardiographic data depict LA diameter (**G**), and E wave to reversed longitudinal strain rate ratio (E/e’ sr) (**H**). **I and K**, Immunofluorescence staining of heart sections to show sarcomeric actin (red), and Ly6G/CCR2 (magenta). DAPI was stained in blue. **J & L**, Quantification of the cardiac infiltrated NE and Mo after I/R injury. Data shown are means ± SEM. Two-way ANOVA was used to test statistical significance. **P < 0.01, ***P < 0.001, ****P < 0.0001. I/R, ischemia-reperfusion; Enpa, enpatoran; ECHO, echocardiography; CO, cardiac output; LVEF, left ventricular ejection fraction; GRS, global radial strain; LA, left atrium; E/e’ sr, mitral valve E wave velocity and reversed longitudinal strain rate ratio; NE, neutrophil; Mo, monocyte.

### Enhanced ICAM1-ITGAM interaction after I/R injury

Leukocyte adhesion to the endothelium is a critical driver of myocardial inflammation following I/R injury(*35*). To delineate the molecular mechanisms underlying this endothelial–leukocyte crosstalk in vivo, we performed *in silico* ligand–receptor (L–R) interaction analyses using our cardiac snRNA-seq dataset and quantified expressed L–R pairs among major cardiac cell types. Among 2,377 identified L–R pairs, 24 were significantly upregulated after I/R and reversed by enpatoran treatment specifically in endothelial cell–monocyte (EC–Mo) or endothelial cell–neutrophil (EC–NE) interactions (**Fig. 7A**). Most of these L-R pairs are functionally linked with inflammatory processes, including leukocyte adhesion (ICAM1–ITGAM), chemotactic recruitment (CXCL2–CXCR2), and immune–vascular signaling (SEMA7A–[ITGB1+ITGA1]). Notably, the ICAM1–ITGAM emerged as a prominent pair, as it was upregulated after I/R and attenuated by enpatoran treatment in both EC–Mo and EC–NE interactions. Given the central role of leukocyte–endothelial adhesion in post-ischemic inflammation, we further examined regulation of the ICAM1–ITGAM axis across relevant cardiac cell populations. Unlike *Itgam*, which was upregulated in neutrophils, monocytes, and endothelial cells after I/R and attenuated by enpatoran treatment (**Fig. 3B–D, Fig. S11A**), *Icam1* expression was increased in endothelial cells and expressed predominantly in EC4 and EC5 when compared with monocytes and neutrophils after I/R and was not significantly affected by enpatoran treatment (**Fig. S11B–C**).

**Figure 7.**
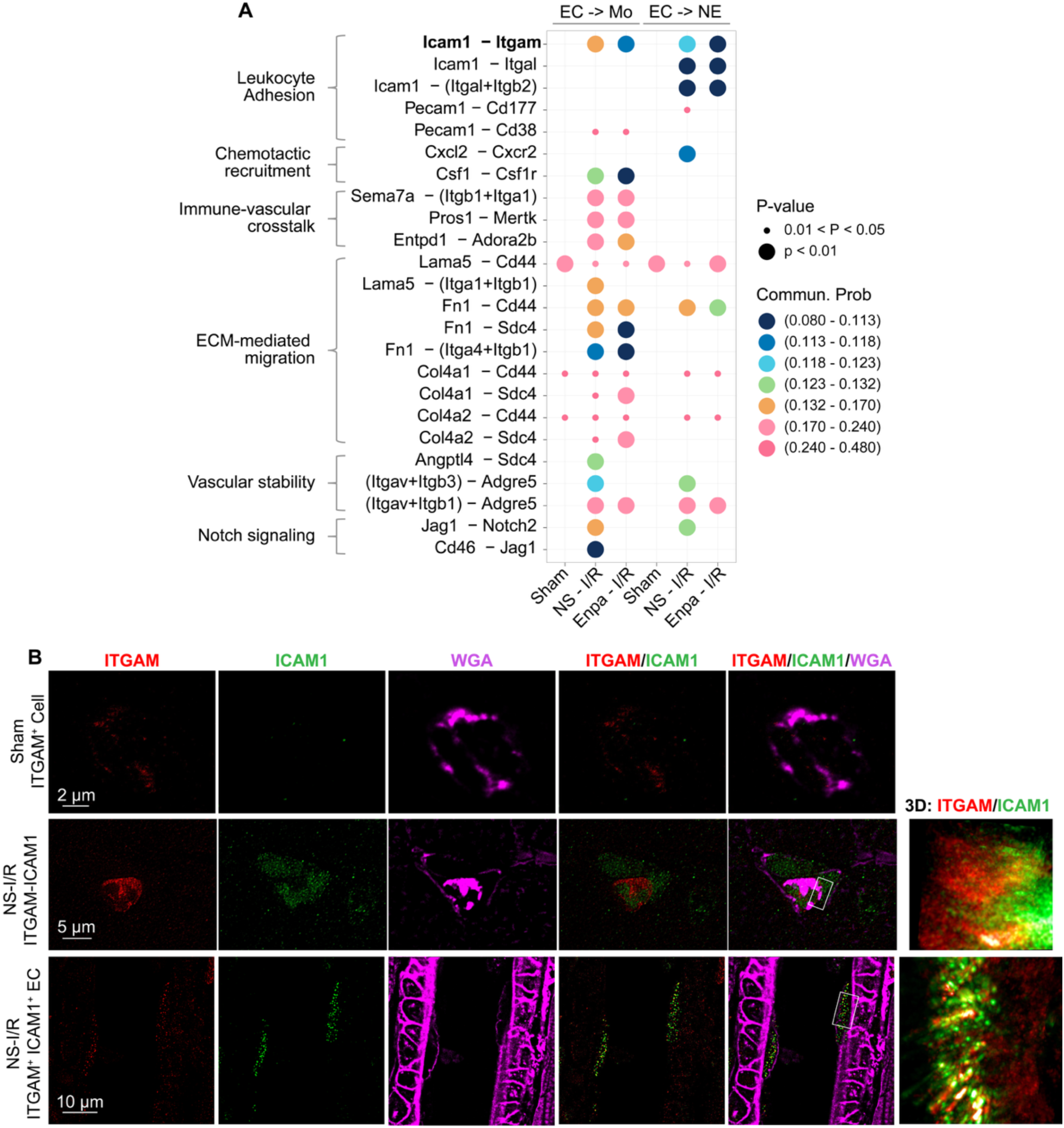
Myocardial I/R enhances ICAM1–ITGAM interaction. **A.** *In silico* ligand–receptor (L–R) interaction analysis of the snRNA-seq dataset. The 24 listed L–R pairs are associated with inflammatory and immune-related functions. Communication probabilities via L-R pair interaction between endothelial cells (ECs) and neutrophils (NEs) or monocytes (Mos) across sham, NS-I/R, and Enpa-I/R groups are shown. Each dot represents an individual L–R pair, with color indicating communication probability and dot size reflecting statistical significance (*P* value). Key endothelial-to-immune interactions, **Icam1–Itgam**, are bolded, revealing the dynamic changes in communication in response to I/R and to enpatoran treatment. **B,** STED super-resolution immunofluorescent images of heart sections to show ITGAM (red), ICAM1 (green), and cell membranes (WGA, magenta). ICAM1–ITGAM colocalization was observed within endothelial junctional membrane protrusions in I/R mice. I/R, ischemia-reperfusion; Mac-1, Itgam + Itgb2; NS, normal saline; Enpa, enpatoran; EC, Endothelial cell; Mo, monocyte; NE, neutrophil; WGA, Wheat Germ Agglutinin.

To determine whether these transcriptional changes translated into physical molecular interactions, we performed STED super-resolution immunofluorescence imaging on myocardial sections. Following I/R injury, ITGAM⁺ leukocytes were frequently observed in close apposition to ICAM1⁺ endothelial cells, with discrete ICAM1–ITGAM interaction sites at cell-to-cell interfaces (**Fig. 7B, middle panel, Video S6**). In addition, colocalization and close proximity of ICAM1 and ITGAM were detected within endothelial junctional membrane protrusions, structures that have been reported to serve as hotspots for neutrophil transmigration(*36*) (**Fig. 7B, lower panel, Videos S7**). In contrast, sham hearts contained a small number of ITGAM-expressing leukocytes without detectable ITGAM–ICAM1 leukocyte–endothelial contacts (**Fig. 7B, Fig. S12, Video S8**). Together, these findings suggest that myocardial I/R enhances ICAM1–ITGAM interaction and endothelial**–**leukocyte crosstalk, a process regulated by TLR7 signaling.

### TLR7 regulates ICAM1 expression and leukocyte-endothelium adhesion

To determine whether TLR7/8 activation is sufficient to induce ICAM1 expression, BMDMs and HCAECs were treated with the TLR7/8 agonists R837 or R848, or with miR-146a-5p. Confocal immunofluorescence microscopy revealed a significant increase in ICAM1 expression as quantified as total integrated fluorescence intensity in BMDMs and HCAECs treated with R837, R848, or miR-146a-5p compared with untreated controls (**Fig. 8A–B**). To assess the functional contribution of ICAM1 to leukocyte–endothelial interactions, an ICAM1-neutralizing antibody was applied to either THP-1 monocytes or HCAECs following R848 stimulation. ICAM1 blockade on HCAECs markedly reduced THP-1 adhesion (**Videos S9**), whereas ICAM1 blockade on THP-1 cells resulted in a very modest reduction (**Fig. 8C–D, Videos S10**). Simultaneous ICAM1 blockade in both cell types produced the greatest inhibition of monocyte adhesion, indicating that endothelial ICAM1 plays a dominant role in mediating TLR7-induced monocyte–endothelial interactions (**Videos S11**). A similar inhibitory pattern was observed in human neutrophil adhesion assays (**Fig. 8E–F**). Collectively, these findings demonstrate that TLR7 signaling is sufficient to induce ICAM1 expression and acts upstream of coordinated ITGAM and ICAM1 induction, thereby promoting enhanced leukocyte–endothelial crosstalk.

**Figure 8.**
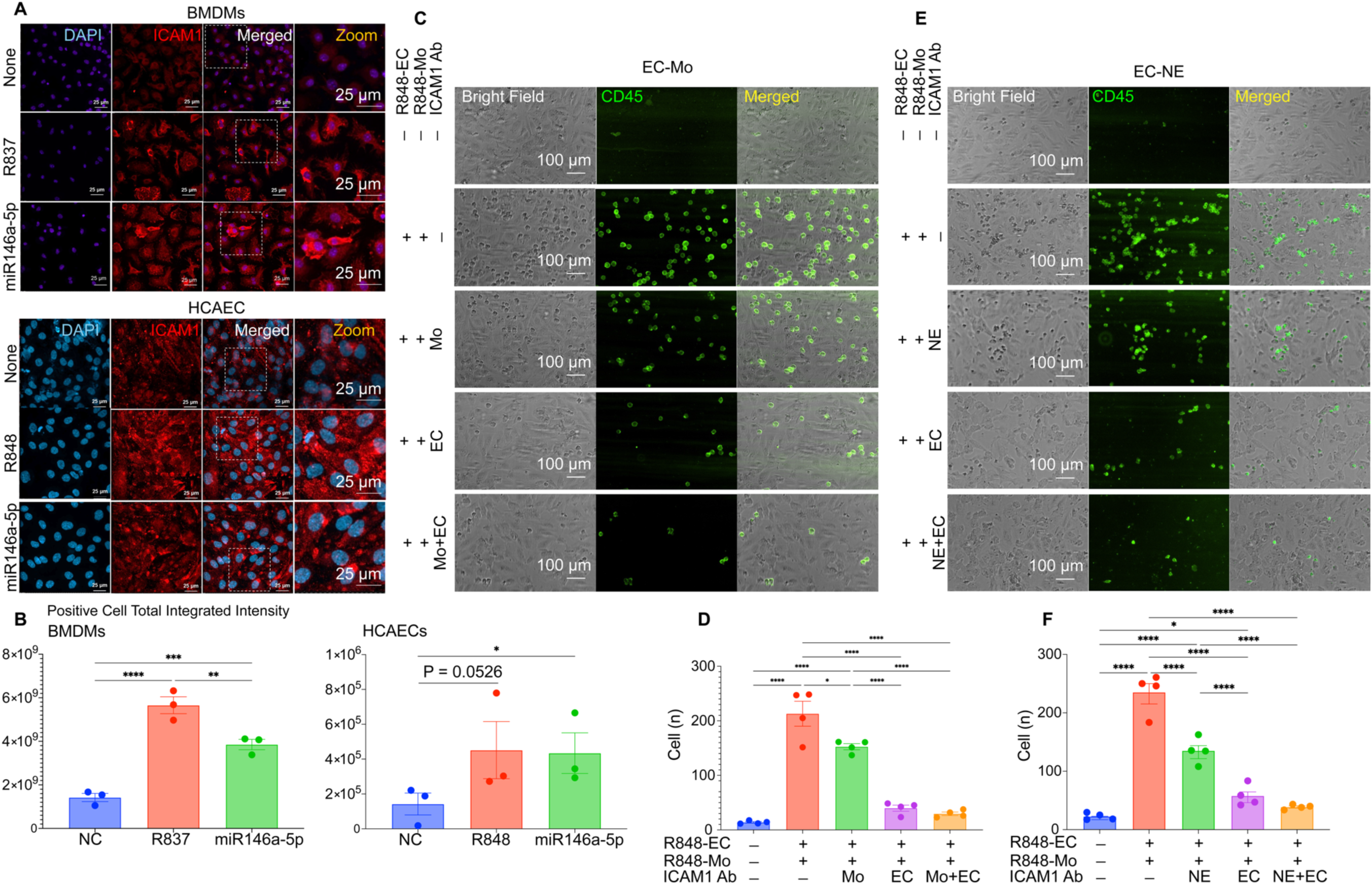
TLR7 activation promotes ICAM1 expression and leukocyte–endothelial interaction. A, The representative images of ICAM1 expression in BMDMs and HCAECs. **B,** Quantification of the ICAM1^+^ cell total integrated intensity. **C-D,** Microfluidic adhesion assay of THP-1 (Mo) adhesion to HCAEC (**C**) and quantification (**D**) under different treatment conditions. **E-F,** Microfluidic adhesion assay of human fresh neutrophil adhesion to HCAEC (**E**) and quantification (**F**) under different treatment conditions. Treatment time for R848/enpatoran: 3h for Mo, 18h for EC; ICAM-1 Ab: 1h. Adherent CD45⁺ cells appear as discrete green puncta, while continuous green streaks reflect perfused monocytes under flow conditions. Data shown are means ± SEM. A one-way ANOVA was used to test for statistical significance. *P < 0.05, **P < 0.01, ***P < 0.001, ****P < 0.0001. NC, nontreatment control; DAPI, 4′,6-diamidino-2-phenylindole; BMDMs, bone-marrow-derived macrophages; HCAECs, human coronary artery endothelial cells; Integrated intensity, the sum of pixel intensities within a designated area of a cell; ICAM1 Ab, ICAM1 neutralizing antibody; Mo, monocyte; NE, neutrophil.

## DISCUSSION

In this study, we uncover a previously unrecognized role for Toll-like receptor 7 (TLR7) as a central innate immune sensor that links ischemia–reperfusion (I/R) injury to pathological leukocyte–endothelial interactions in the heart. Using single-nucleus transcriptomics, we identify coordinated expansion of cardiac myeloid cells and distinct endothelial cell (EC) populations enriched for inflammatory and leukocyte adhesion programs following I/R. Among these shared responses, *Itgam*, encoding the leukocyte adhesion receptor CD11b, emerged as a convergent and conserved signature, increased not only in cardiac immune and endothelial compartments after I/R but also in circulating monocytes from mice and from patients with STEMI 24 hours after coronary stenting. Functionally, pharmacologic inhibition of TLR7 with the selective mTLR7/hTLR8 antagonist enpatoran reduced infarct size and improved cardiac function across a clinically relevant therapeutic window encompassing pre-, peri-, and early post-ischemia administration. Mechanistically, TLR7 activation drove leukocyte–endothelial adhesion by promoting coordinated induction of ITGAM and ICAM1, whereas TLR7 inhibition or ITGAM neutralization disrupted this interaction, attenuated myocardial inflammation, and limited post-ischemia MI. Together, these findings position TLR7 as a druggable upstream regulator of leukocyte–endothelial adhesion and suggest that targeting innate immune RNA sensing may represent an effective strategy to limit cardiac ischemic injury during reperfusion therapy.

Previous studies have shown that myocardial I/R promotes the release of cellular RNA and miRNAs into the extracellular space and that removal of extracellular RNA (ex-RNA) confers cardioprotection, implicating ex-RNA as a mediator of myocardial injury (*9, 11, 13, 14*). Consistent with these observations, plasma bulk RNA-seq in the present study revealed increased circulating miRNAs, several of which exerted potent proinflammatory effects through TLR7-dependent signaling. In parallel, snRNA-seq analysis of the myocardium identified *Tlr7* as one of the most highly expressed Toll-like receptors in the heart at baseline, second only to *Tlr4*. Following I/R injury, *Tlr7* expression was increased, with particularly pronounced upregulation observed in inflammation-associated cell populations, including proliferating TNK cells and macrophage subsets. Notably, CD14⁺ monocytes exhibited the highest baseline *Tlr7* expression among all cardiac cell types under Sham conditions, yet *Tlr7* expression in this population was markedly reduced after I/R, despite the overall inflammatory milieu. Interestingly, enpatoran treatment partially restored *Tlr7* expression in monocytes, suggesting a dynamic and context-dependent regulation of TLR7 signaling. The biological significance and mechanism of TLR regulation in I/R are unclear. One possible explanation is that acute myocardial I/R triggers TLR7 activation in monocytes, which leads to receptor internalization and/or degradation, a regulatory mechanism described for endosomal TLRs to limit excessive innate immune activation. A similar TLR7-mediated downregulation has been reported for IRAK-1, a signaling kinase of TLR4 and TLR7, in macrophages stimulated by TLR7 agonist R837 and miR-146a-5p(*20*). Pharmacologic inhibition with enpatoran may attenuate this activity-dependent TLR7 downregulation, thereby preserving or restoring detectable *Tlr7* expression. However, the precise mechanisms governing cell–type–specific TLR regulation following I/R and in response to TLR7 inhibition warrant further investigation.

Enpatoran (M5049), developed by Merck KGaA, is a first-in-class, selective inhibitor of murine TLR7 and human TLR8 that can be administered orally or intravenously (*28*). Enpatoran inhibits signaling induced by both synthetic and endogenous RNA ligands and has been investigated as a therapeutic candidate for autoimmune and inflammatory diseases characterized by aberrant TLR7/8 activation, including systemic and cutaneous lupus erythematosus (SLE and CLE) and myositis(*37, 38*). Early-phase clinical studies show that enpatoran is safe and well-tolerated in healthy volunteers and in patients with SLE and CLE (*39–41*). Subsequent phase 2 trials showed a significant, dose-dependent reduction in cutaneous disease activity in patients with CLE or mild SLE(*37, 42*). In patients with COVID-19, enpatoran treatment was associated with a trend toward reduced clinical deterioration, disease progression events, and all-cause mortality compared with placebo(*43*). The data presented here extend the therapeutic potential of enpatoran to cardiovascular disease, demonstrating a previously unrecognized application in preventing and treating cardiac I/R injury during reperfusion therapy.

To model clinically relevant reperfusion scenarios after a heart attack, we evaluated multiple dosing regimens spanning pre-ischemia, peri-ischemia, and early post-ischemia administration. Enpatoran significantly reduced infarct size and preserved left ventricular function when administered before ischemia, during ischemia, or shortly after reperfusion; this benefit was absent in TLR7-deficient mice, confirming TLR7-dependent efficacy, not an off-target benefit. In contrast, protection was lost when treatment was delayed until 2 hours after reperfusion, underscoring a narrow therapeutic window for TLR7-mediated I/R injury. These findings are consistent with prior reports linking TLR7 to other cardiac conditions such as myocarditis(*26*), conduction abnormalities(*27*), and adverse post–MI remodeling(*25*).

To determine an appropriate time window for mechanistic interrogation, we performed a pilot time-course analysis of cardiac inflammatory cell dynamics from 1 to 14 days after I/R or sham surgery. The data confirmed that neutrophils and proinflammatory monocytes peaked between 24 and 72 hours after I/R. snRNA-seq at 24 hours demonstrated a marked shift in cardiac cellular composition, characterized by a reduction in cardiomyocytes and robust expansion of immune cell populations, particularly neutrophils and monocytes, consistent with prior reports (*44–47*). In contrast, the overall endothelial cell (EC) population was largely preserved. A key observation, however, was the selective expansion of a distinct endothelial subpopulation, EC8, which increased from ∼0.3% of total cardiac cells at baseline to ∼3.6% after I/R. EC8 localized in close proximity to neutrophils and monocytes on UMAP projections, demonstrating a pronounced proinflammatory and immune-interactive program that closely mirrored infiltrating innate immune cells, suggesting coordinated endothelial–immune reprogramming during post-ischemic inflammation. Importantly, both immune cell infiltration and EC8 expansion were markedly attenuated by enpatoran treatment, thus linking TLR7 to these downstream events. Together, these findings identify a TLR7-dependent inflammatory endothelial state that emerges during I/R and provide a cellular framework connecting innate immune RNA sensing to immune–endothelial crosstalk in myocardial I/R injury.

Moreover, transcripts associated with leukocyte transendothelial migration pathway were selectively upregulated in EC8 after I/R and attenuated by enpatoran treatment. Among these genes, *Itgam* emerged as a particularly compelling candidate, as its expression was robustly induced by I/R and consistently suppressed by TLR7 inhibition across myocardial EC8, neutrophils, and monocytes. *Itgam* encodes the αM subunit of the Mac-1 integrin complex, a key regulator of immune cell adhesion, migration, and phagocytosis (*48–52*), largely through binding of its I-domain to ligands such as ICAM1(*33*). The shared, TLR7-dependent induction of *Itgam* across endothelial and myeloid compartments therefore supports ITGAM as a convergent molecular effector linking innate immune sensing to immune–endothelial interactions during I/R. Beyond the myocardium, myocardial I/R was associated with an increased proportion of Itgam-expressing circulating monocytes in mice and in patients with STEMI, both prior to and 24 hours after coronary stenting, suggesting that the TLR7-dependent upregulation of Itgam identified in cardiac cells is also reflected in peripheral blood monocytes. Although the exact functional significance of this finding in circulating monocytes remains unclear, one possibility is that elevated cellular Itgam expression facilitates the recruitment of circulating monocytes to the heart during reperfusion.

A key finding of the current study is that TLR7 emerged as a critical upstream regulator of leukocyte–endothelial adhesion through coordinated control of the ICAM1–ITGAM communication. *In silico* L-R pair analyses based on snRNA-seq of the heart uncovered dynamic change in *Icam1–Itgam* communication between EC, mainly EC4 and EC5 subpopulations, and leukocyte after I/R and in response to enpatoran treatment, which was supported by super-resolution STED imaging at cell-cell interface and within endothelial junctional membrane protrusions in I/R heart. *In vitro*, TLR7 activation was sufficient to induce robust ITGAM and ICAM1 expression in immune and endothelial cells and to enhance leukocyte–endothelial adhesion under physiological shear stress, whereas neutralizing antibody studies identified leukocyte-expressed ITGAM and EC-associated ICAM1 as the dominant determinant of leukocyte-endothelial interaction. These findings were further supported in vivo, where ITGAM neutralization significantly reduced neutrophil and monocyte infiltration and limited post-ischemia MI size, although the relative contributions of neutrophil-, monocyte-, and endothelial**–**specific ITGAM are unclear and will require future investigation.

Several limitations of this study warrant consideration. *In vivo* experiments were conducted in young, healthy mice, which do not fully reflect the clinical complexity of myocardial infarction, where aging and cardiometabolic comorbidities may influence innate immune signaling and therapeutic responsiveness. Since both pharmacologic TLR7 inhibition and antibody-mediated ITGAM neutralization modulate the global TLR7-ITGAM pathway, the relative contributions of endothelial-, neutrophil-, and monocyte-derived TLR7–ITGAM axis to leukocyte–endothelial adhesion and I/R injury remain unresolved and will require further investigation using cell–type–specific genetic approaches.

In conclusion, our findings identify TLR7 as a central innate immune RNA sensor that mechanistically links I/R–induced immune activation to endothelial dysfunction and myocardial injury. By coordinating ICAM1–ITGAM expression and engagement, TLR7 signaling drives leukocyte recruitment and sustains post-ischemic inflammatory responses. Pharmacologic inhibition of TLR7 disrupted immune–endothelial interactions, limited infarct expansion, and preserved cardiac function within a defined therapeutic window. Together, these data position the TLR7–ITGAM axis as a pivotal regulator of endothelial–leukocyte crosstalk in myocardial I/R injury and highlight TLR7 signaling as a promising target for therapeutic intervention in acute MI.

## MATERIALS AND METHODS

### Study Design

We hypothesized that Toll-like receptor 7 (TLR7), a sensor of single-stranded RNA, mediates myocardial I/R injury. To test this, we first performed bulk plasma RNA sequencing to identify miRNAs upregulated after myocardial I/R and assessed their ability to induce MIP-2 production in a TLR7-dependent manner. We then evaluated the therapeutic efficacy of pharmacologic TLR7 inhibition in a well-established murine model of cardiac I/R injury. The selective TLR7/8 antagonist enpatoran was administered before, during, or after ischemia to model clinically relevant timing relative to coronary reperfusion therapy. Myocardial infarct size and cardiac function were assessed by histochemical staining and echocardiography to define the temporal window during which TLR7 blockade attenuates reperfusion injury.

To elucidate downstream mechanisms regulated by TLR7 signaling during reperfusion, we performed snRNA-seq on cardiac tissue from WT mice subjected to sham surgery or I/R with or without enpatoran treatment. This unbiased approach enabled systematic identification of cell compositions and cell–type–specific transcriptional programs associated with reperfusion injury. To establish translational relevance, we analyzed publicly available single-cell transcriptomic datasets of blood monocytes from patients with STEMI following coronary stenting, focusing on the expression of *Itgam*, *Tlr7*, and *Tlr8*.

We next performed mechanistic and functional validation studies to define how TLR7 activation regulates leukocyte–endothelial crosstalk. Using *in vitro* systems, including BMDMs. THP-1 monocytes, and HCAECs, we tested whether selective TLR7 activation was sufficient to induce adhesion molecule expression. Expression of ITGAM and ICAM1 was quantified using ImageXpress automated cell imaging. Functional consequences of TLR7 activation were assessed using a microfluidic adhesion assay under physiological shear stress, in which monocytes or neutrophils were perfused over endothelial monolayers with or without pharmacologic TLR7 inhibition. To delineate cell–type–specific contributions, neutralizing antibodies targeting ITGAM or ICAM1 were applied selectively to leukocytes or ECs to evaluate their relative roles in mediating adhesion.

Guided by snRNA-seq findings, we further performed *in silico* ligand–receptor interaction analyses using CellChat to identify adhesion-related interactions among immune and endothelial cell populations, with particular focus on the ICAM1–ITGAM pair. These predicted interactions were subsequently validated at the molecular level using super-resolution stimulated emission depletion (STED) microscopy to directly visualize ICAM1–ITGAM interactions at leukocyte–endothelial interfaces and within cells on sliced myocardium.

For all *in vivo* studies, mice were randomly assigned to treatment groups, and investigators performing surgeries, administering treatments, and analyzing data were blinded to genotype and treatment allocation. Sample sizes were determined by power analysis to achieve 80% power (two-sided *P* < 0.05) with a minimum detectable effect size comparable to prior I/R studies. All animal experiments were conducted in accordance with protocols approved by the Institutional Animal Care and Use Committee of the University of Maryland School of Medicine.

### Statistical analysis

Statistical analyses were performed using GraphPad Prism (v10.4.0; GraphPad Software, La Jolla, CA), R (v4.3.0), and relevant bioinformatics packages as specified below. Continuous variables are presented as mean ± SEM unless otherwise indicated. Normality was assessed where appropriate. For comparisons between two groups, Student’s t test or the Mann–Whitney U test was used as appropriate. For comparisons involving more than two groups, one-way ANOVA followed by Tukey’s post hoc test was applied. For experiments involving multiple treatment protocols and interventions, two-way ANOVA was used to assess the effects of treatment and protocol, followed by appropriate post hoc testing.

### RNA sequencing data analysis

snRNA-seq data underwent quality control and computation of gene-barcode matrices using the Cell Ranger pipeline (10x Genomics) with default parameters. Gene-barcode matrices were imported into Seurat (v4.3.0) and underwent integration, batch effect correction, and normalization for downstream analysis. Low-quality nuclei were filtered based on read depth, gene complexity, and mitochondrial content to remove potential doublets or damaged nuclei. Raw UMI counts were normalized using Seurat’s SCTransform workflow, and highly variable genes were identified for downstream analyses. Dimensionality reduction was performed using principal component analysis (PCA), followed by Uniform Manifold Approximation and Projection (UMAP) was used for clustering of individual cells. Cell clustering identification was conducted using the Louvian community detection algorithm in Seurat, and cluster-specific marker genes were identified using the marker analysis (FindAllMarkers function, default parameters). Cell type annotation was based on comparison of cluster-specific marker genes to cell–type–specific genes from literature and public databases. Differential gene expression was assessed between IR and treatment conditions using the Wilcoxon method implemented in Seurat. Functional enrichment analyses of differentially expressed genes were performed against KEGG pathway annotations to identify enrichment of signaling programs and cellular processes.

Plasma small RNA sequencing data were processed and normalized using DESeq (21.48.2) to identify differentially expressed miRNAs across experimental groups. Cell–cell communication analyses were inferred using ligand–receptor interaction models implemented in the CellChat-v2 R package, which enables quantification of intercellular signaling strength and visualization of inferred communication networks across cellular compartments. Unless stated otherwise, a two-sided P value < 0.05 was considered statistically significant for all computational analyses.

## LIST OF SUPPLEMENTARY MATERIALS

### 1. Supplemental Materials and Methods

#### 2. Supplemental Tables

**Table S1.** Upregulated plasma miRNA after cardiac I/R injury

**Table S2.** Detailed echocardiography data (baseline vs. post-I/R) in four treatment protocols

**Table S3.** Detailed baseline echocardiography data in control sham mice with the Prevention Protocol

**Table S4.** Detailed echocardiography data (baseline and post-I/R) in WT and TLR7 KO male mice

**Table S5.** Detailed echocardiography data (baseline and post-I/R) in female WT and KO mice

**Table S6.** Quality assessment of nucleus preparations for snRNA-seq

**Table S7.** Nucleus number and percentage (cluster nuclei / total nuclei x 100%) of each cell cluster

**Table S8.** KEGG pathway analysis – enriched transcriptomic signatures of EC8, monocytes, and neutrophils

**Table S9.** Detailed echocardiography data (baseline and post-I/R) in male mice treated with ITGAM and control antibody

**Table S10.** Global marker genes

#### 3. Supplemental Figures

**Figure S1.** Enpatoran-induced cardioprotection in ischemic MI is TLR7-dependent.

**Figure S2.** TLR7 KO and Enpatoran treatment reduce MI sizes and preserve cardiac function after I/R in female mice.

**Figure S3.** Temporal dynamics of cardiac immune cell infiltration and cytokine-expressing cells after I/R injury.

**Figure S4:** Cardiac cell type-specific TLR family gene expression and dynamic changes in response to I/R and enpatoran.

**Figure S5.** ITGAM expression in endothelial cells.

**Figure S6.** ITGAM expression in neutrophils.

**Figure S7.** ITGAM expression in monocytes.

**Figure S8.** Blood neutrophil and monocyte dynamics and ITGAM expression following myocardial I/R injury.

**Figure S9.** Spleen and bone marrow neutrophil and monocyte dynamics and ITGAM expression following I/R injury.

**Figure S10.** Microfluidic adhesion assay of THP-1 adhesion to HCAEC.

**Figure S11.** *Itgam* and *Icam1* expression level.

**Figure S12.** STED super-resolution imaging of endothelial cells in the normal heart.

**Figure S13.** Detailed snRNA-seq analysis data.

#### 4. Legends for Videos 1 to 11

**Video S1–S5 (.mp4 format).** Microfluidic adhesion assay of THP-1 adhesion to HCAEC under different treatment conditions: **1:** non-treatment control; **2:** R848 treatment for both Mo and EC; **3:** R848 for both Mo and EC, with ITGAM-neutralizing antibody applied only to Mo; **4:** R848 for both Mo and EC, with ITGAM-neutralizing antibody applied only to EC; **5:** R848 and ITGAM-neutralizing antibody applied to both Mo and EC. EC, endothelial cell; Mo, monocyte.

**Video S6–S8 (.mp4 format).** STED super-resolution 3D imaging: immunofluorescence staining of heart sections showing ITGAM (red), ICAM1 (green), and cell membranes (WGA, magenta) in I/R cell–cell interface (**6**), I/R endothelial cells (**7**), and Sham heart (**8**).

**Video S9–S11 (.mp4 format).** Microfluidic adhesion assay of THP-1 adhesion to HCAEC under different treatment conditions: **9:** R848 for both Mo and EC, with ICAM1-neutralizing antibody applied only to Mo; **10:** R848 for both Mo and EC, with ICAM1-neutralizing antibody applied only to EC; **11:** R848 and ICAM1-neutralizing antibody applied to both Mo and EC. EC, endothelial cell; Mo, monocyte.

## ACKNOWLEDGMENTS

We thank Dr. S. Ament and Ms. M. Cortes of the Institute for Genome Sciences, University of Maryland School of Medicine, for expert assistance with nuclei isolation.

## Author contributions

Conceptualization: W.C., L.Z., Y.L. Methodology: Y.L., Y.Y, C.P., B.R., R.L., A.S., and Z.L. Investigation: Y.L., Y.Y, C.P., B.R., R.L., A.S., and Z.L. Funding acquisition: W.C. Project administration: W.C., L.Z., B.W. Supervision: W.C., L.Z., B.W. Writing – draft: Y.L., W.C. Writing – review & editing: Y.L. C.P., B.R., R.L., B.W., Z.L., L.Z., and W.C.

## Data and materials availability

Source data are provided and are deposited in the NIH GEO repository upon publication (plasma small RNA-seq: GSE148864; cardiac snRNA-seq: GSE316860). Additional datasets are available from the corresponding author upon request.

## SUPPLEMENTAL MATERIALS

### Supplemental Materials and Methods Experimental animals

All animal experiments conformed to the Guide for the Care and Use of Laboratory Animals published by the US National Institutes of Health(*1*) and were approved by the University of Maryland Institutional Animal Care and Use Committee (AUP-00001355). Nine to 12-week-old male or female C57BL/6J mice were purchased from The Jackson Laboratory (Bar Harbor, ME). TLR7 knockout (KO) (Tlr7^tm1Flv^/J, stock no.008380) mice were originally purchased from The Jackson Laboratory and have been bred in-house with C57BL/6J mice for more than 10 generations. All animals were housed at the veterinary facilities of the University of Maryland School of Medicine for at least one week before experiments in an air-conditioned, pathogen-free environment with free access to water and a bacteria-free diet. The operator (YL) was blinded to mouse genotype and treatment allocation until completion of all analyses. Randomization was performed using simple sequential numbering generated manually to determine group assignment.

### Myocardial ischemia-reperfusion protocol

Myocardial I/R injury was induced by surgical procedures as previously described(*2*). Briefly, mice were anesthetized by intraperitoneal injection of ketamine (120 mg/kg) and xylazine (8 mg/kg), intubated, and ventilated. Body temperature was maintained at about 37.5°C with a heating pad. A left intercostal thoracotomy was performed, and the left anterior descending coronary artery (LAD) was visualized and ligated with 7-0 silk sutures under a surgical microscope. For I/R injury, 45 minutes after LAD ligation, the ligature was released, and reperfusion was visually confirmed. Sham-operated mice underwent the same procedure without LAD ligation. Heart rate, electrocardiogram, and body temperature were closely monitored throughout the procedures. The operators (YL) were blinded to the mouse strain and intervention information.

### Assessment of myocardial ischemic area and infarct size

Twenty-four hours after reperfusion, LAD was re-ligated. Fluorescent microspheres (FluSpheres^TM^ polystyrene, Invitrogen, Carlsbad, California) were injected into the left ventricle while the ascending aorta was briefly clamped as previously described(*2*). The heart was isolated and cut into five sections with a thickness of 1mm for each section. The areas devoid of microspheres were designated as area-at-risk (AAR). To measure MI size, each section was stained with 1% triphenyltetrazolium chloride (TTC) (Sigma-Aldrich, St. Louis, MO). Infarct size was calculated as the ratio of infarcted area (MI/LV, %) over AAR (AAR/LV, %), *i.e.,* MI/AAR x 100%(*2*).

### Echocardiography

Echocardiography (ECHO) was performed using a Vevo2100 ultrasound system (VisualSonics; Toronto, Canada). In brief, mice were placed in the supine position on a heated platform with all electrocardiographic electrodes on the legs for recording. Mice were initially anesthetized with 2% isoflurane and then 1% during the ultrasound procedure to maintain a heart rate between 500 and 550 bpm. The mouse’s body temperature was maintained within a range of 37.0 °C +/- 0.5 °C. Hair was removed from the chest area using chemical hair remover before imaging.

Transthoracic echocardiography was performed within one week before the surgical procedure (baseline) and 24 hours after I/R injury. Echocardiography images were obtained in the short-axis B-mode and M-mode at the level of the mid-papillary muscles for analysis of systolic function and dimensions. Parameters include end-diastolic left ventricular internal diameter (LVIDd), end-systolic left ventricular internal diameter (LVIDs), LV ejection fraction (LVEF), and cardiac output (CO). Diastolic function was determined using B-mode at the parasternal long-axis view and apical four-chamber view and pulsed Doppler (PW) as previously described(*3*). The left atrial (LA) diameter was measured with a parasternal long-axis view. PW was used to obtain the early mitral inflow E wave (E). Long-axis and short-axis B-mode images were collected for speckle-tracking strain (S) and strain rate (S e’ sr) analysis. E/e’ sr was then calculated. Parameters were measured offline with VevoLab v3.2.6 (VisualSonics).

### Reagents

1. Administration of enpatoran Enpatoran, a highly selective TLR7/8 antagonist, was recently developed by Merck (*4*). WT or TLR7 KO mice were given 2 doses–approx. 2 hours apart–of either enpatoran (InvivoGen, San Diego, CA) or normal saline (NS) by intraperitoneal (i.p.) injection. To model clinical scenario of patients with acute coronary syndrome admitted for coronary stenting, four protocols of enpatoran administration were designed: 1) Prevention protocol (PP): the two doses of NS or enpatoran were given 1 hour before LAD occlusion (*i.e.,* before ischemia) and again right after reperfusion; 2) Treatment protocol (TP): the two doses were given 35 min during LAD occlusion (*i.e.,* during ischemia) and 2 hours after reperfusion; 3) Early rescue protocol (ERP): the two doses given 30 min and 2.5 h after reperfusion started (*i.e.,* early after ischemia); 4. Late rescue protocol (LRP): the two doses given 2 hours and 4 hours after reperfusion (*i.e.,* late after ischemia).
2. Administration of Integrin alpha M (ITGAM) neutralizing antibody WT mice were given an ITGAM neutralizing antibody (Cat# BE0007, Bio X Cell, USA) or an isotype control (Cat# BE0090, Bio X Cell, USA) one hour prior to I/R surgery.
3. Cell treatment BMDMs or HCAECs were pretreated with pre-warmed FBS-free RPMI 1640 medium containing 0.05% BSA and 1% Penicillin-Streptomycin for 2 hours at 37°C with 5% CO_2_. Cells were then treated by either medium only (as non-treatment control), R837 or R848 (1μg/ml, InvivoGen, San Diego, CA), or miRs (miR146a-5p, miR-3061-3p, miR-741-3p, miR-5113, miR-8116, miR-29c-5p, miR-1843-5p, and miR-8097 [5nM, 50nM, or 500nM, Coralville, IA], Lipofectamine 3000 (Invitrogen, Thermo Fisher Scientific, Waltham, MA), and enpatoran (500nM, InvivoGen, San Diego, CA) at 37 °C with 5% CO_2_ for 3 hours, 6 hours, or 18 hours.

### Collection of mouse blood and tissue samples

Animals were euthanized at 24 hours after I/R or sham procedure by inferior vena cava puncture under general anesthesia. Blood samples were collected in K2EDTA phlebotomy tubes (MiniCollect, Greiner Bio-One) and immediately processed by two-step centrifugation at 1,000 *g* and 10,000 *g* for 10 min at 4°C to obtain a cell-free plasma aliquot, which was stored at -80°C until further analysis. Tissues and thoracic and abdominal organ samples were collected sterilely and rinsed in cold PBS. Samples were immediately snap-frozen in liquid nitrogen and stored at - 80°C until further analysis.

### Quantification of plasma cardiac troponin I

Plasma cardiac troponin I (cTnI) levels were measured using the high-sensitivity mouse cardiac troponin-I ELISA immunoassay kit (Life Diagnostics, United Kingdom) and quantified in a SpectraMax Pro 5 plate reader (Molecular Devices, Sunnyvale, CA). The assay was performed in duplicates for each plasma sample.

### Single-nucleus RNA-sequencing

1. Isolation of nuclei from frozen heart tissues WT C57BL/6J mice underwent either Sham surgery with NS treatment (NS-Sham), I/R surgery with NS treatment (NS-I/R), or I/R surgery with enpatoran treatment (Enpa-I/R) (n=3 in each group). Animals were euthanized at 24 hours after surgery and left ventricular tissues just below the LAD ligation level were collected and frozen for later nucleus isolation. To prepare nucleus samples of high quality, a detergent-mechanical cell lysis method was used(*5*), involving three major steps: lysis, homogenization, and density barrier centrifugation. In a laminar hood, a ∼50-mg powdered heart tissue was placed into a pre-frozen BioPulverizer (BioSpec), then immediately added lysis buffer (10mM Tris-HCl, 10 mM NaCl, 3 mM MgCl_2_, 0.1% Nonidet P40, 1X RNase inhibitor, 1μl/ml DEPC). Tissue was disaggregated, flushing up and down, first using a 1 ml pipette tip. Homogenized tissue was diluted to 2 ml with lysis buffer and filtered through a 70 μm filter. Homogenate was spun down at 350 *g* for 5 min at 4°C. The pellet was washed twice with washing buffer (1X PBS, 1% BSA, 1X RNase inhibitor). Pellet was then resuspended and 500 μl of pellet suspension was diluted with 500 μl 5% iodixanol solution (5% iodixanol, 250 mM sucrose, 150 mM KCl, 130 mM MgCl_2_, 1X RNase inhibitor, and 60mM Tris-HCl, pH 8.0), filtered for the second time with a 20 μm mesh and placed on bottom of 500 μl layer of 8.5% iodixanol solution (8.5% iodixanol, 250 mM sucrose, 150 mM KCl, 130 mM MgCl_2_, 1X RNase inhibitor, 60mM Tris-HCl, pH 8.0). The density barrier was centrifuged at 10,000 *g* for 10 minutes at 4°C. The pellet was collected and washed with washing buffer, centrifuged at 350 *g* for 5 min at 4°C, and finally suspended in 50 μl washing buffer. Single-nucleus suspensions were counted and evaluated for integrity using Propidium Iodide in a MoxiGo cytometer using 650 nm filter. The nucleus count was adjusted to 1000 nuclei/ml.
2. Library preparation and sequencing Library preparation and sequencing were performed at the Institute for Genome Sciences, University of Maryland School of Medicine. A total of 16,500 nuclei were loaded into each well of a chromium microfluidics controller (10x Genomics) to target 10,000 cells/nuclei per sample. Sequencing libraries were generated using the Chromium Next GEM Single Cell 3’ Reagent Kits v3.1 (dual Index). Sequencing targeted 50,000 reads per nucleus. Samples were sequenced using Illumina paired-end 100 bp sequencing (NovaSeq 6000 S4 Lane - 2250M read pairs, 450 Gbp).
3. Nuclei quality control and data preprocessing To ensure high-quality nuclei were utilized for downstream single-nucleus analyses, we selected nuclei with a minimum of 100 genes and a maximum of 8000 genes expressed and retained genes that were expressed in at least 3 cells (**Fig. S13A-13B**). Since nuclei are often stressed, damaged, and undergo necrosis during sample preparation, we retained nuclei with mitochondrial gene proportions below 25% (**Fig. S13C**). Using these thresholds, we retained 55,876 high-quality nuclei ranging from 4,022 - 7,985 cells per sample (**Fig. S13D, Table S6)**.
4. Integration, batch effect correction, and normalization 55,876 high-quality cells from 9 cardiac tissue samples were analyzed using Seurat (v4.3.0) for normalization of the gene expression data, identification of variable gene features and dimensionality reduction using Principal Component Analysis (PCA) and Uniform Manifold Approximation and Projection (UMAP). Cells across multiple samples were integrated using integration anchors identified using the canonical correlation analysis method to remove potential batch effects, followed by data scaling, and dimensionality reduction of the integrated dataset.
5. Cell type clustering and labeling After data integration, cells were grouped into 9 major cell clusters, which were further refined into 37 different cell clusters using Louvian community detection algorithm in Seurat (v4.3.0). Cluster-specific marker genes (**Table S10**) were identified for each of the 37 clusters using the differential gene expression analysis (FindAllMarkers function, default parameters). These markers were then systematically compared against curated reference marker genes from the CellMarker database as well as cardiovascular-relevant literature, allowing for biologically meaningful annotation of each cluster. Using these cell annotations, we labeled the 37 cell clusters to identify eight cardiomyocyte sub-clusters, eight endothelial cell sub-clusters, six fibroblast sub-clusters, one smooth muscle cell cluster, three pericyte sub-clusters, one mesothelial cell cluster, one neuronal cell cluster, three lymphoid immune cell sub-clusters, and six myeloid immune cell clusters, wherein each cluster consisted of 72 - 4,973 cells per cluster (**Fig. S13E**).
6. Cell type-specific DEGs and enrichment analysis Differentially expressed genes (DEGs) were identified for each of the annotated clusters separately for comparisons between cells belonging to two different treatment conditions, namely NS-I/R vs NS-Sham and Enpa-I/R vs NS-I/R. Default methods available through Seurat (v4.3.0) utilizing the Wilcoxon Rank Sum test were used to identify DEGs with a minimum log2-scaled fold-change of 0.25 and for genes expressed in at least 25% of cells in either of the two groups. Over-representation analysis method was then used to assess and evaluate the overlap of the DEGs with known functional gene sets curated from Gene Ontology databases, Hallmark Pathways, and KEGG Pathways which enables the biological interpretation and functional significance of the DEGs identified for different cell clusters between different treatment conditions.
7. Cell-to-cell communication between cell types using in-silico ligand-receptor analyses To investigate intercellular communication among major cell types across experimental groups, we performed ligand-receptor-based cell–cell interaction analysis using the CellChat-v2 R package. CellChat infers intercellular signaling networks by integrating a curated database of known ligand–receptor pairs with scRNA-seq data, enabling the identification of active signaling pathways and quantification of communication strength between cell types. In total, 2,377 curated mouse L–R pairs from the CellChat database were evaluated. Ligands and receptors were considered expressed if their corresponding genes were detected in more than 50 cells within a given cell type. For this analysis, preprocessed and annotated scRNA-seq expression matrices from the three study groups (NS-Sham, NS-I/R, and Enp-I/R) were analyzed independently to preserve group-specific signaling features. Within each group, we constructed a cell-cell communication probability matrix using the computeCommuProb function, which calculates the likelihood of interactions between sender and receiver cell types based on co-expression of known ligand-receptor pairs. We subsequently performed identification of dominant signaling pathway, quantification of communication network properties, and pairwise group comparisons. For pairwise comparisons, we assessed dynamic changes in intercellular communication across experimental conditions and conducted differential network analyses between NS-Sham vs. NS-I/R and NS-I/R vs. Enp-I/R groups. These analyses enabled the identification of upregulated and downregulated signaling pathways and cell-type pairs that exhibit condition-specific alterations in communication. The results were visualized using network graphs, bubble plots, and heatmaps to highlight both global and cell–type–specific communication changes.

### Plasma small RNAseq

RNA was isolated using TRIzol LS from 100 μl of cell-free plasma and was analyzed by an Agilent 2100 Bioanalyzer with the Small RNA kit (Agilent Technologies, Santa Clara, CA). Six microliters of each RNA solution were used for small RNA sequencing library preparation. Libraries were sequenced on an Illumina NextSeq 500 (Illumina, San Diego, CA) and analyzed by Norgen Biotech Corp. The raw sequence data were analyzed using the exceRpt small RNAseq pipeline (Genboree Bioinformatics). Differential expression (DE) analysis was performed using R package DESeq (21.48.2), which incorporates empirical Bayes estimation into a negative binomial distribution(*6*). The log2 fold change and log p values from DE analysis were used to draw heatmaps using R package ComplexHeatmap (2.24.1). Target genes for selected miRNAs were predicted using the R package multiMiR and functional analysis on these predicted targets was performed using the R package clusterProfiler II. Sample would be excluded from DEG analysis if it is an outlier in principal component analysis.

### Human database analysis

Human single-cell RNA sequencing (scRNA-seq) data were obtained from a published study investigating the immune response to ST-segment-elevated myocardial infarction (STEMI) (European Genome-phenome Archive accession EGAS00001007021; dataset ID EGAD00001010064)(*7*). The dataset comprises peripheral blood mononuclear cells (PBMCs) collected from 38 healthy controls and 38 patients sampled at hospital admission and 24 hours after reperfusion (coronary stenting). Data preprocessing was performed using the Seurat R package following standard workflows. Overall gene expression patterns were visualized using UMAP across healthy controls and patients with STEMI before and after reperfusion. The expression of *ITGAM*, *TLR7*, and *TLR8* was examined to characterize transcriptional changes among the three groups. In addition, we assessed both the expression level and the proportion of ITGAM^+^ monocytes across conditions.

### Cell culture and treatment

1. Bone marrow-derived macrophage (BMDM) culture Bone marrow cells were cultured as described previously with minor modifications(*8*). Briefly, the bone marrow cells were flushed out from isolated tibia and femur bones using a 27G syringe by injecting cold DPBS with heparin (1 USP Unit/ml) into the lumen of the bones. The cells were resuspended in RPMI 1640 medium-supplemented with 20 ng/ml M-CSF (R&D systems, Minneapolis, MN), 1% Penicillin-Streptomycin, 10% heat-inactivated fetal bovine serum (FBS), and 5% horse serum, seeded in either 12-well plate (5X10^6^ cells/well) 96-well plate (2X10^5^ cells/well) and incubated at 37°C with 5% CO_2_. The culture media were changed 48 hours after seeding, and the cultures were used for treatment the following day.
2. Human coronary artery endothelial cell (HCAEC) culture Human coronary artery endothelial cells (HCAECs) were purchased from Lonza (Morristown, NJ) and cultured as described previously with minor modifications(*9*). HCAECs were cultured in EBM®-2 Basal Medium (Cat.# CC-3156) and EGM®-2 SingleQuots® Supplements (Cat.# CC-4176), containing human epidermal growth factor (hEGF), vascular endothelial growth factor (VEGF), R3-insulinlike growth factor-1 (R3-IGF-1), ascorbic acid, hydrocortisone, human fibroblast growth factor (hFGF-b), gentamicin/amphotericin-B, and 5% FBS. For the experiments, cells at passages three to six were used and the serum concentration was decreased to 2% FBS to prevent excessive cell proliferation. Cells were cultured at 37°C with 5% CO_2_.
3. THP-1 culture and human neutrophil isolation The human pro-monocytic cell line THP-1 was maintained using RPMI 1640 (ATCC modification, Cat# A1049101, Gibco) supplemented with 10% FBS and 1% Penicillin/streptomycin in a 5% CO_2_ incubator at 37°C. To obtain neutrophils, blood was obtained from healthy donors after consenting. Neutrophils were purified using the EasySep direct human neutrophil isolation kit (StemCell Technologies, Cat# 19666) as per the manufacturer’s instructions. Blood was utilized within 6 h after the blood draw. Protocols were approved by the institutional review board at the University of Maryland, Baltimore (HP-00081592). Isolated neutrophils were resuspended in Iscove’s Modified Dulbecco’s Media with 20% FBS (Gibco, ThermoFisher Scientific, Cat# SH3022801).

### Cytokine ELISA

IL-6 levels in culture media or serum were tested with mouse IL-6 DuoSet and CXCL2/MIP-2 ELISA kits (R&D Systems, Minneapolis, MN). The samples were prepared by centrifugation at 12,000 rpm for 5 minutes.

### RNA extraction and qRT-PCR

Total RNA was extracted from BMDMs using TRI Reagent (Sigma-Aldrich) and was resolved in DEPC-treated water (Invitrogen). The concentration of RNA was determined by a NanoDrop spectrophotometer (Thermo Scientific Inc., Waltham, MA). 1 μg of RNA was used for reverse transcription. qRT-PCR was performed as described previously(*8*). In brief, cDNA was synthesized using M-MLV reverse transcriptase and random hexamer primers (Promega, Madison, WI). The real-time PCR was performed with GoTaq SYBR qPCR master mix (Promega, Madison, WI) using a QuantStudio™ 3 Real-Time PCR cycler (Applied biosystems, Thermo Fisher Scientific).

Gene expression was calculated using the relative CT method normalized to GAPDH (2^-ΔΔ^Ct), and the fold changes in the treatment groups were determined relative to the non-treatment control. The following are the primer sets for the qPCR: mouse Gapdh: (Forward: 5′-AACTTTGGCATTGTGGAAGG-3′, Reverse: 5′-GGATGCAGGGATGATGTTCT-3′); mouse Itgam: (Forward: 5’-CCGTCTGCGCGAAGGAGATA-3’, Reverse: 5’- CGCCTGCGTGTGTTGTTCTT-3’).

### Histology and Immunofluorescence Staining

After the mice were euthanized, the hearts were excised and cut into five sections (1mm/section, from basal to apical). The sections were fixed in 10% neutral buffered formalin and then embedded in paraffin. Three 5 μm-thick sections of basal pieces were sliced, deparaffinized (xylene [Fisher Scientific, Fair Lawn, NJ]), and dehydrated (100, 95, 70, and 50% ethanol, sequentially, Fisher Scientific, Fair Lawn, NJ) for staining.

Heart sections were stained with ITGAM (Invitrogen, PA5-90724, 1:200, RRID: AB_2806205), together with either CD31 (R&D, AF3628, 1:100, RRID: AB_2161028), Ly6G (Invitrogen, 16-9668-82, 1:100, RRID: AB_2573128), or CCR2 (Novus, NBP2-35334, 1:100, RRID: AB_3294607). Cell nuclei were stained with 4’,6-diamidino-2-phenylinodole (Sigma-Aldrich, F6057, mounting medium).

The secondary antibodies were donkey anti-rabbit IgG (Invitrogen, A32790, 1:1000, RRID: AB_2762833), donkey anti-goat IgG (Invitrogen, A21447, 1:1000, RRID: AB_2762833), goat anti-rabbit IgG (Invitrogen, A1108, 1:1000, RRID: AB_143165), goat anti-rat IgG (Invitrogen, A21247, 1:1000, RRID: AB_141778), goat anti-mouse IgG (Invitrogen, A21235, 1:1000, RRID: AB_2535804).

To quantify immunofluorescence-stained cells of heart section, images of the left ventricle were acquired using the “Large Image” automated tiling function in NIS-Elements acquisition software (Nikon) on a Nikon A1 confocal microscope equipped with a 20X objective. Images of heart section were reconstructed and processed using Imaris and NIH ImageJ software. In Imaris, cell numbers were quantified by segmenting individual cells into discrete objects using watershed-based algorithms. Segmentation was performed based on nuclear (DAPI) and/or marker signals, depending on the cell type. The Imaris Cell Wizard or Spots module was applied to detect and segment individual cells, followed by manual refinement to resolve closely apposed or overlapping cells. Within each defined cell type, ITGAM expression was then assessed to identify ITGAM-positive cells. The Imaris statistics module was subsequently used to quantify total cell numbers and the number of target cells co-expressing ITGAM, and the resulting data were exported for downstream analysis. In parallel, NIH ImageJ was used with a custom-written script to independently quantify immunofluorescence signals. Briefly, for each heart section, the total number of nuclei was first determined based on DAPI staining. Target cell populations were then identified using cell type–specific markers, and the proportion of target cells expressing ITGAM were quantified by applying intensity thresholds defined from sham controls. Quantification was performed on 5–6 serial sections per heart spanning from basal to apical regions. For each staining, data from all sections of a given heart were combined to generate a single quantitative value per animal.

### Immunofluorescent Staining of ITGAM and ICAM1 in vitro and image analysis

ITGAM and Intercellular Adhesion Molecule 1 (*ICAM1*) expression in BMDMs and HCAECs was detected using a confocal microscope (Ti2, Nikon, Japan) and displayed as representative images. The intensity of *Itgam* expression in the stimulated cells was also quantified using an ImageXpress PICO Automated Cell Imaging System (Molecular Devices). The cultured cells were seeded in an 8-well chamber slide (Nunc Lab-TekII, ThermoFisher Scientific) for a confocal microscope and were seeded in a Poly-D-lysine-coated 96-well plate for an ImageXpress PICO system. After treatment, the cells on the slide or plate were fixed with 4% paraformaldehyde and proceeded to immunofluorescent staining using blocking buffer (0.1% Triton x-100, 5% BSA, 5% NGS in PBS) and anti-ITGAM antibody (PA5-90724, Invitrogen, 1:200) and ICAM1 antibody (MA 5407, Invitrogen, 1:100). For nuclear staining, DAPI was used in a mounting medium (Fluoroshield with DAPI, Millipore SIGMA) during confocal imaging. After staining, the cells were covered with cover slips and their images were analyzed using NIS-Elements (Nikon, Japan). For imaging with the ImageXpress PICO system, Hoechst33342 (Cell Signaling Technology, MA) in PBS was incubated with the stained cells for 10 minutes following secondary antibody incubation. After staining, each well in the 96-well plate was filled with 200 µL of cold PBS, and images were acquired using the PICO imaging system (CellReporterXpress, Molecular Devices, CA, USA). The cell numbers were quantified based on the nucleus signal. Total integrated cell intensity was quantified as the sum of the CD11b signal within the cell.

### Flow cytometry

Cells from 30μL blood, spleen (5mL resuspension), and bone marrow (1mL resuspension) (tibia) were collected, washed once with PBS at 400g, 5min, followed by red blood cell lysis with 2mL ACK lysis buffer for 5min (blood), 2min (spleen), and 1min (bone marrow) at room temperature. All cells from 30μL blood, 50μL of splenocytes, and 50μL bone marrow cells were subjected to the following staining and FACS detection. Cells were washed once with PBS and stained with 1:500 Zombie Red for viability detection at 4°C for 15min. After staining, wash once with FACS buffer (PBS with 2% FBS) and block with anti-CD16/32 at 1:100 for 10min at 4 °C. Antibody cocktail was then added as 1:500 CD45 AF 700, 1:500 Ly6G Sparkblue 550, 1:500 Ly6C BV570, 1:500 CD11b PE F4/80 Real blue 705, and incubated at 4°C for 30 minutes. Wash once with FACS buffer, the samples were then fixed with 1x IC staining buffer (Thermo, eBioscience # 00-8222-49) overnight at 4°C, washed once with 1x permeabilization buffer (Thermo eBioscience #00-8333-56), and resuspended with 300μL FACS buffer (PBS with 2% FBS) and record data for the same volume (128μL).

### Microfluidic adhesion assays

HCAEC monolayers were grown in Vena8 Endothelial+ Biochips (Cellix Ltd, Dublin, Ireland, internal channel dimensions: length 20 mm, width 0.8 mm, height 0.12 mm) as previously described(*10*) and treated by R848 (1 μg/ml) with/without enpatoran (500nM) for 18 h prior to the adhesion assay. THP-1 or fresh isolated human neutrophil cells were treated with R848 (1 μg/ml) alone, or together with enpatoran (500nM) for three hours at 37°C before the experiment, followed by FITC anti-human CD45 antibody staining (Biolegend, San Diego, USA, Cat#304006, RRID: AB_314394) for 1 hour. ITGAM (BioXCell, BE0007, 25μg/ml, RRID: AB_1107693) or ICAM1 neutralizing antibody (BioXCell, BE0020, 25μg/ml, RRID: AB_1107639) THP-1 or fresh isolated human neutrophil cells were perfused for 15 min at 0.5 dyn/cm^2^ through the biochip channels containing HCAEC monolayers, using ExiGo^TM^ Nanopumps (Cellix Ltd, Dublin, Ireland). THP-1 or neutrophil adhesion was monitored using a Nikon Eclipse Ti microscope (Nikon, Tokyo, Japan). Images were taken in 10 representative fields (10X object) at the centerline of each channel. Adhesion levels were quantified using the mean number of 10 pictures for each condition.

### Stimulated Emission Depletion (STED)

Super-resolution images were acquired with the Abberior Facility Line STED microscope at the Confocal Microscopy Core at UMB. STED imaging was performed using an Olympus 60x/1.42 STED-UPLXAPO60xO oil immersion lens. ITGAM and ICAM1 were labeled using similar primary antibodies as mentioned before, but specific secondary antibodies: Abberior STAR Orange (anti-mouse IgG, 1:200) and STAR Red (goat anti-rabbit IgG, 1:200) were excited at 561nm and 640nm, respectively, and depletion was performed with a 775nm pulsed laser with a gating of 1-7ns and a pixel dwell time of 5µs. The emission from both channels was collected with the matrix detector and the images were post-processed with the Abberior Imspector software. Each line was scanned 15-30 times during acquisition (line accumulations). The three-dimensional STED data were acquired using the Easy3D module with voxel sizes set to 30nm, and the pinhole was set to 1AU.

## SUPPLEMENTAL TABLES

**Table S1.**
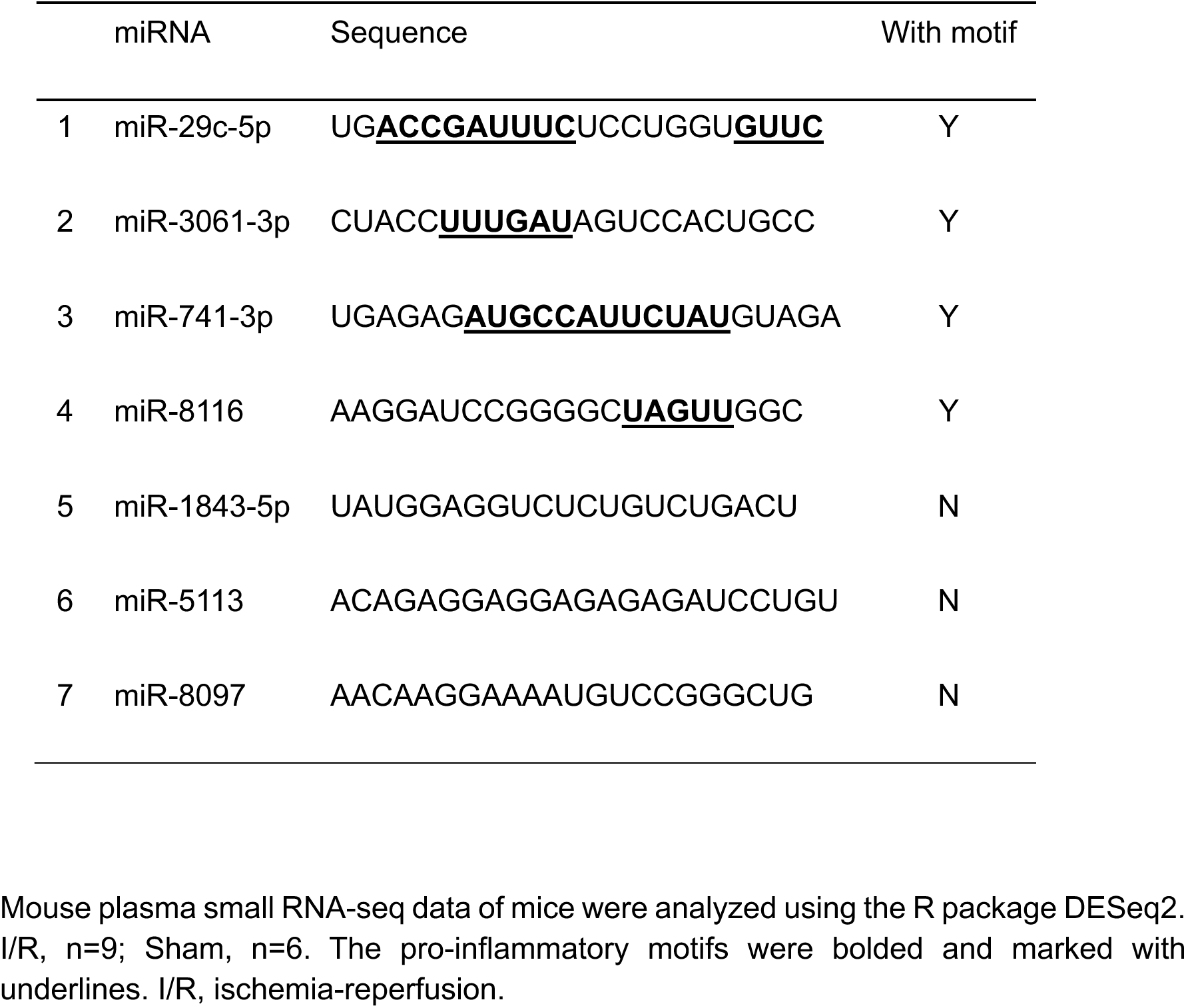
Upregulated plasma miRNA after cardiac I/R injury.

**Table S2.**
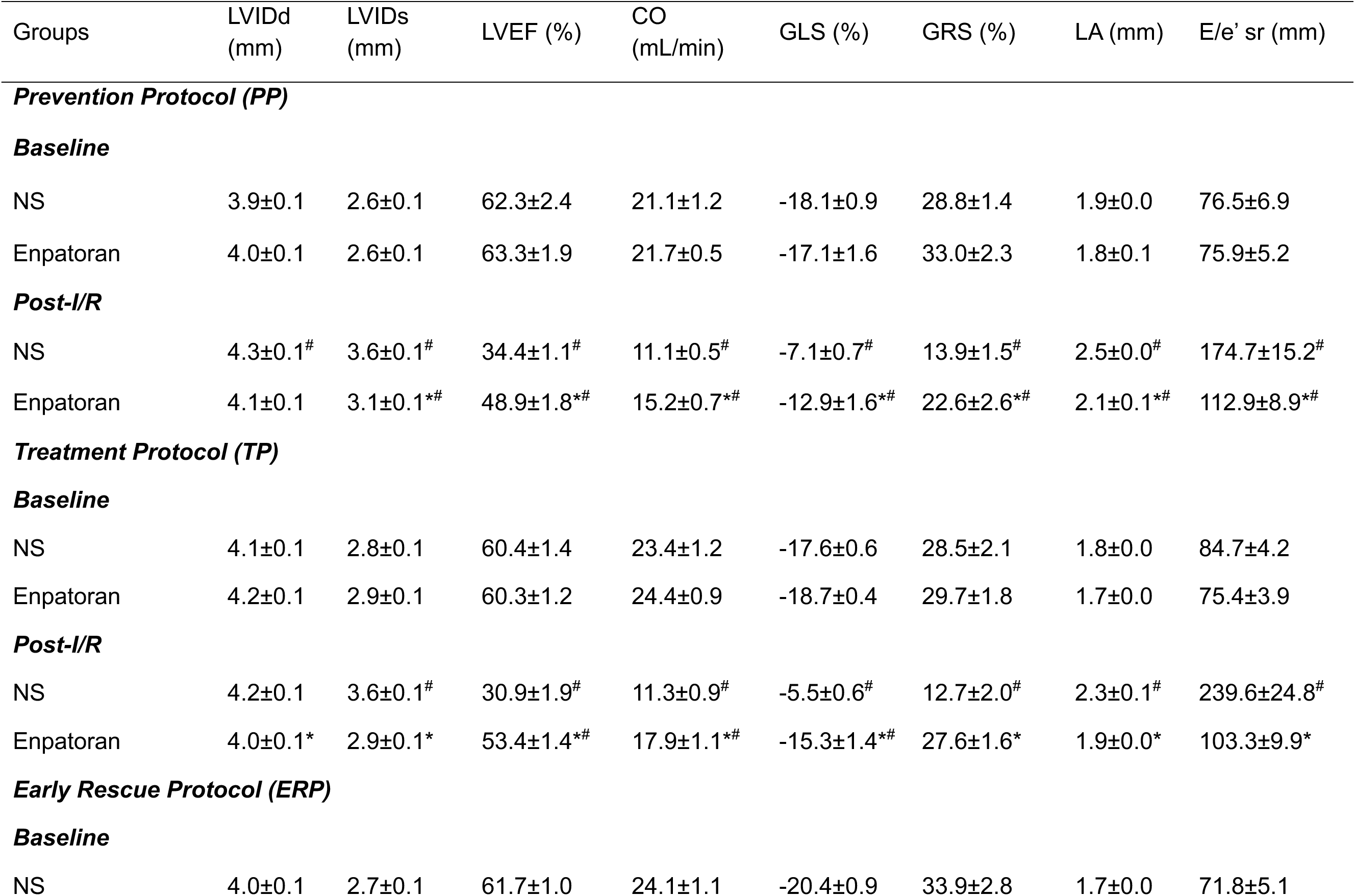
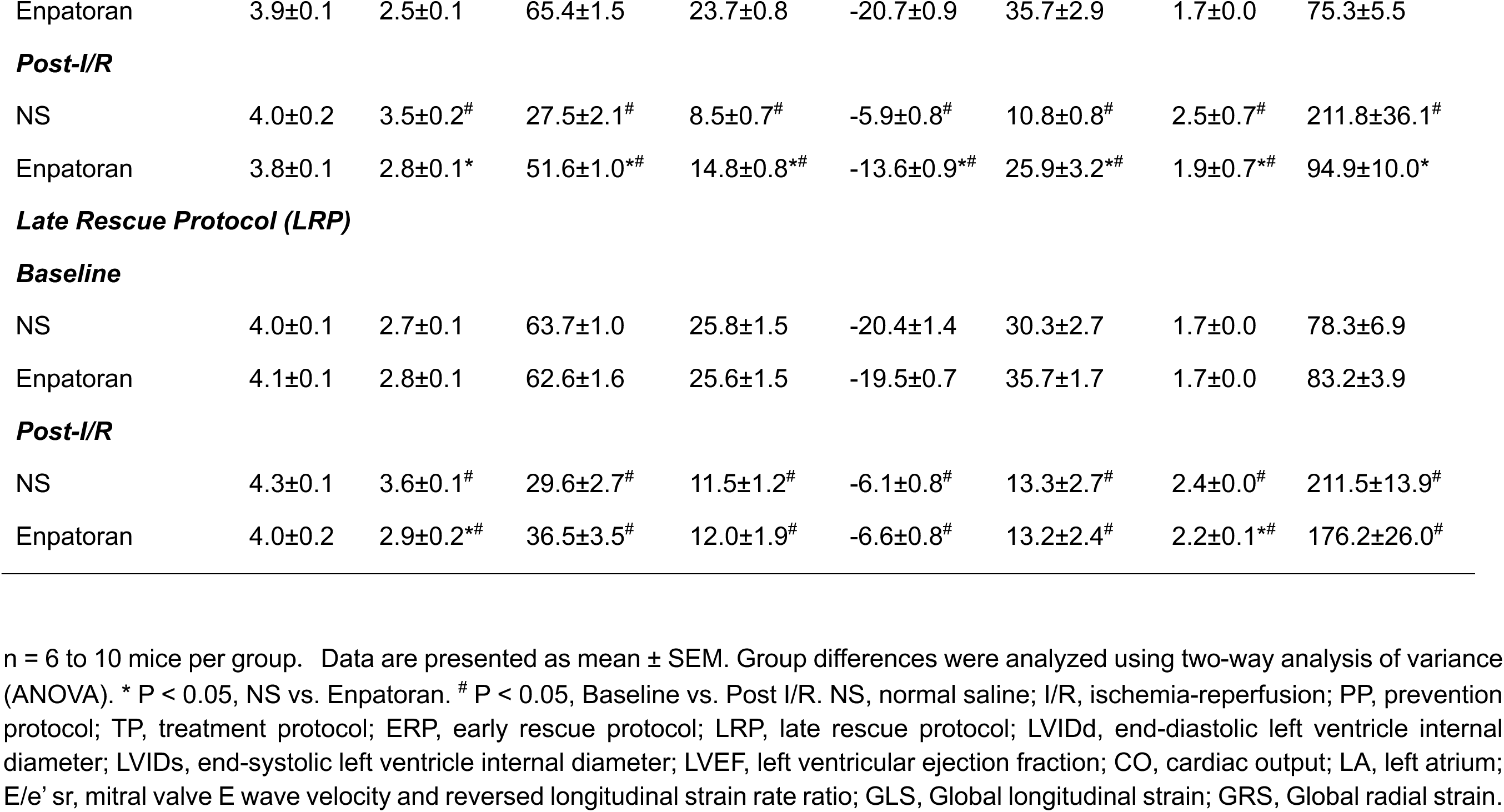
Detailed echocardiography data (baseline vs. post-I/R) in four treatment protocols.

**Table S3.**
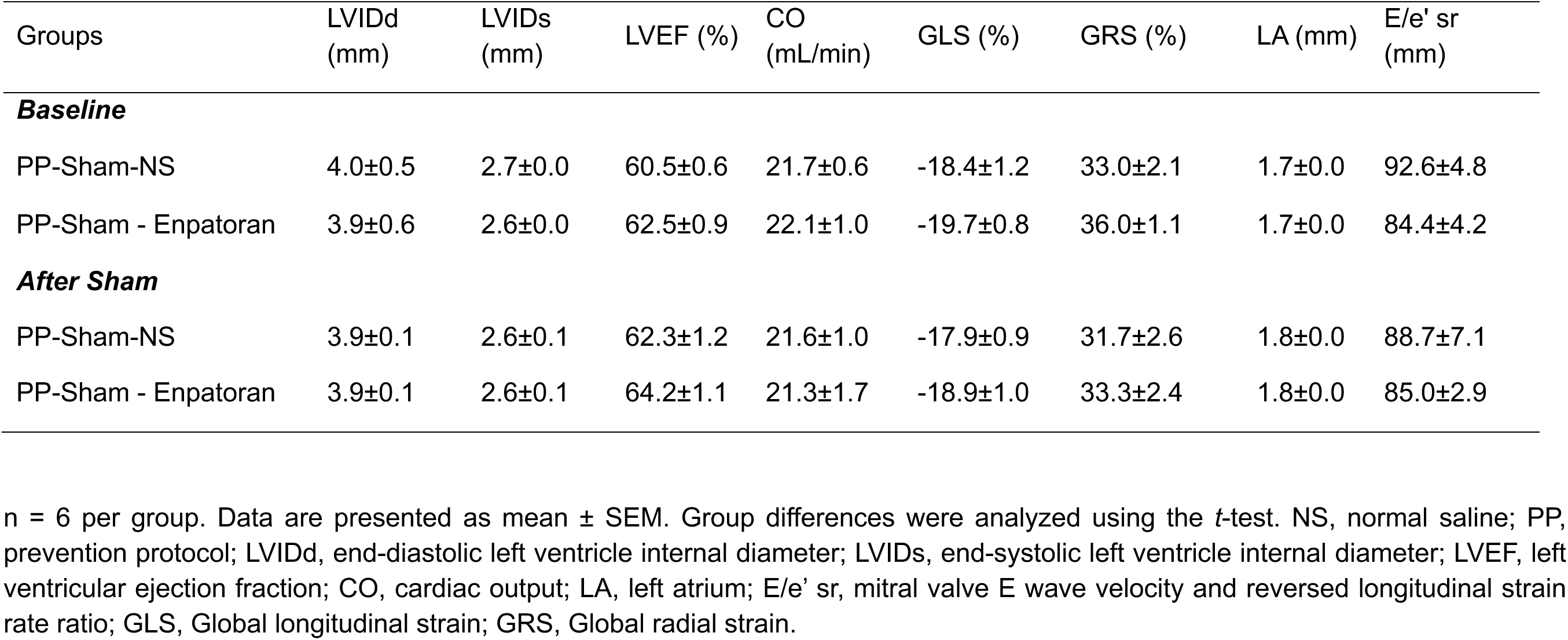
Detailed baseline echocardiography data in control sham mice with the Prevention Protocol.

**Table S4.**
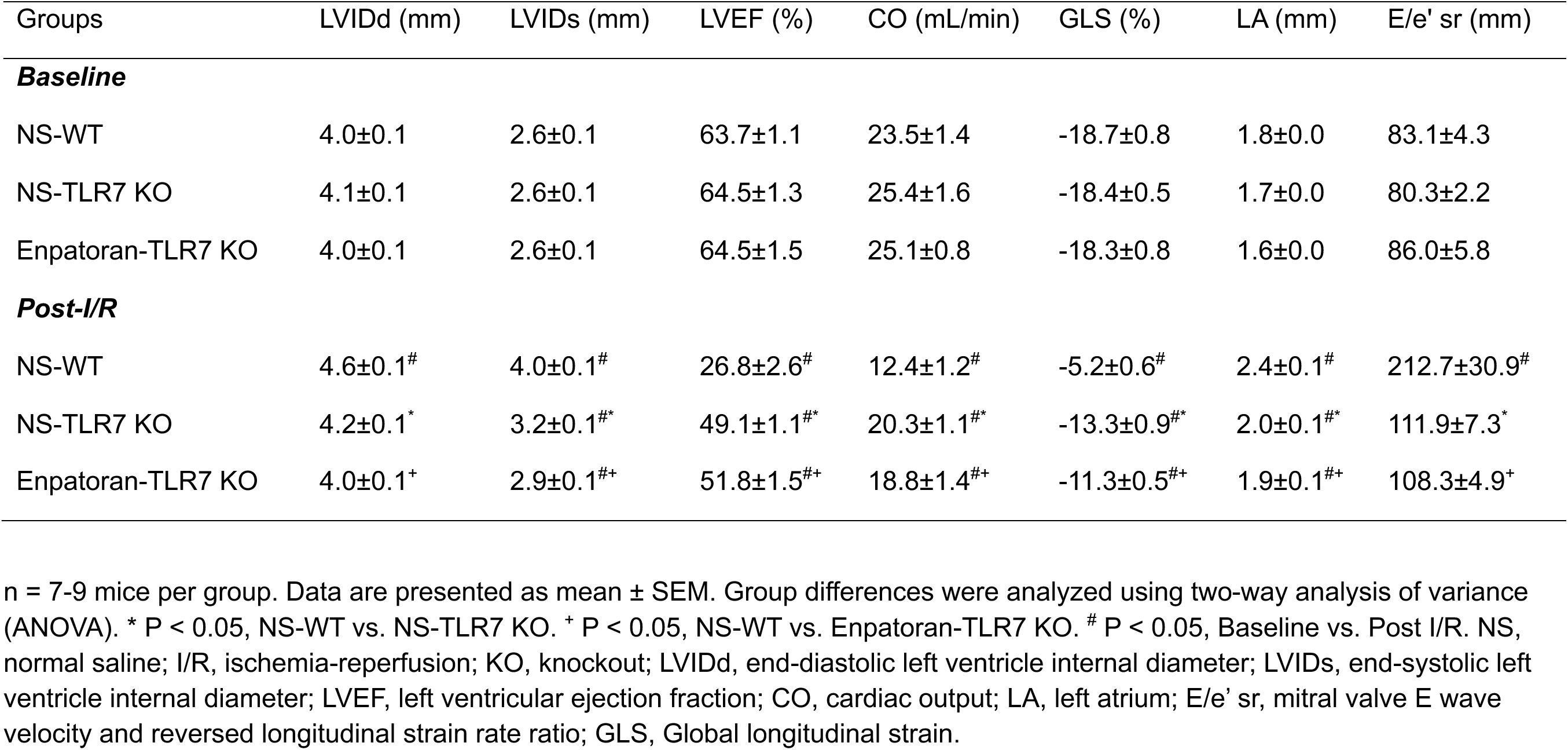
Detailed echocardiography data (baseline and post-I/R) in WT and TLR7 KO male mice.

**Table S5.**
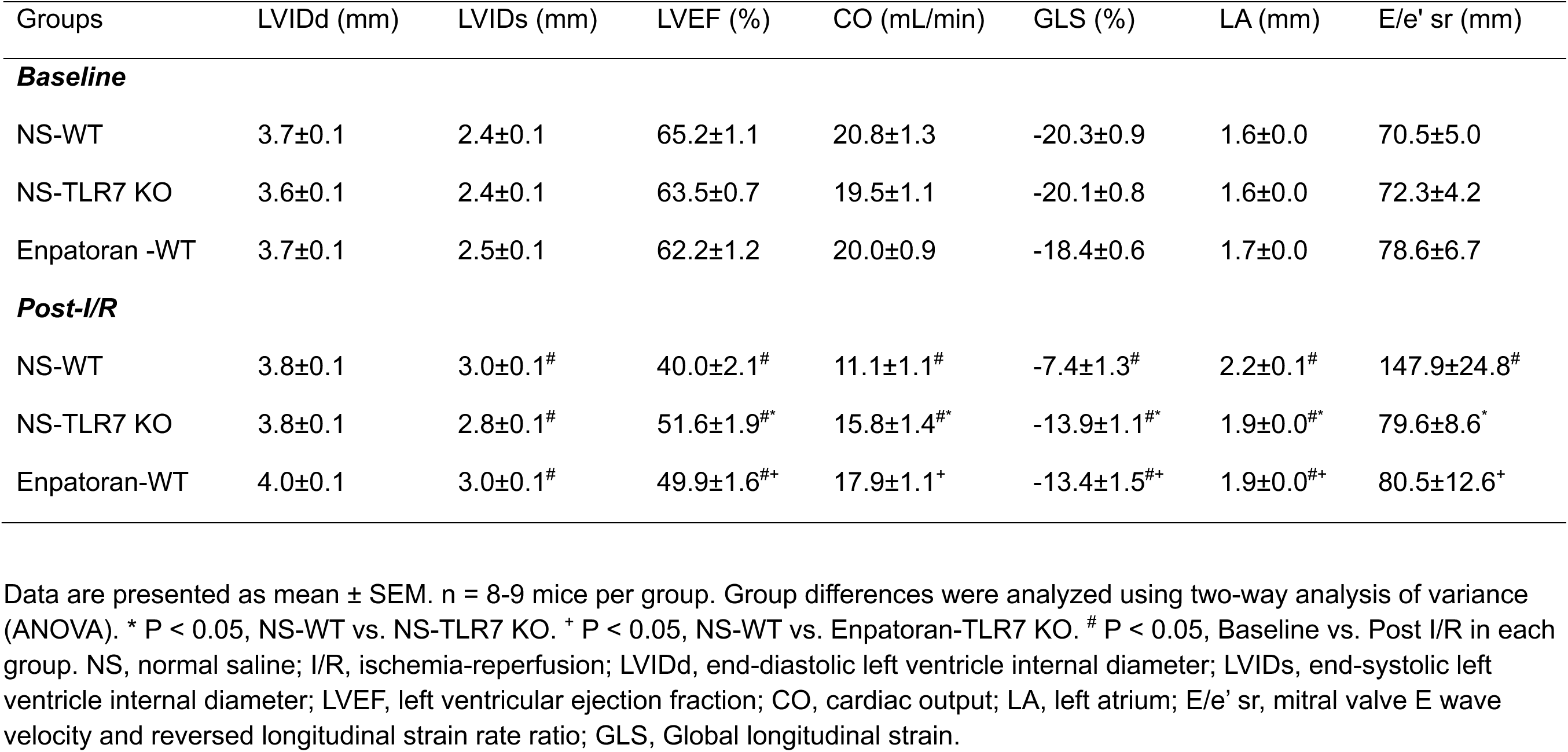
Detailed echocardiography data (baseline and post-I/R) in female WT and KO mice.

**Table S6.**
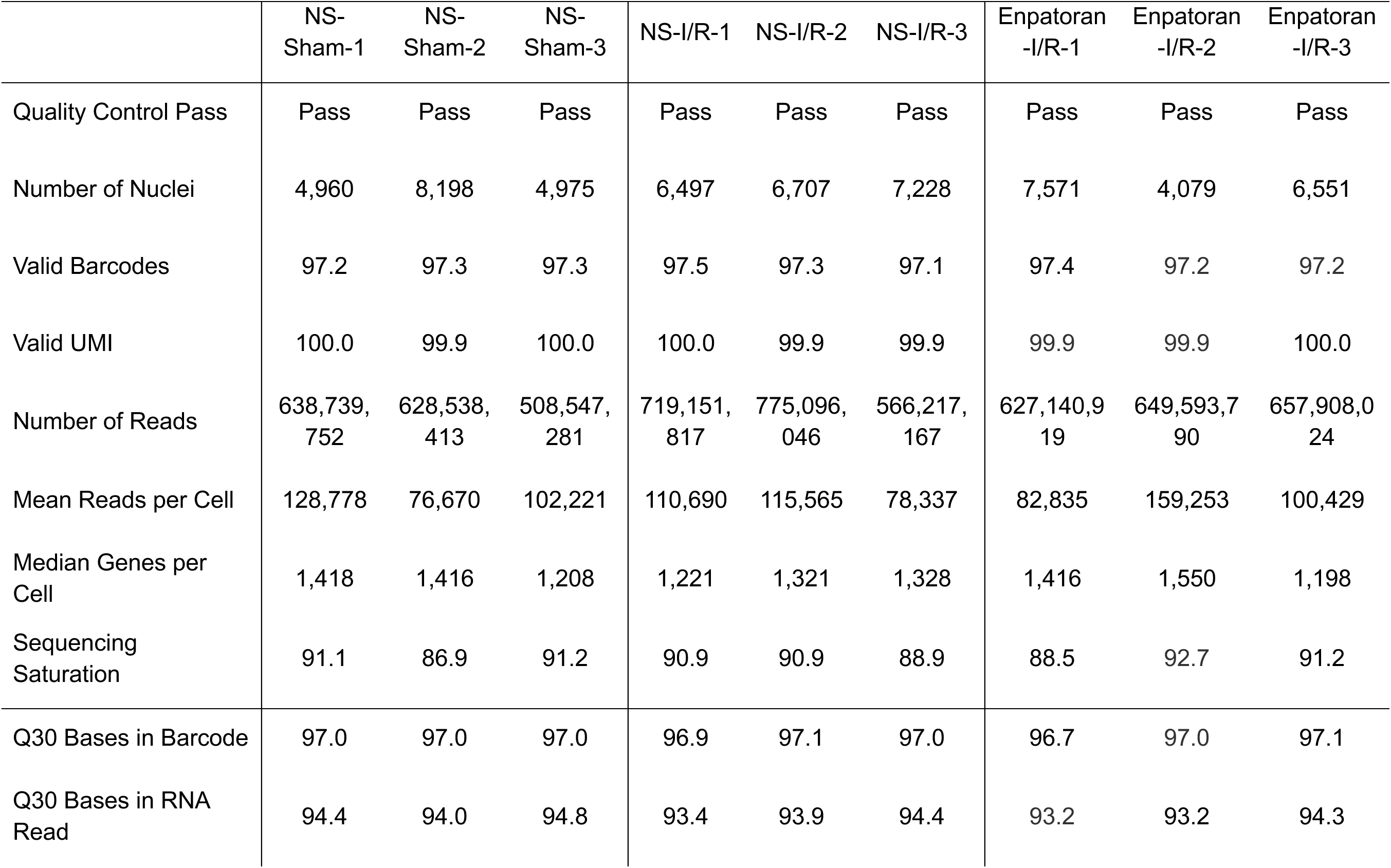
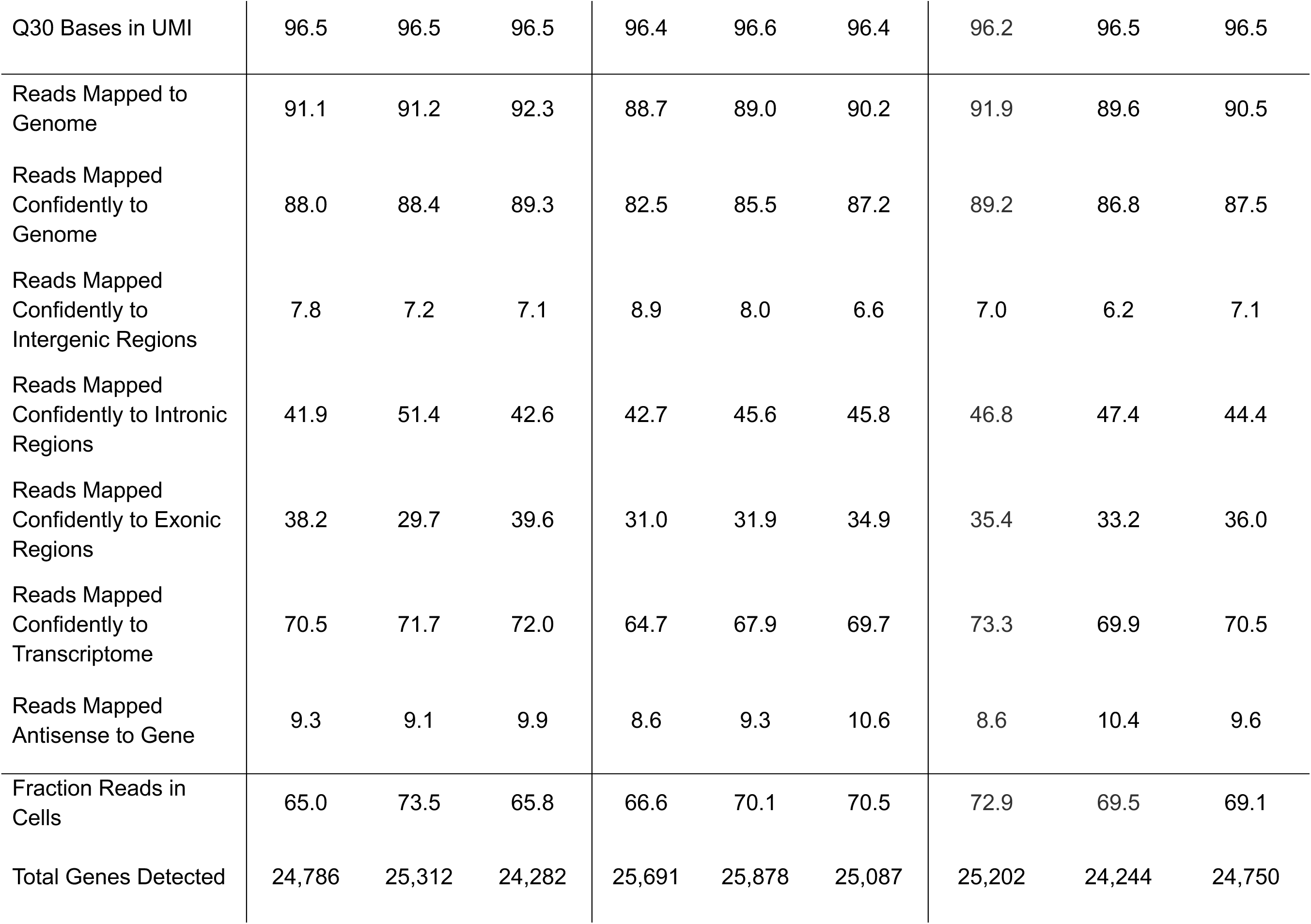
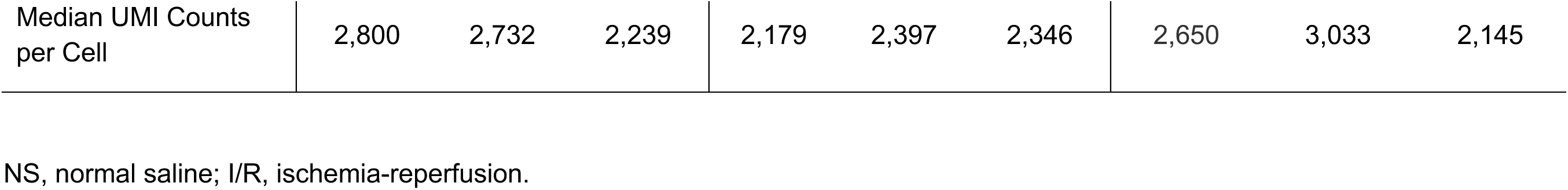
Quality assessment of nucleus preparations for snRNA-seq.

**Table S7.**
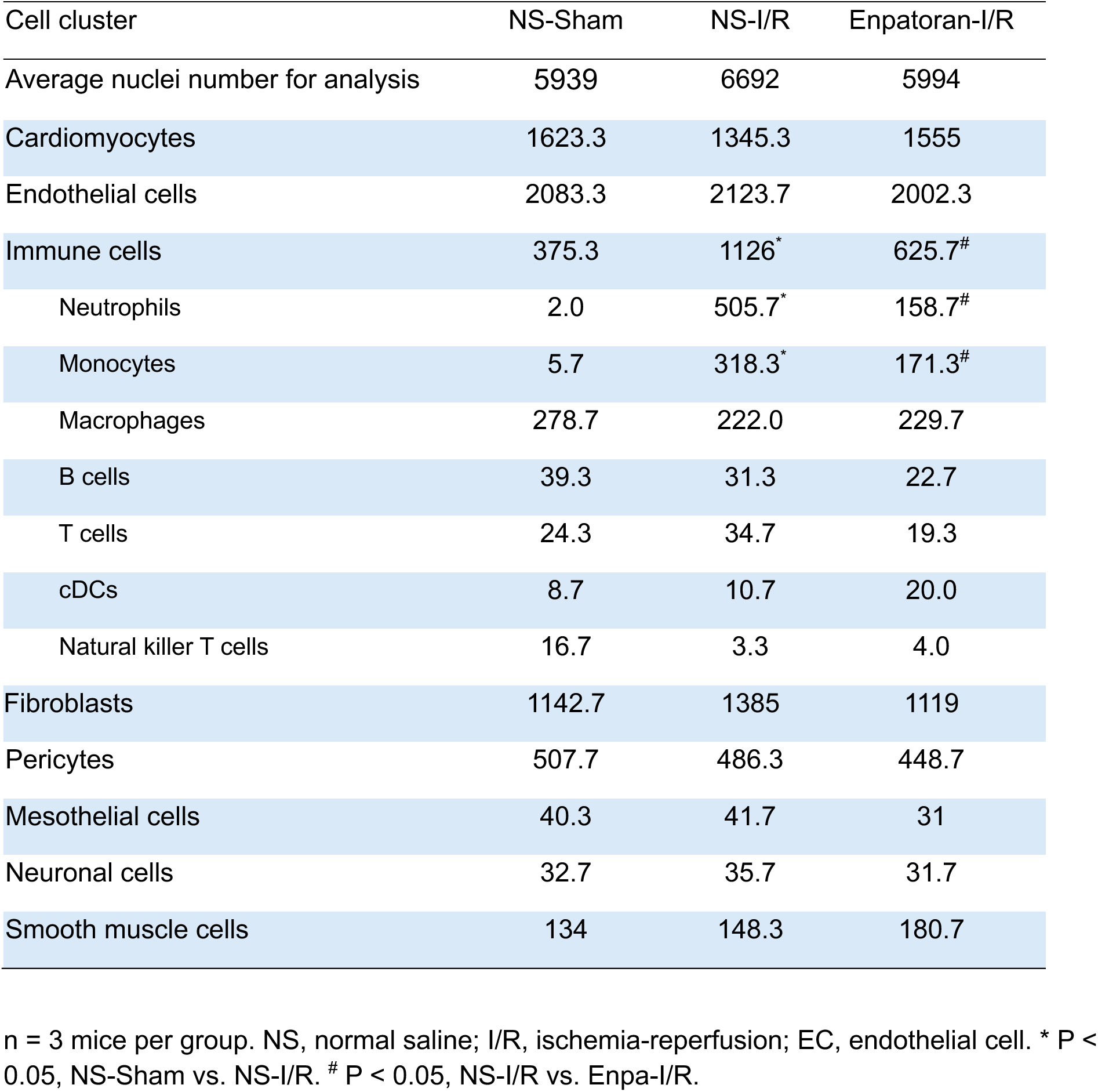
Nucleus number and percentage (cluster nuclei / total nuclei x 100%) of each cell cluster.

**Table S8.**
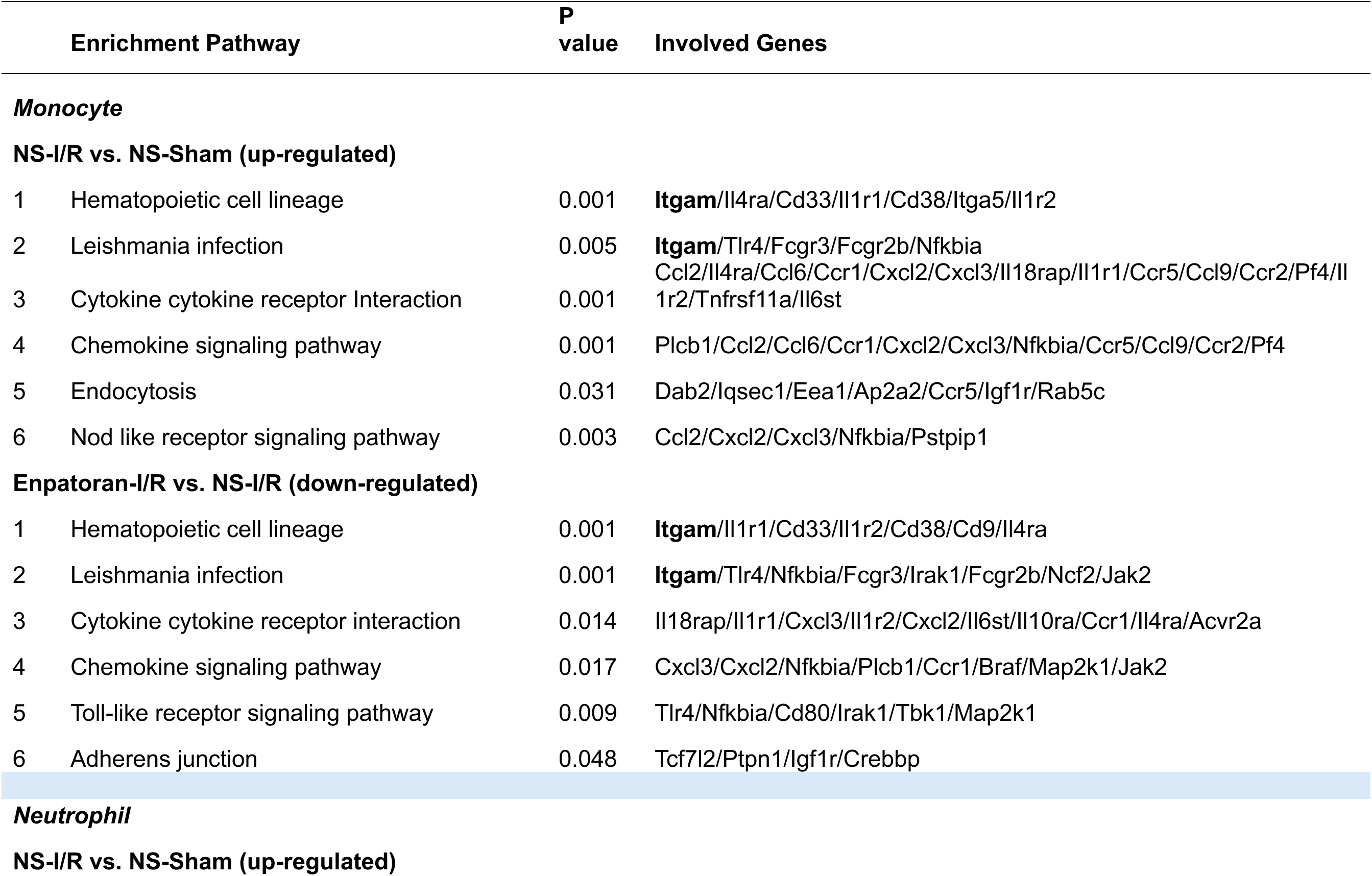
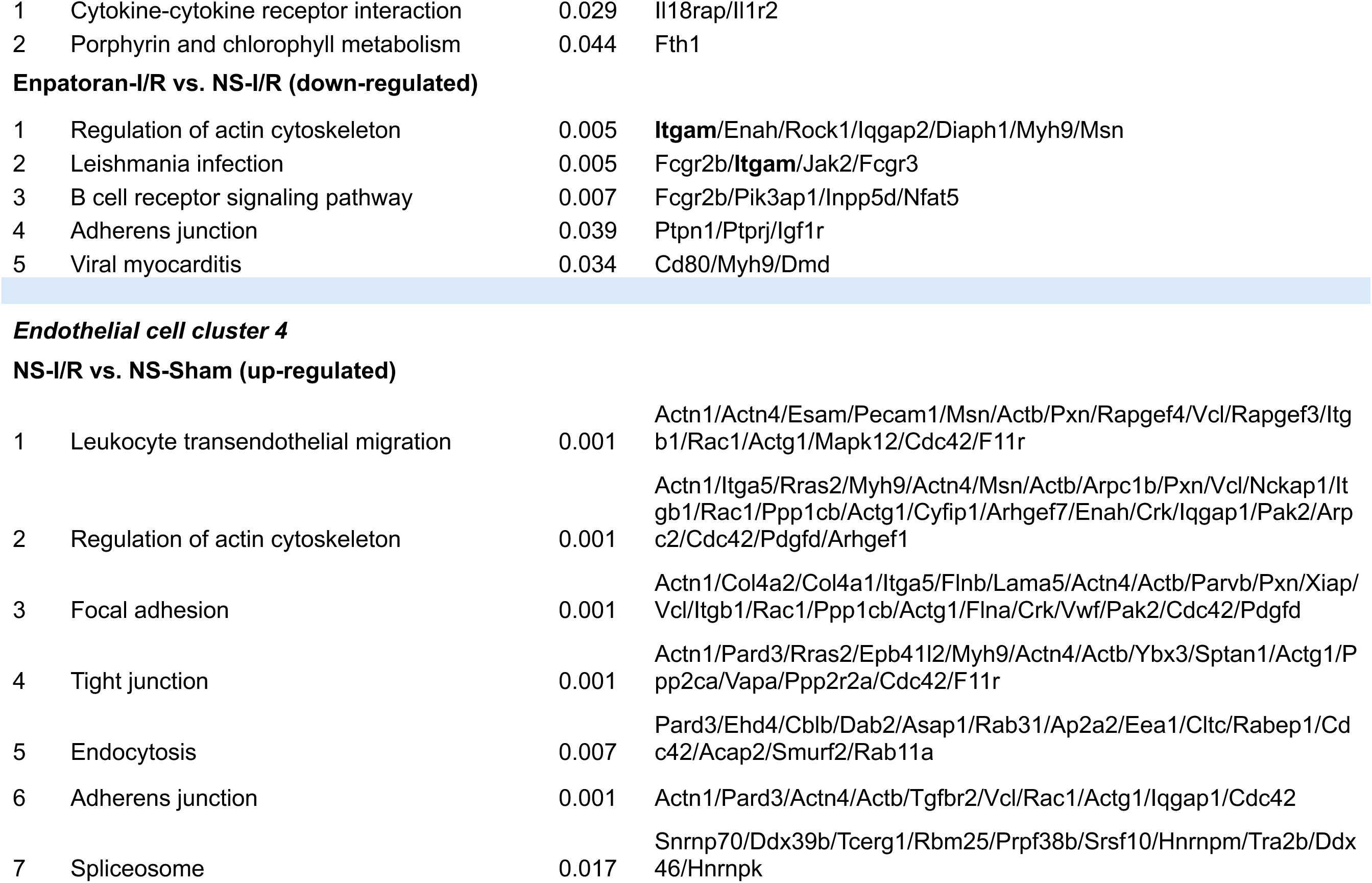
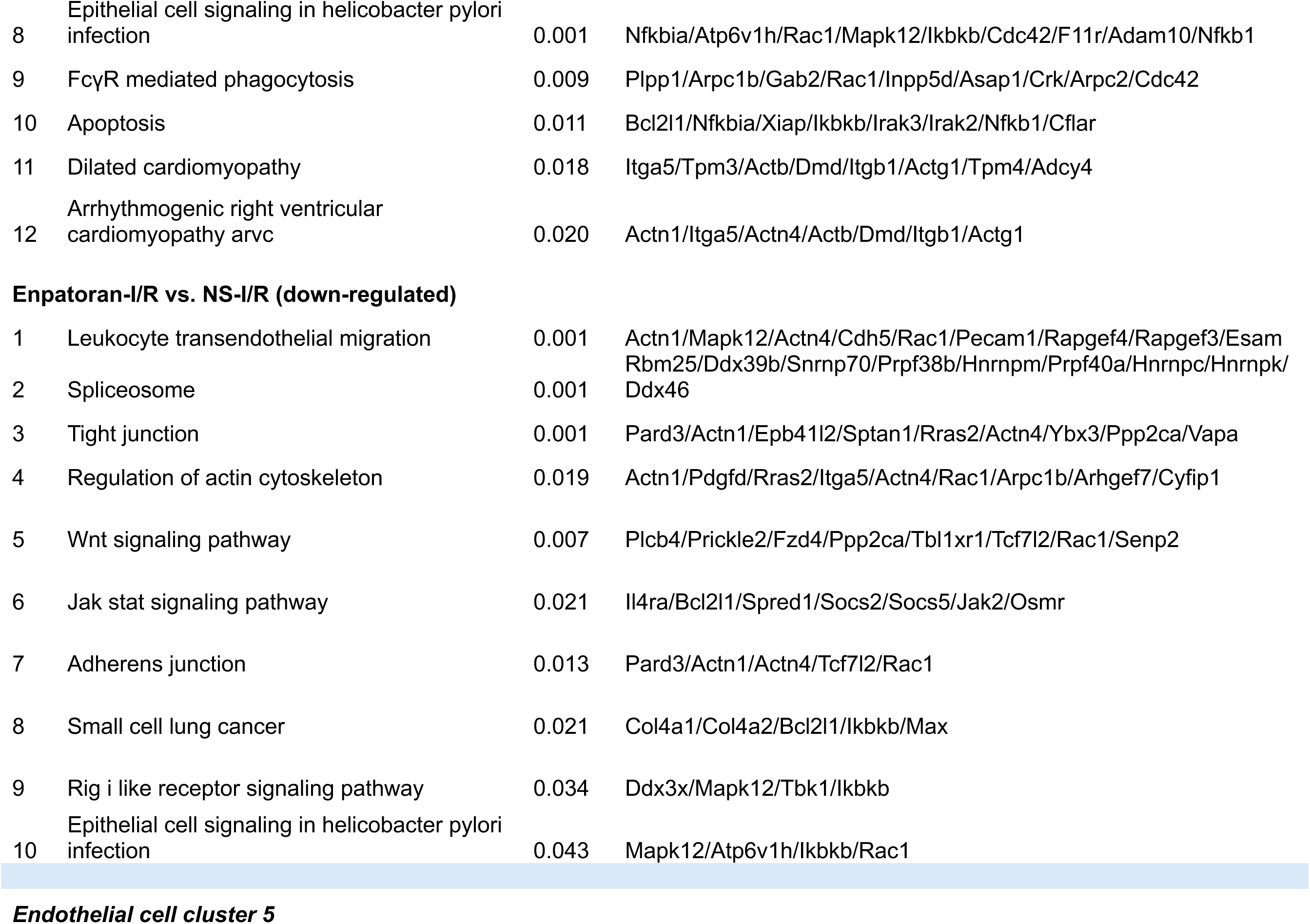
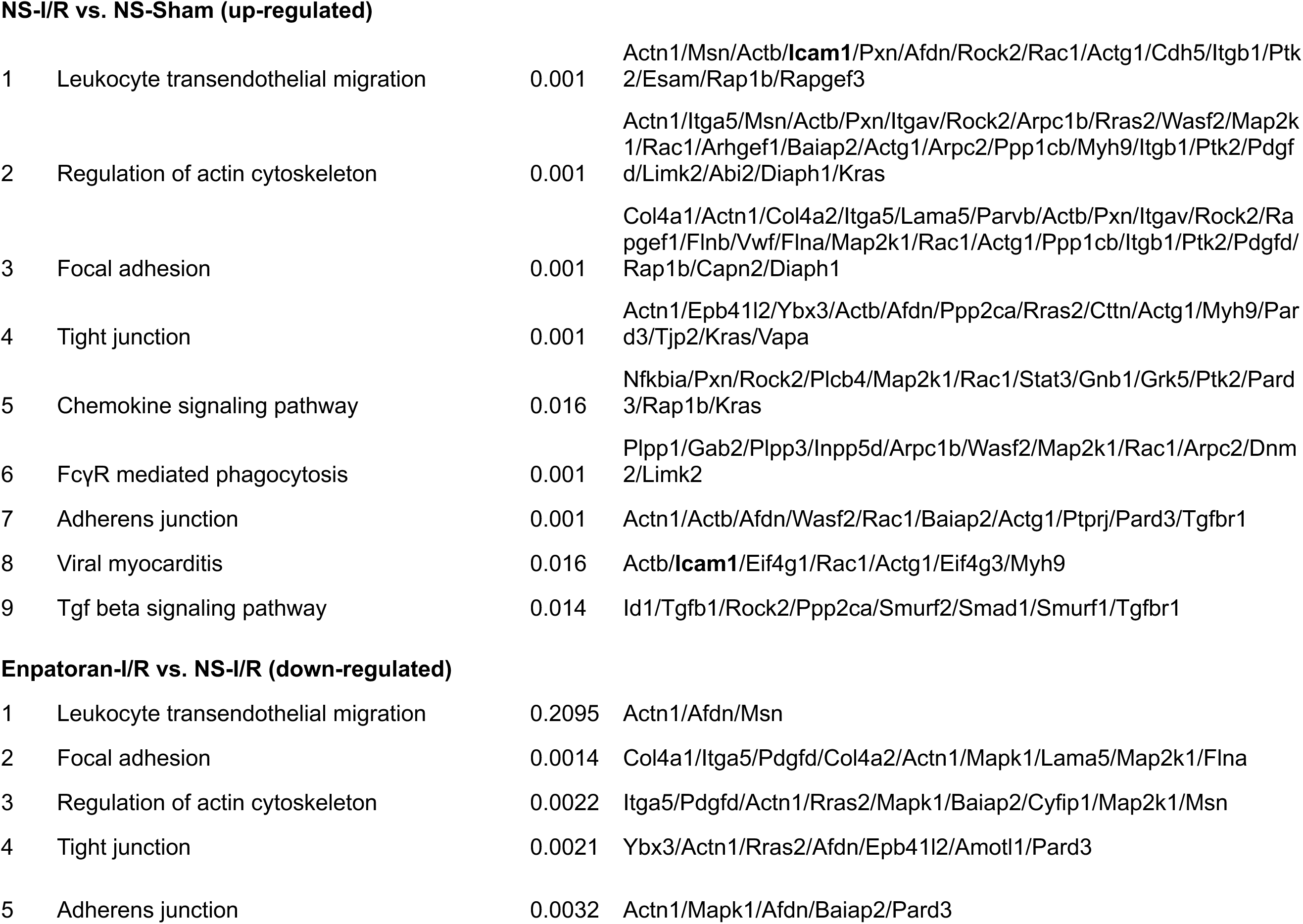
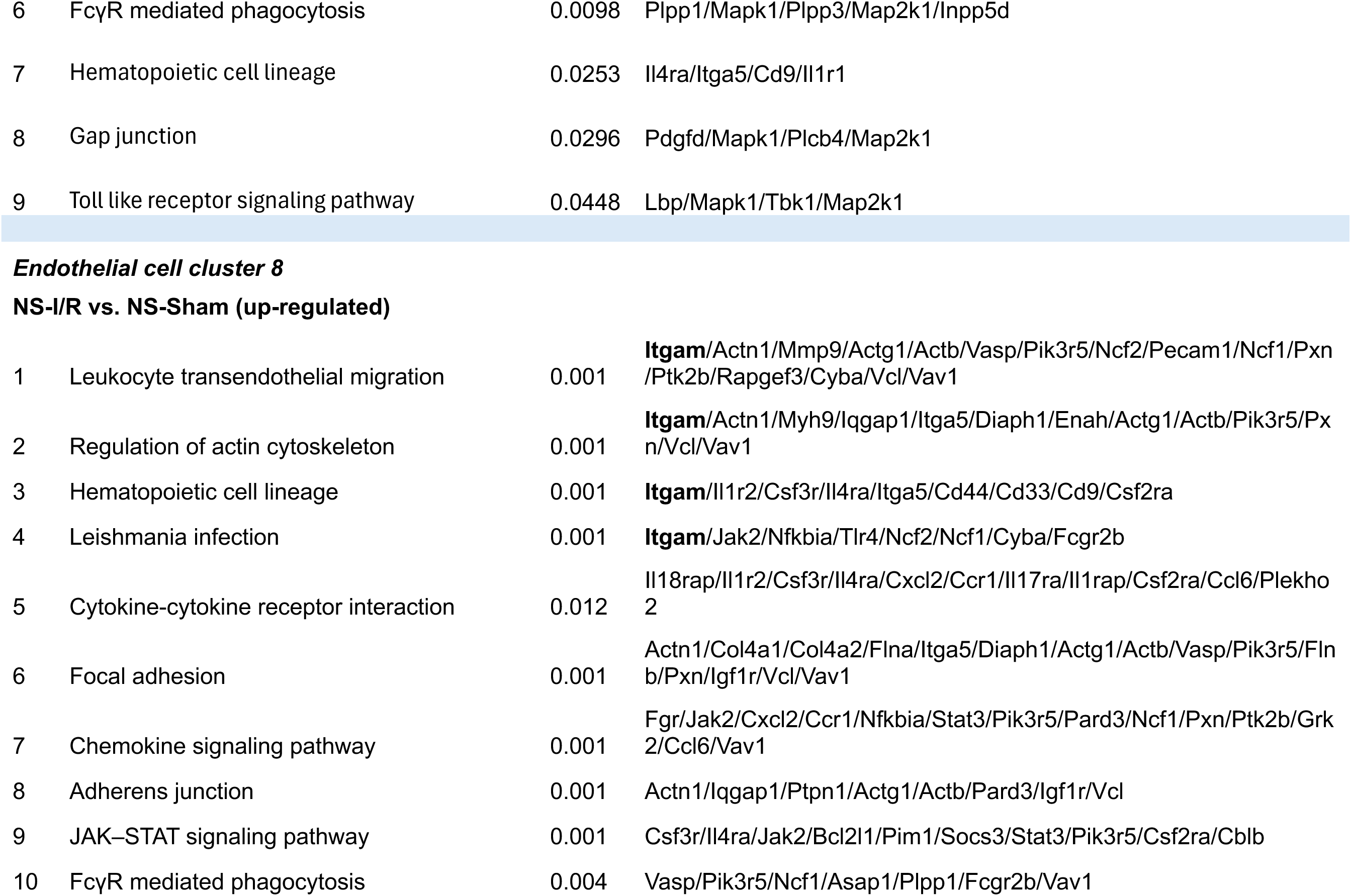
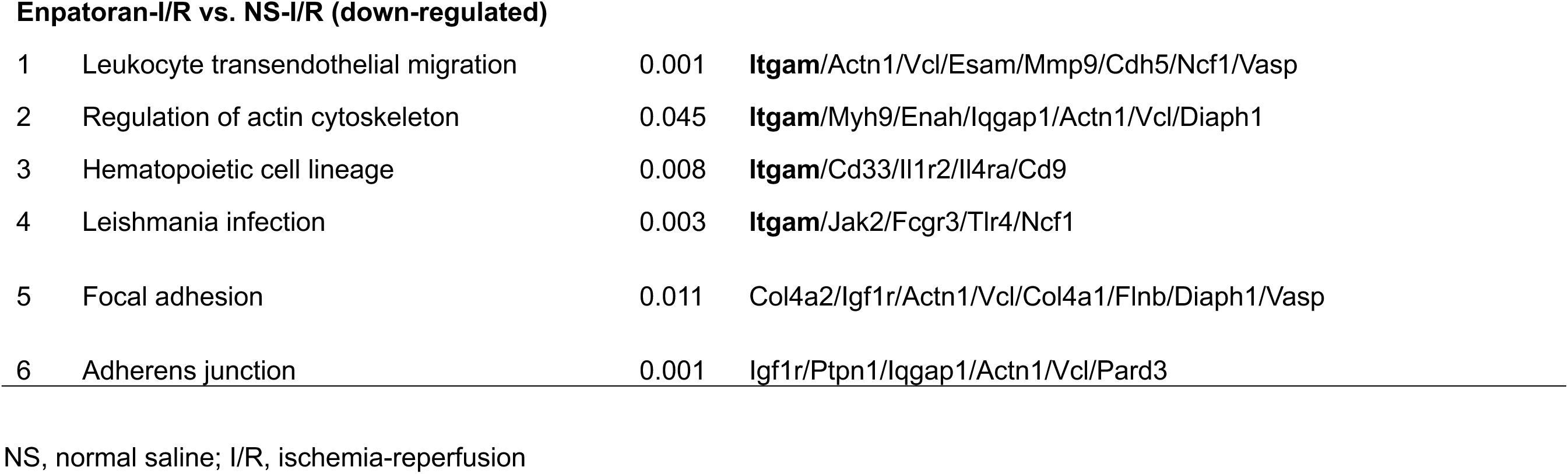
KEGG pathway analysis – enriched transcriptomic signatures of EC8, monocytes, and neutrophils.

**Table S9.**
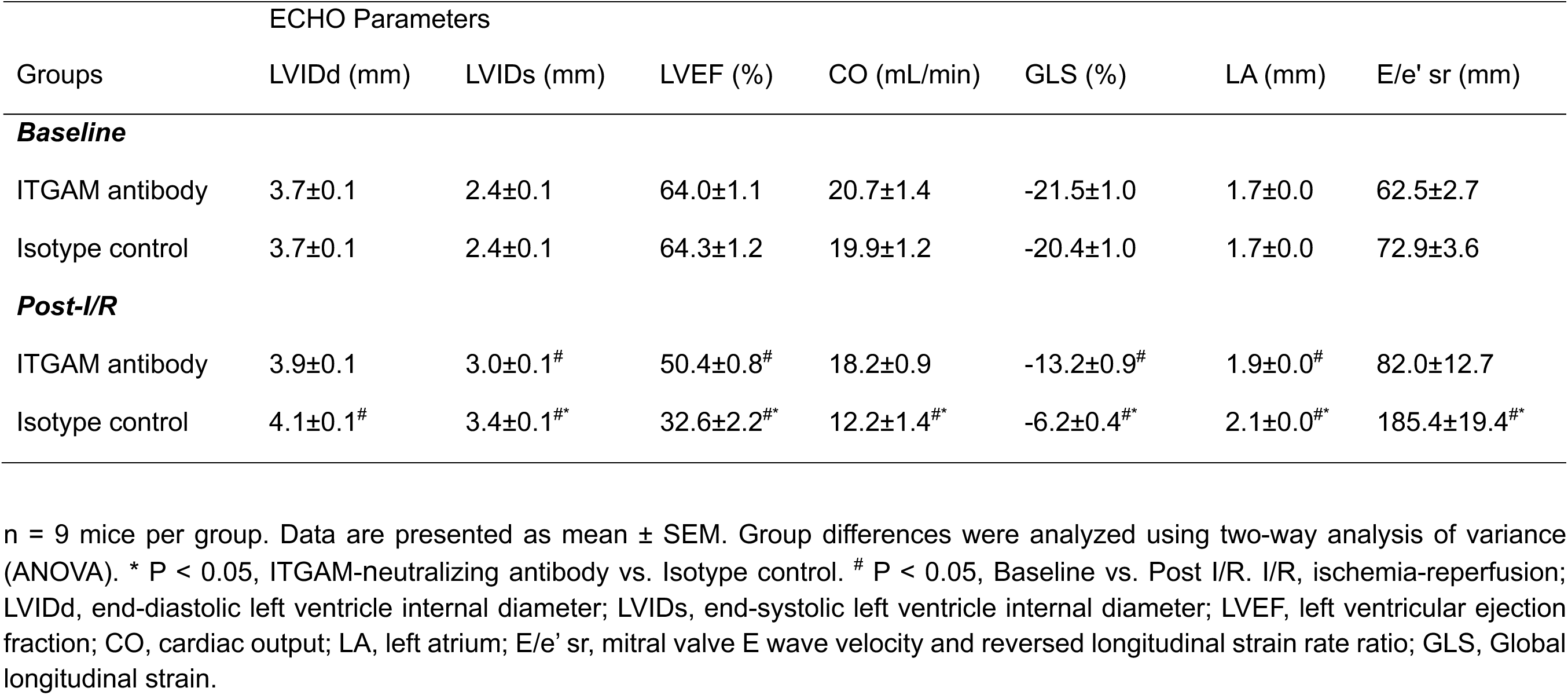
Detailed echocardiography data (baseline and post-I/R) in male mice treated with ITGAM and control antibody.

**Table S10.** Global marker genes.

## SUPPLEMENTAL FIGURES

**Figure S1.**
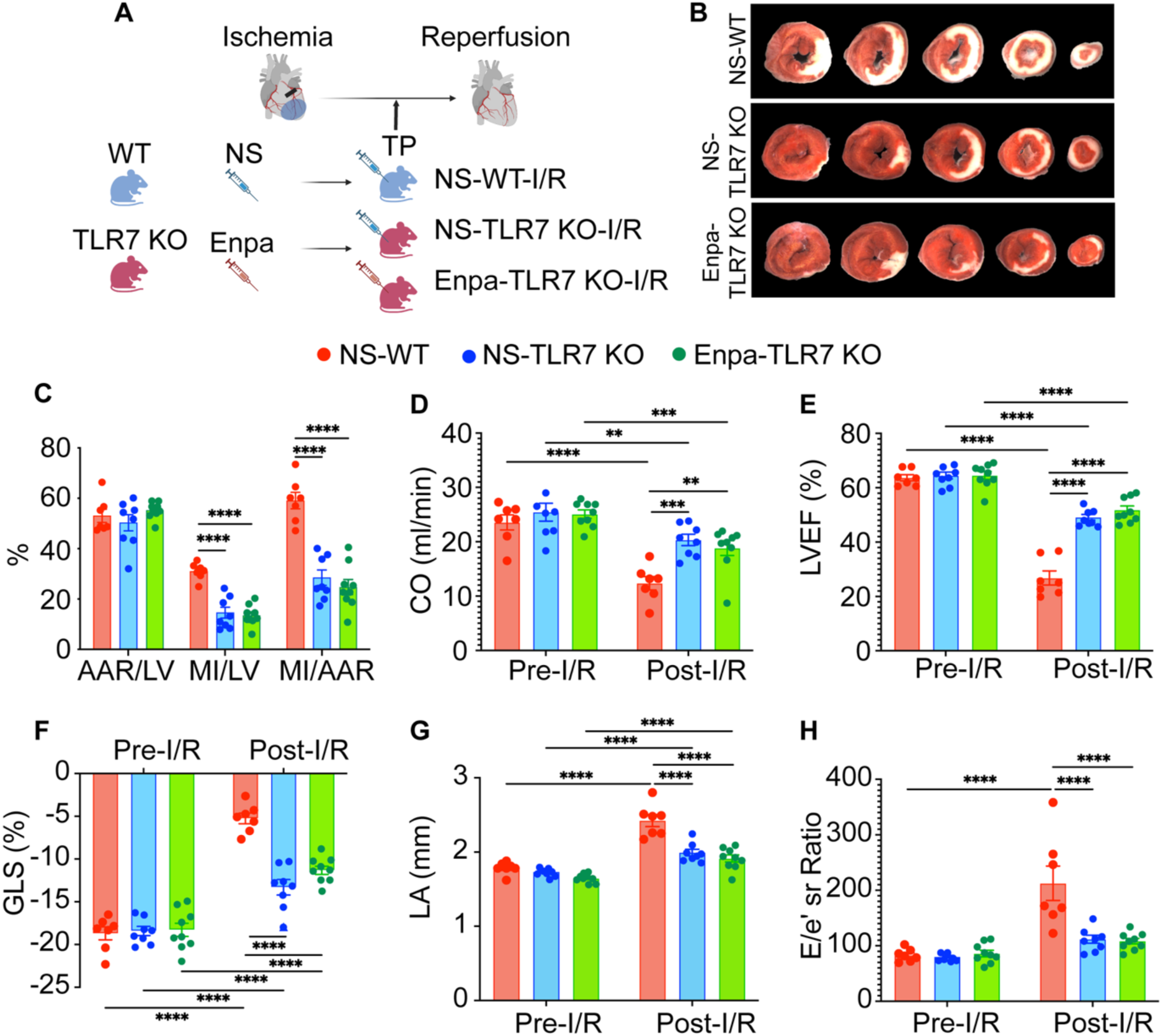
Enpatoran-induced cardioprotection in ischemic MI is TLR7-dependent. A, Study diagram for enpatoran administration (Treatment Protocol) in WT and TLR7 KO male mice subjected to myocardial I/R (n = 7-9 mice per group). **B**, Representative images of MI after triphenyl tetrazolium chloride (TTC) staining. **C,** AAR/LV, MI/LV, and MI/AAR 24h after I/R. **D-E**, Cardiac output (CO) and left ventricular ejection fraction (LVEF). **F,** Speckle tracking measurements with global longitudinal strain (GLS). **G,** Echocardiographic data depict LA diameter. **H**, Mitral valve E wave velocity, and reversed longitudinal strain rate ratio (E/e’ sr). Data shown are means ± SEM. Two-way ANOVA was used to compare groups for statistical significance. *P < 0.05, **P < 0.01, ***P < 0.001, ****P < 0.0001. NS, normal saline; Enpa, Enpatoran; WT, wild-type; TLR, toll-like receptor; KO, knockout; I/R, ischemia-reperfusion; AAR, area-at-risk; MI, myocardial infarction; LV, left ventricle; LA, left atrium; E/e’ sr, mitral valve E wave velocity and reversed longitudinal strain rate ratio.

**Figure S2.**
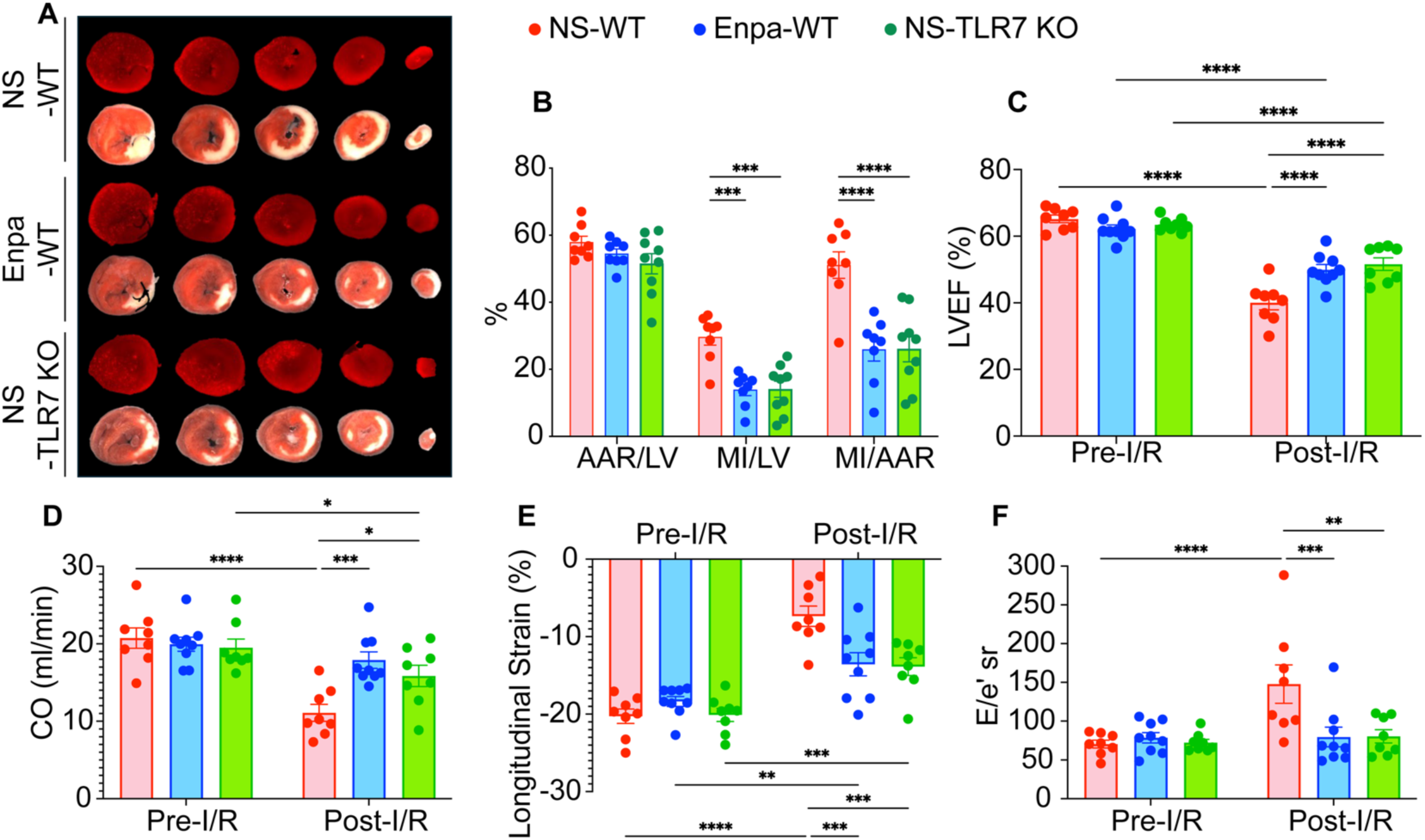
TLR7 KO and Enpatoran treatment reduce MI size and preserve cardiac function after I/R in female mice. A, Representative images of MI after triphenyl tetrazolium chloride (TTC) staining (n = 8-9 mice per group). **B,** AAR/LV, MI/LV, and MI/AAR 24h after I/R. **C-D**, Cardiac output (CO) and left ventricular ejection fraction (LVEF). **E**, Echocardiography speckle tracking imaging showing longitudinal strain. **F**, Echocardiographic data depict E wave to reversed longitudinal strain rate ratio (E/e’ sr). Data shown are means ± SEM. ANOVA was used to assess group differences for statistical significance. *P < 0.05, **P < 0.01, ***P < 0.001, ****P < 0.0001 among groups. NS, normal saline; Enpa, Enpatoran; WT, wild-type; KO, knockout; I/R, ischemia-reperfusion; AAR, area-at-risk; MI, myocardial infarction; LV, left ventricle. LVEF, left ventricular ejection fraction; E/e’ sr, mitral valve E wave velocity and reversed longitudinal strain rate ratio.

**Figure S3.**
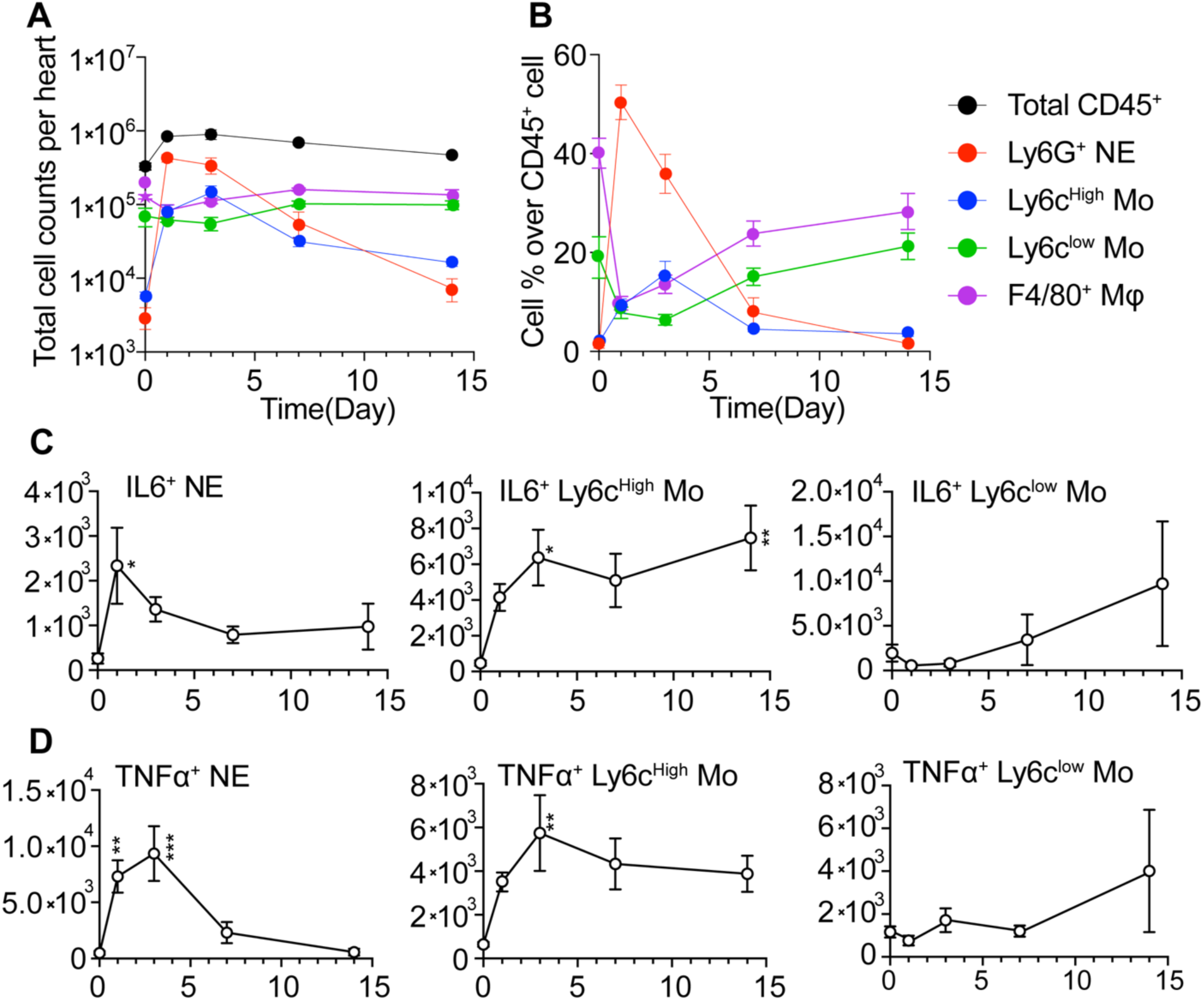
Temporal dynamics of cardiac immune cell infiltration and cytokine-expressing cells after I/R injury. A-B, Time-course analysis of leukocyte infiltration in mouse hearts following I/R injury. **C-D,** Temporal pattern of cytokine-expressing (IL6 or TNFα) immune cells in the heart after I/R. Symbols in the legend represent the following immune cell populations: black circle (Total CD45⁺), red circle (Ly6G⁺ neutrophils), blue circle (Ly6C^High^ monocytes), green circle (Ly6C^low^ monocytes), and purple circle (F4/80⁺ macrophages). n=5-6 mice per time point. Data are shown as mean ± SEM. One-way ANOVA with post-hoc Dunnett’s test was used to assess statistical differences across time points relative to baseline. *P < 0.05, **P < 0.01, ***P < 0.001 compared with time 0. I/R, ischemia-reperfusion; NE, neutrophil; Mo, monocyte; Mφ, macrophage; SEM, standard error of the mean.

**Figure S4.**
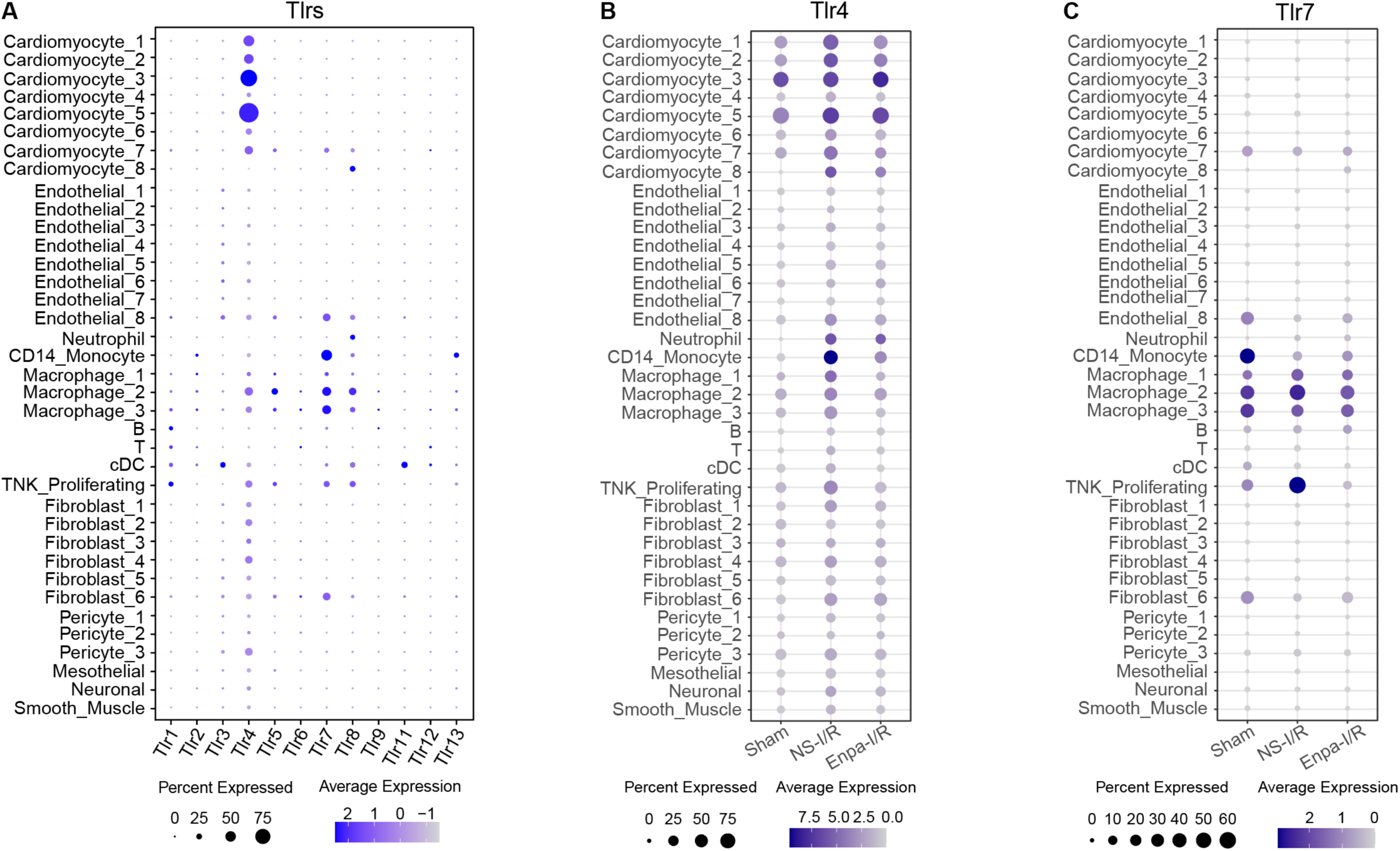
Cardiac cell type-specific TLR family gene expression and dynamic changes in response to I/R and enpatoran. A, Dot plot showing the expression patterns of TLR family genes across major cardiac cell types under Sham conditions. **B-C,** Dot plots illustrating the expression of *Tlr4* (**B**) and *Tlr7* (**C**) across various cardiac cell types under Sham, NS-I/R, and Enpa-I/R conditions. The color scale represents normalized average expression intensity. The dot size indicates the percentage of cells expressing each gene. *Tlr*, toll-like receptor; NS, normal saline; Enpa, enpatoran.

**Figure S5.**
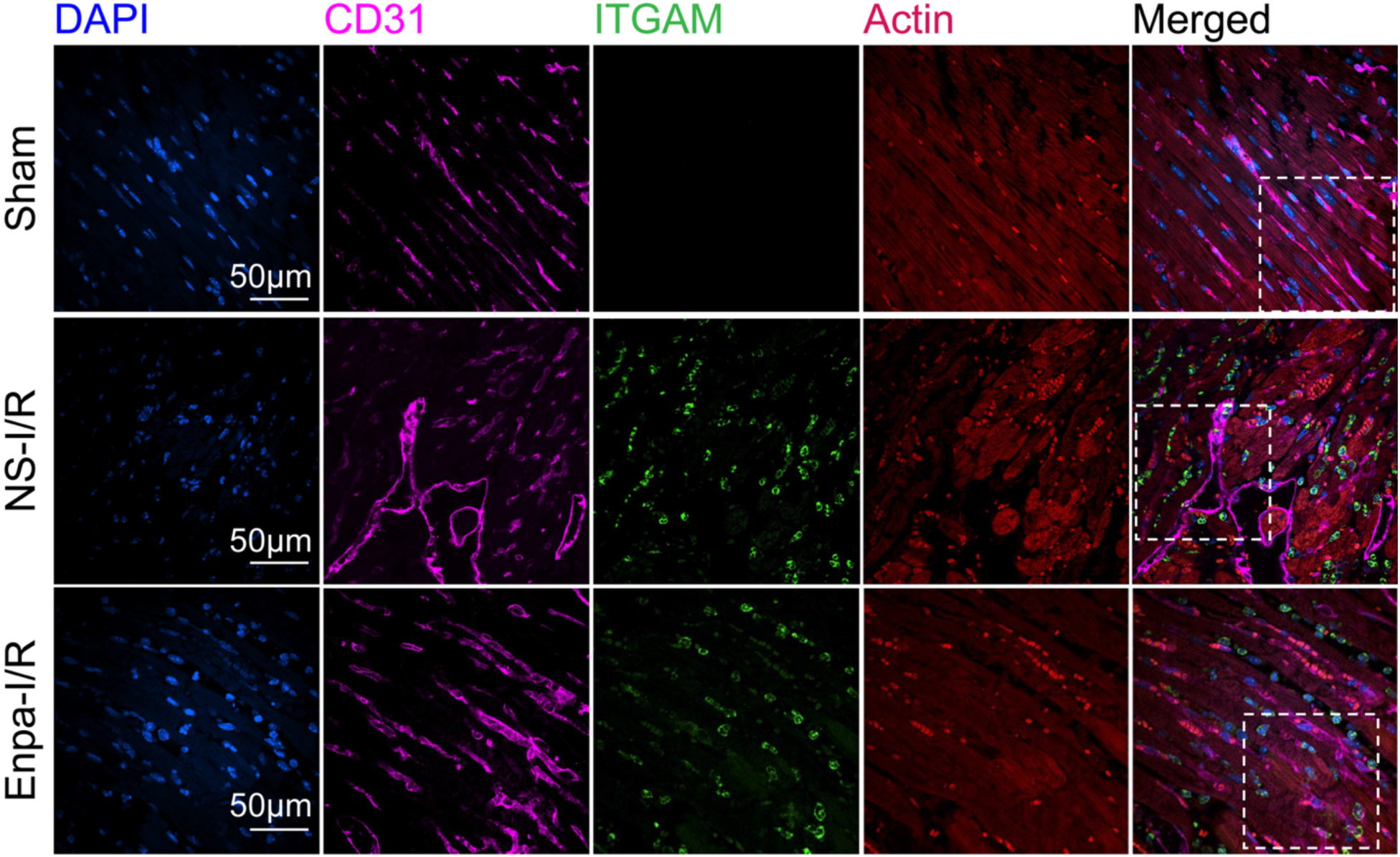
ITGAM expression in endothelial cells. Fluorescent imaging of DAPI (Blue), CD31 (magenta), ITGAM (green), and sarcomeric actin (red) on heart sections (60X). Dashed boxes indicate the regions shown at higher magnification in Fig. 3D - EC panel. NS, normal saline; Enpa, Enpatoran; I/R, ischemia-reperfusion; EC, endothelial cell; DAPI, 4′,6-diamidino-2-phenylindole.

**Figure S6.**
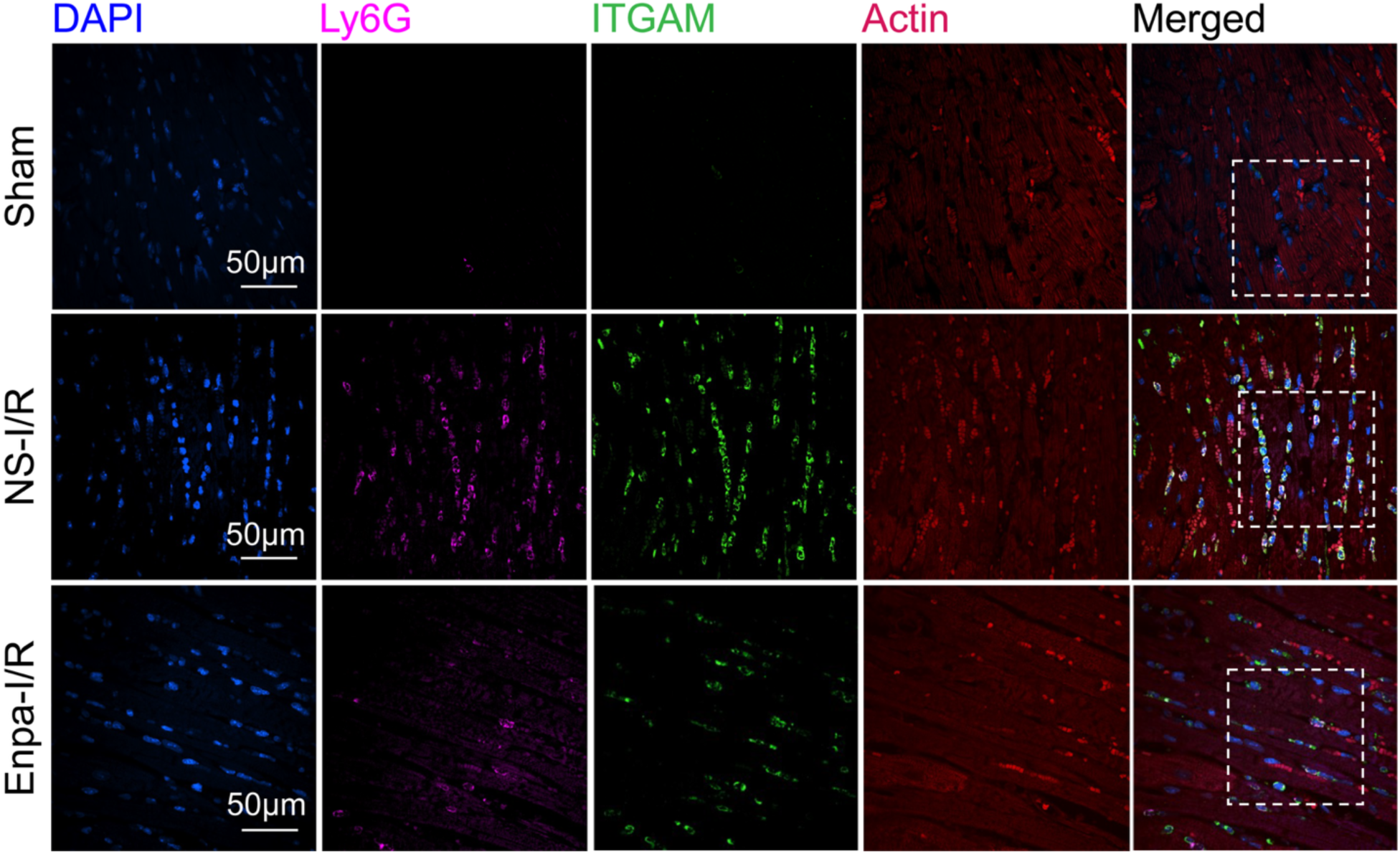
ITGAM expression in neutrophils. Fluorescent imaging of DAPI (Blue), Ly6G (magenta), ITGAM (green), and sarcomeric actin (red) on heart sections (60X). Dashed boxes indicate the regions shown at higher magnification in Fig. 3D - NE panel. NS, normal saline; Enpa, Enpatoran; I/R, ischemia-reperfusion; NE, neutrophil; DAPI, 4′,6-diamidino-2-phenylindole.

**Figure S7.**
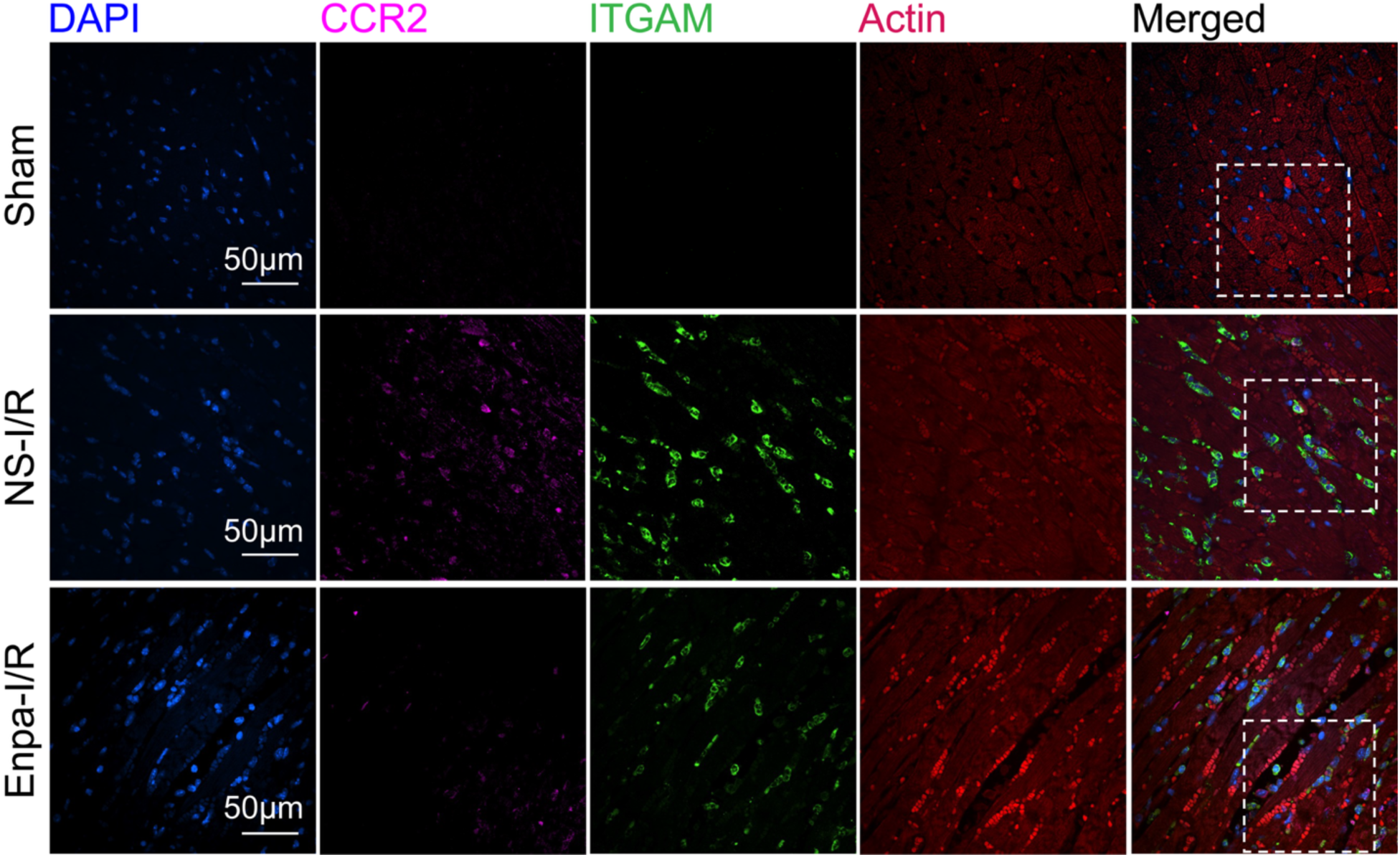
ITGAM expression in monocytes. Fluorescent imaging of DAPI (Blue), CCR2 (magenta), ITGAM (green), and sarcomeric actin (red) on heart sections (60X). Dashed boxes indicate the regions shown at higher magnification in Fig. 3D - Mo panel. NS, normal saline; Enpa, Enpatoran; I/R, ischemia-reperfusion; Mo, monocyte; DAPI, 4′,6-diamidino-2-phenylindole.

**Figure S8.**
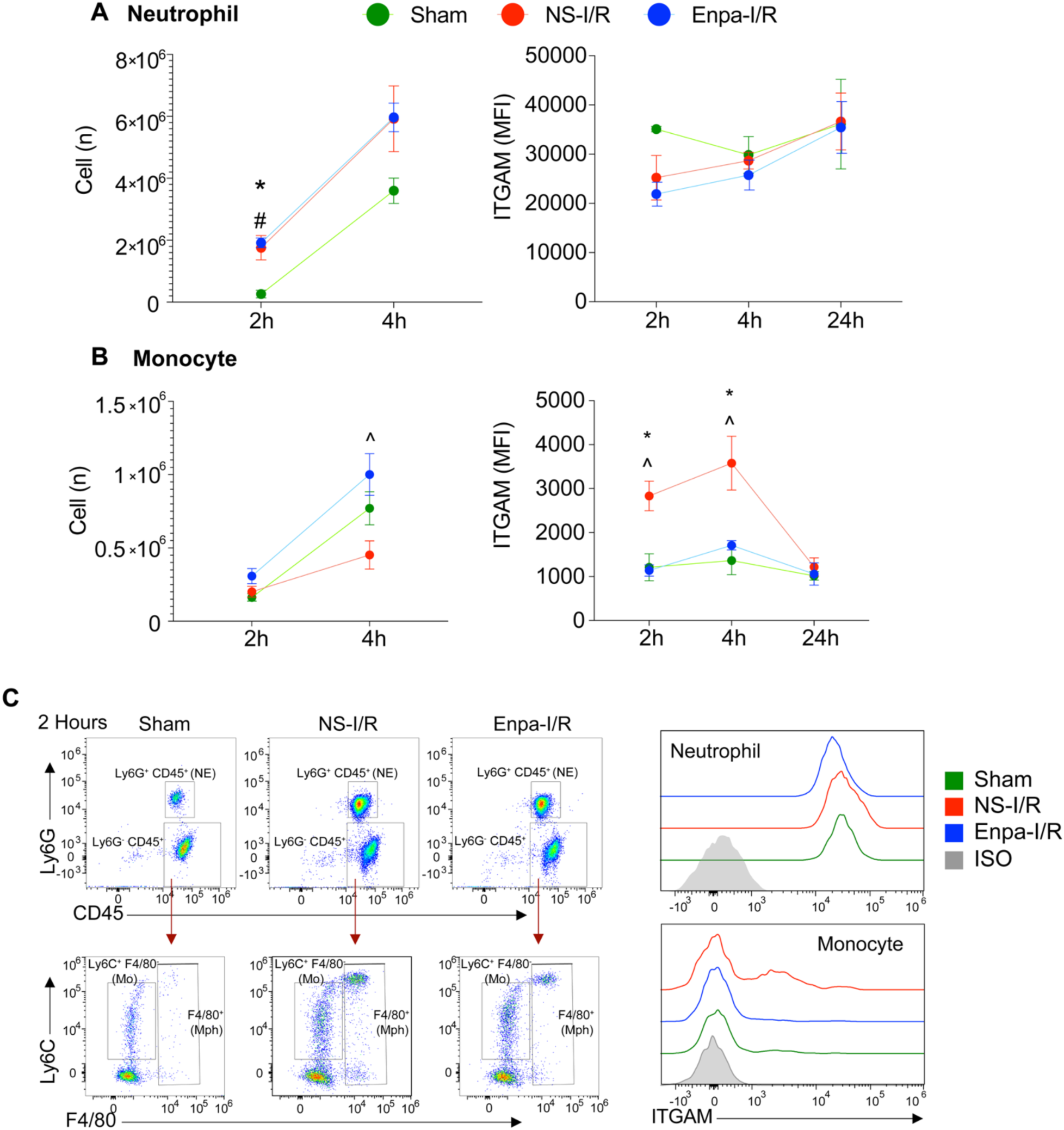
**Blood neutrophil and monocyte dynamics and ITGAM expression following myocardial I/R injury. A-B**, Flow cytometric quantification of circulating neutrophils (**A**) and monocytes (**B**) and ITGAM expression at 2h, 4h, and 24h after I/R injury, with Enpa or NS treatment. n=3-6 mice per time point. **C**, Representative gating strategy and histograms showing ITGAM expression in neutrophils and monocytes at 2h after I/R. Data are presented as mean ± SEM. One-way ANOVA was used for statistical comparisons among groups. *: Sham vs. NS-I/R, P < 0.05; #: Sham vs. Enpa-I/R, P < 0.05; ^: Enpa-I/R vs. NS-I/R, P < 0.05. NS, normal saline; Enpa, Enpatoran; I/R, ischemia-reperfusion; NE, neutrophil; Mo, monocyte; MFI, mean fluorescence intensity.

**Figure S9.**
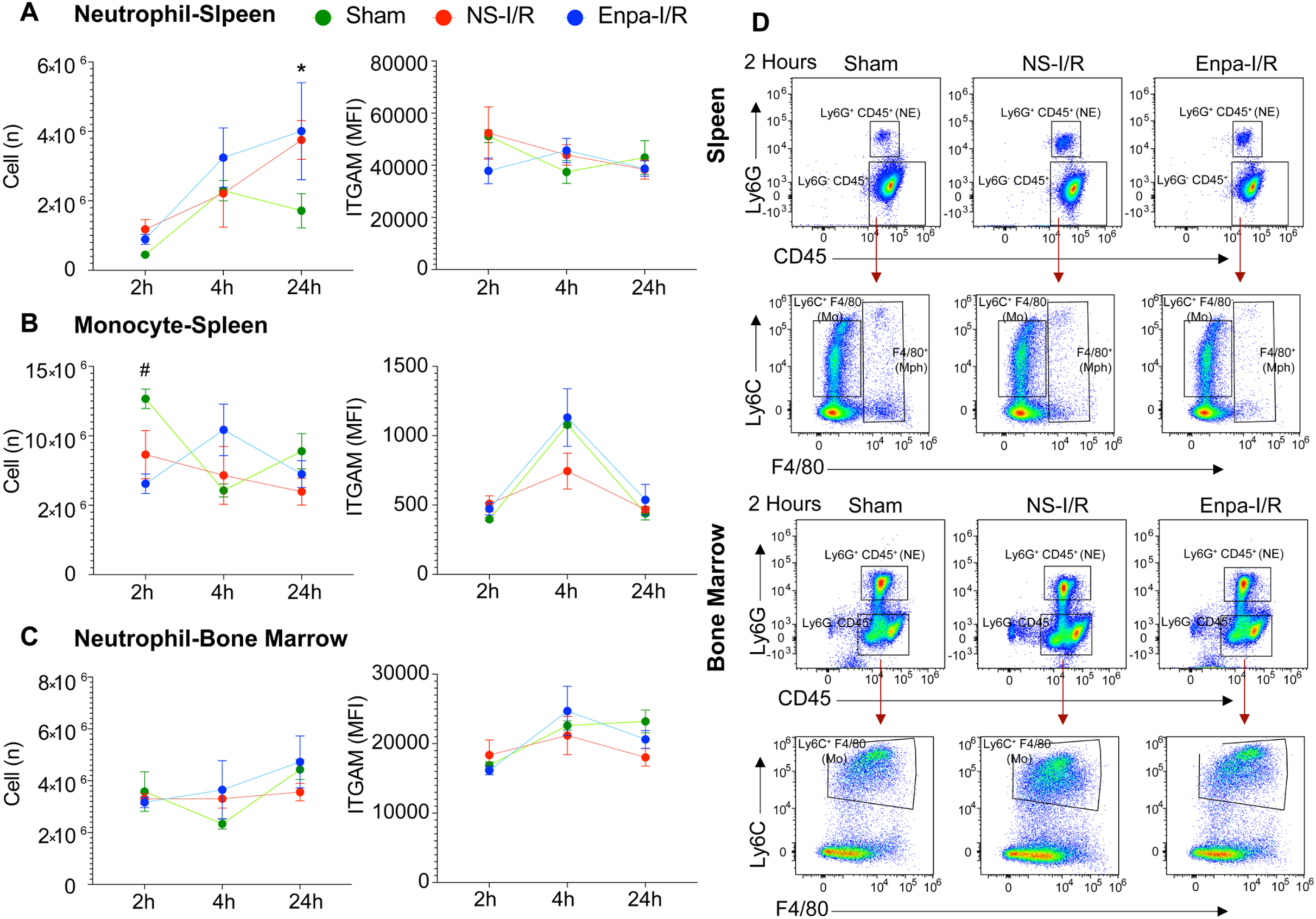
**Spleen and bone marrow neutrophil and monocyte dynamics and ITGAM expression following I/R injury. A-C**, Flow cytometric quantification of spleen and bone marrow neutrophils (**A and C**) and spleen monocytes (**B**), and ITGAM expression at 2h, 4h, and 24h after ischemia-reperfusion (I/R) injury, with or without Enpa treatment. n=3-6 mice per time point. **D**, Representative gating strategy and histograms showing ITGAM expression in neutrophils and monocytes at 2h after I/R. NS, normal saline; Enpa, Enpatoran; I/R, ischemia-reperfusion; NE, Neutrophil; Mo, Monocyte. Data are presented as means ± SEM. One-way ANOVA was used for statistical comparisons among groups. *: Sham vs. NS-I/R, P < 0.05; #: Sham vs. Enpa-I/R, P < 0.05; ^: Enpa-I/R vs. NS-I/R, P < 0.05. NS, normal saline; Enpa, Enpatoran; I/R, ischemia-reperfusion; NE, neutrophil; Mo, monocyte; MFI, mean fluorescence intensity.

**Figure S10.**
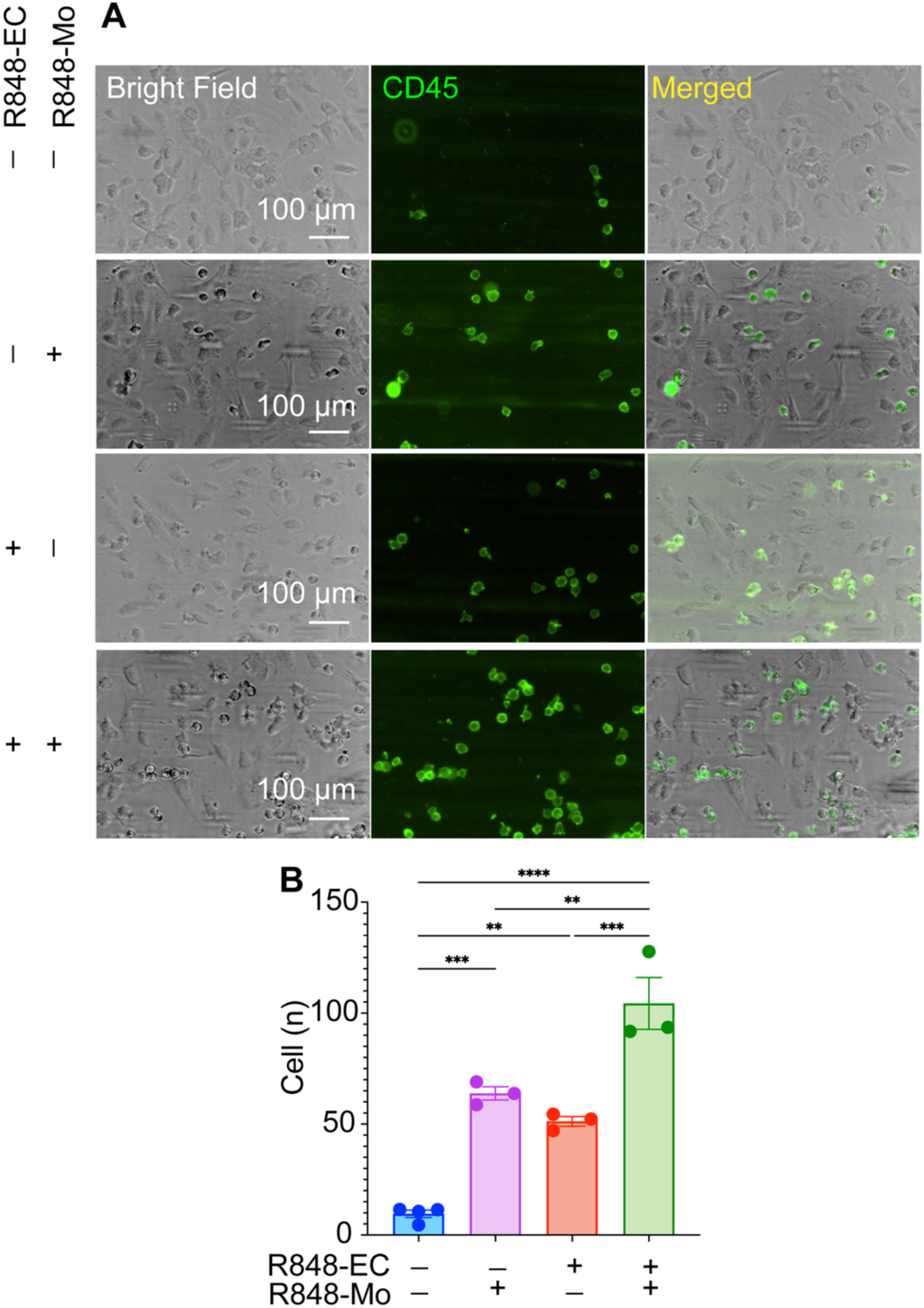
Microfluidic adhesion assay of THP-1 adhesion to HCAEC. Representative photomicrographs (**A**) and quantification (**B**) showing FITC anti-human CD45 antibody–labeled THP-1 cells (green) adhering to human coronary artery endothelial cells (HCAECs) within microfluidic channels. For each EC treatment condition, 3-4 corresponding channels were infused with THP-1 cells prepared under the corresponding treatment condition. Quantification of adherent monocytes was performed across 3-4 channels in one experiment. CD45⁺ immune cells are shown in green. Adherent CD45⁺ cells appear as discrete green puncta, while continuous green streaks reflect moving monocytes under flow conditions. nc, non-treated control; R, R848 (1μg/mL); Mo, monocytes; EC, endothelial cell.

**Figure S11.**
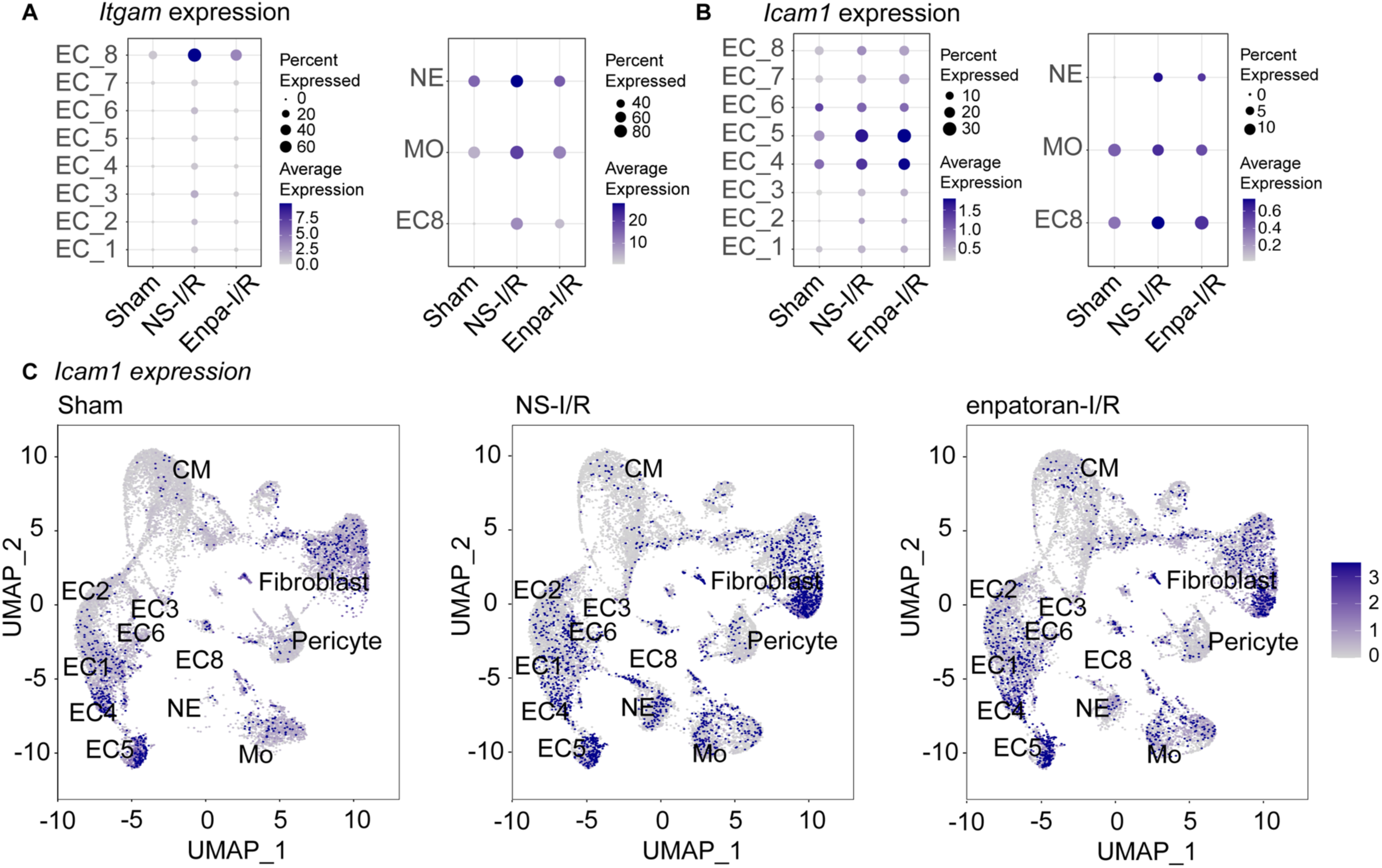
*Itgam* and *Icam1* expression level. A and B, Dot plots showing *Itgam* and *Icam1* expression levels and the proportion of *Itgam*-expressing cells across endothelial subtypes (EC1–EC8), neutrophils (NE), and monocytes (Mo). **C,** UMAP plots showing the spatial distribution and *Icam1* expression in snRNA-seq data from Sham, NS-I/R, and Enpa-I/R mouse hearts. The color intensity represents normalized expression levels of *Icam1* across cardiac cell clusters. *Icam1* expression was increased in NS-I/R hearts compared with Sham and was reduced with Enpa treatment. I/R, ischemia-reperfusion; NS, normal saline; Enpa, Enpatoran; EC, endothelial Cell; Mo, monocyte; NE, neutrophils; UMAP, uniform manifold approximation and projection.

**Figure S12.**
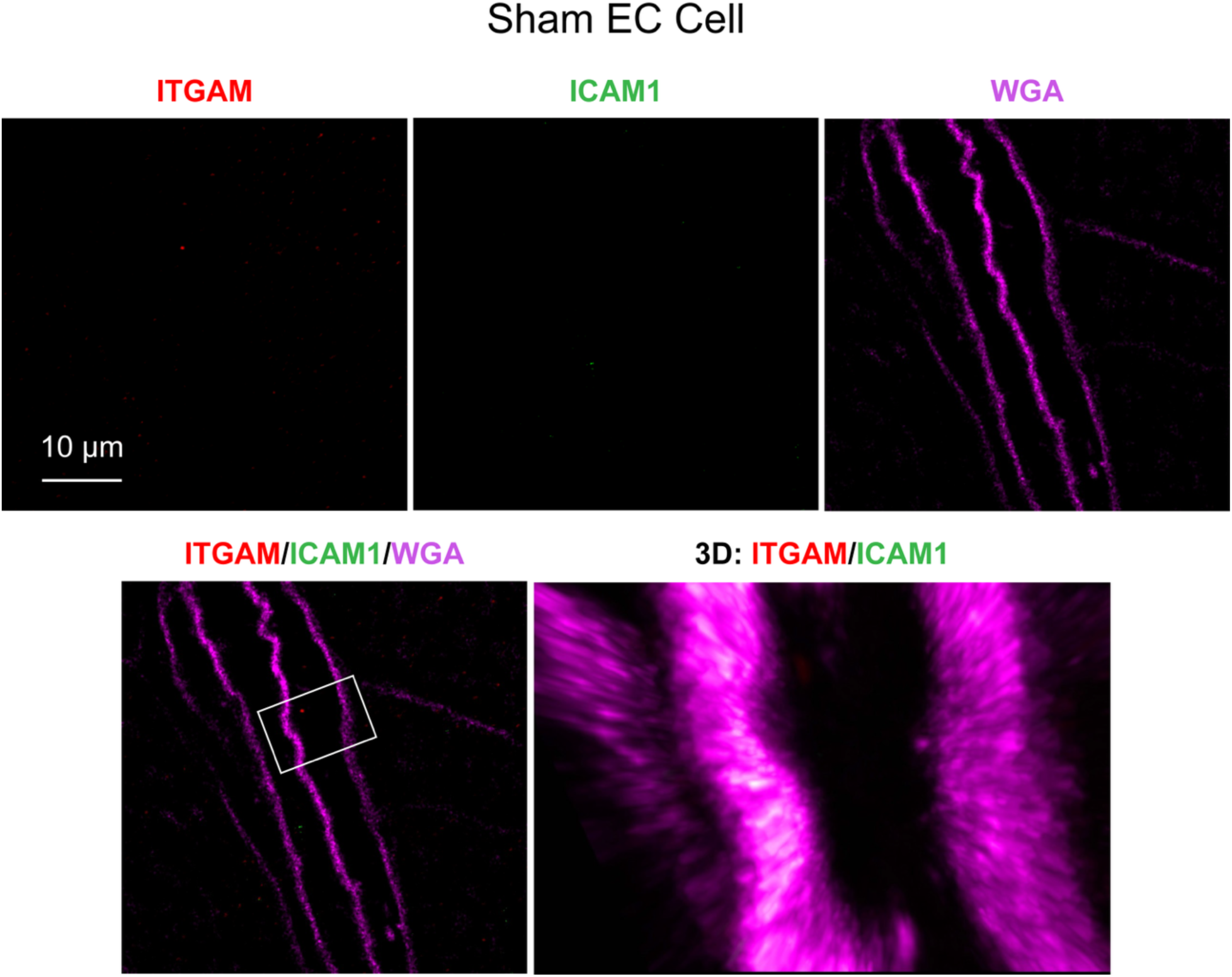
STED super-resolution imaging of endothelial cells in the normal heart. Cell membranes were imaged with Wheat Germ Agglutinin (WGA, magenta). There is a low level of ITGAM and ICAM-1 expression in the normal heart.

**Figure S13.**
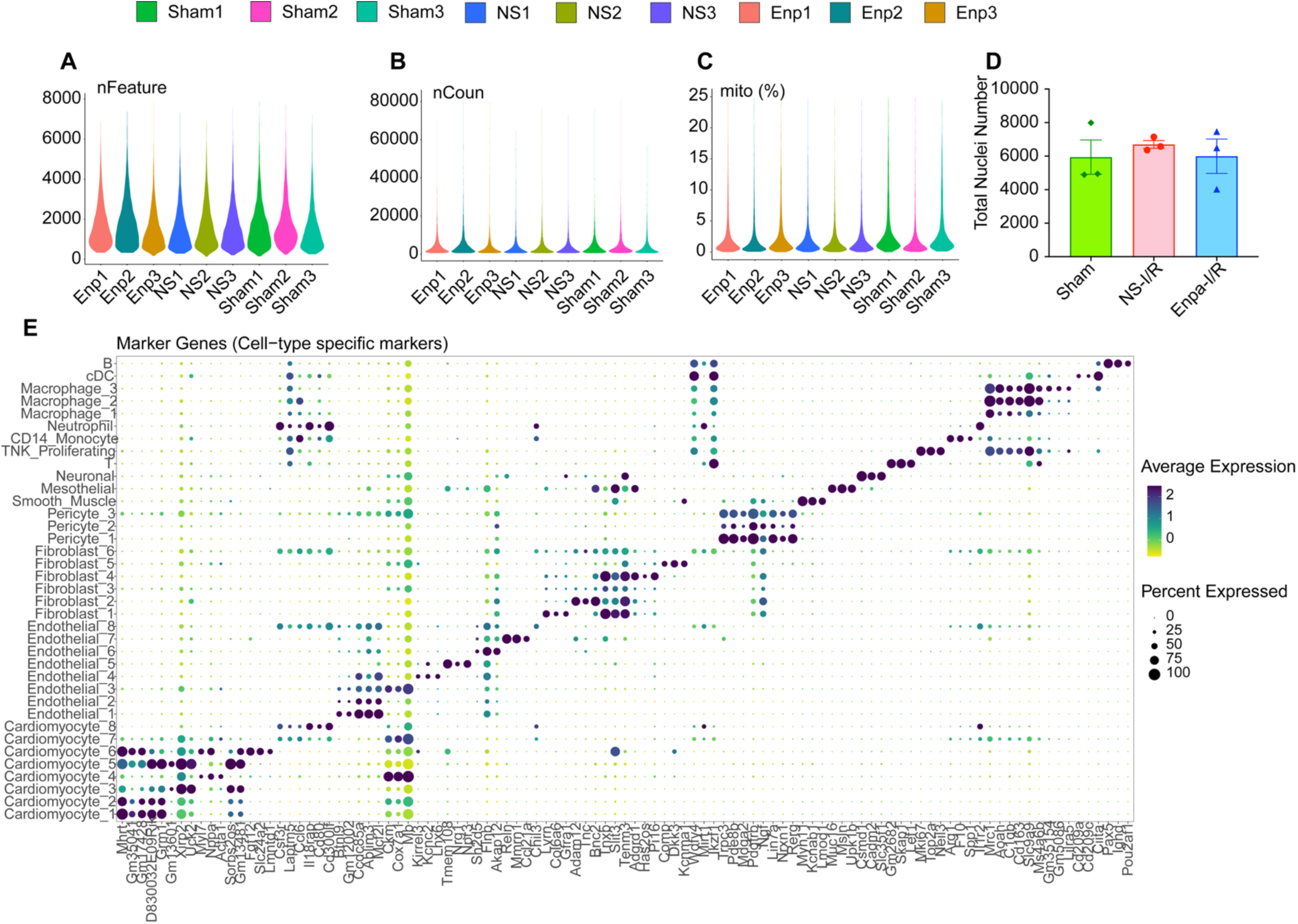
Detailed snRNA-seq analysis data. A-C, Quality control metrics for snRNA-seq across all samples, showing the number of detected genes per nucleus (*nFeature*, **A**), total RNA counts (*nCount*, **B**), and mitochondrial gene percentage (*mito%*, **C**); **D,** Quantification of the total number of nuclei analyzed per experimental condition (NS-Sham, NS-I/R, and Enpa-I/R); **E,** Cell cluster identification and validation based on cell-type-specific marker genes. The dot plot demonstrates the expression of canonical markers across distinct clusters, confirming clear separation. NS, normal saline; Enpa, Enpatoran; I/R, ischemia-reperfusion.

