## Supplemental Tables for "TLR7 inhibition limits cardiac ischemic injury by disrupting ITGAM-dependent immune–endothelial interaction"

**Table S1.** Upregulated plasma miRNAs after cardiac I/R injury

| | miRNA | Sequence | With motif |
|---|---|---|---|
| 1 | miR-29c-5p | UG**ACCGAUUUC**UCCUGGU**GUUC** | Y |
| 2 | miR-3061-3p | CUACC**UUUGAU**AGUCCACUGCC | Y |
| 3 | miR-741-3p | UGAGAG**AUGCCAUUCUAU**GUAGA | Y |
| 4 | miR-8116 | AAGGAUCCGGGGC**UAGUU**GGC | Y |
| 5 | miR-1843-5p | UAUGGAGGUCUCUGUCUGACU | N |
| 6 | miR-5113 | ACAGAGGAGGAGAGAGAUCCUGU | N |
| 7 | miR-8097 | AACAAGGAAAAUGUCCGGGCUG | N |

Mouse plasma small RNA-seq data of mice were analyzed using the R package DESeq2. I/R, n=9; Sham, n=6. The pro-inflammatory motifs were bolded and marked with underlines. I/R, ischemia-reperfusion.

**Table S2.** Detailed echocardiography data (baseline vs. Post-I/R) in four treatment protocols

| Groups | LVIDd (mm) | LVIDs (mm) | LVEF (%) | CO (mL/min) | GLS (%) | GRS (%) | LA (mm) | E/e' sr (mm) |
|---|---|---|---|---|---|---|---|---|
| ***Prevention Protocol (PP)*** | | | | | | | | |
| ***Baseline*** | | | | | | | | |
| NS | 3.9±0.1 | 2.6±0.1 | 62.3±2.4 | 21.1±1.2 | -18.1±0.9 | 28.8±1.4 | 1.9±0.0 | 76.5±6.9 |
| Enpatoran | 4.0±0.1 | 2.6±0.1 | 63.3±1.9 | 21.7±0.5 | -17.1±1.6 | 33.0±2.3 | 1.8±0.1 | 75.9±5.2 |
| ***Post-I/R*** | | | | | | | | |
| NS | 4.3±0.1$^{\#}$ | 3.6±0.1$^{\#}$ | 34.4±1.1$^{\#}$ | 11.1±0.5$^{\#}$ | -7.1±0.7$^{\#}$ | 13.9±1.5$^{\#}$ | 2.5±0.0$^{\#}$ | 174.7±15.2$^{\#}$ |
| Enpatoran | 4.1±0.1 | 3.1±0.1$^{*\#}$ | 48.9±1.8$^{*\#}$ | 15.2±0.7$^{*\#}$ | -12.9±1.6$^{*\#}$ | 22.6±2.6$^{*\#}$ | 2.1±0.1$^{*\#}$ | 112.9±8.9$^{*\#}$ |
| ***Treatment Protocol (TP)*** | | | | | | | | |
| ***Baseline*** | | | | | | | | |
| NS | 4.1±0.1 | 2.8±0.1 | 60.4±1.4 | 23.4±1.2 | -17.6±0.6 | 28.5±2.1 | 1.8±0.0 | 84.7±4.2 |
| Enpatoran | 4.2±0.1 | 2.9±0.1 | 60.3±1.2 | 24.4±0.9 | -18.7±0.4 | 29.7±1.8 | 1.7±0.0 | 75.4±3.9 |
| ***Post-I/R*** | | | | | | | | |
| NS | 4.2±0.1 | 3.6±0.1$^{\#}$ | 30.9±1.9$^{\#}$ | 11.3±0.9$^{\#}$ | -5.5±0.6$^{\#}$ | 12.7±2.0$^{\#}$ | 2.3±0.1$^{\#}$ | 239.6±24.8$^{\#}$ |
| Enpatoran | 4.0±0.1* | 2.9±0.1* | 53.4±1.4$^{*\#}$ | 17.9±1.1$^{*\#}$ | -15.3±1.4$^{*\#}$ | 27.6±1.6* | 1.9±0.0* | 103.3±9.9* |
| ***Early Rescue Protocol (ERP)*** | | | | | | | | |
| ***Baseline*** | | | | | | | | |
| NS | 4.0±0.1 | 2.7±0.1 | 61.7±1.0 | 24.1±1.1 | -20.4±0.9 | 33.9±2.8 | 1.7±0.0 | 71.8±5.1 |
| Enpatoran | 3.9±0.1 | 2.5±0.1 | 65.4±1.5 | 23.7±0.8 | -20.7±0.9 | 35.7±2.9 | 1.7±0.0 | 75.3±5.5 |

| | | | | | | | | |
|---|---|---|---|---|---|---|---|---|
| ***Post-I/R*** | | | | | | | | |
| NS | 4.0±0.2 | 3.5±0.2# | 27.5±2.1# | 8.5±0.7# | -5.9±0.8# | 10.8±0.8# | 2.5±0.7# | 211.8±36.1# |
| Enpatoran | 3.8±0.1 | 2.8±0.1* | 51.6±1.0*# | 14.8±0.8*# | -13.6±0.9*# | 25.9±3.2*# | 1.9±0.7*# | 94.9±10.0* |
| ***Late Rescue Protocol (LRP)*** | | | | | | | | |
| ***Baseline*** | | | | | | | | |
| NS | 4.0±0.1 | 2.7±0.1 | 63.7±1.0 | 25.8±1.5 | -20.4±1.4 | 30.3±2.7 | 1.7±0.0 | 78.3±6.9 |
| Enpatoran | 4.1±0.1 | 2.8±0.1 | 62.6±1.6 | 25.6±1.5 | -19.5±0.7 | 35.7±1.7 | 1.7±0.0 | 83.2±3.9 |
| ***Post-I/R*** | | | | | | | | |
| NS | 4.3±0.1 | 3.6±0.1# | 29.6±2.7# | 11.5±1.2# | -6.1±0.8# | 13.3±2.7# | 2.4±0.0# | 211.5±13.9# |
| Enpatoran | 4.0±0.2 | 2.9±0.2*# | 36.5±3.5# | 12.0±1.9# | -6.6±0.8# | 13.2±2.4# | 2.2±0.1*# | 176.2±26.0# |

n = 6 to 10 mice per group. Data are presented as mean ± SEM. Group differences were analyzed using two-way analysis of variance (ANOVA). * $P < 0.05$, NS vs. Enpatoran. # $P < 0.05$, Baseline vs. Post I/R. NS, normal saline; I/R, ischemia-reperfusion; PP, prevention protocol; TP, treatment protocol; ERP, early rescue protocol; LRP, late rescue protocol; LVIDd, end-diastolic left ventricle internal diameter; LVIDs, end-systolic left ventricle internal diameter; LVEF, left ventricular ejection fraction; CO, cardiac output; LA, left atrium; E/e’ sr, mitral valve E wave velocity and reversed longitudinal strain rate ratio; GLS, Global longitudinal strain; GRS, Global radial strain.

**Table S3.** Detailed baseline echocardiography data in control sham mice with the Prevention Protocol

| Groups | LVIDd (mm) | LVIDs (mm) | LVEF (%) | CO (mL/min) | GLS (%) | GRS (%) | LA (mm) | E/e' sr (mm) |
|---|---|---|---|---|---|---|---|---|
| ***Baseline*** | | | | | | | | |
| PP-Sham-NS | 4.0±0.5 | 2.7±0.0 | 60.5±0.6 | 21.7±0.6 | -18.4±1.2 | 33.0±2.1 | 1.7±0.0 | 92.6±4.8 |
| PP-Sham - Enpatoran | 3.9±0.6 | 2.6±0.0 | 62.5±0.9 | 22.1±1.0 | -19.7±0.8 | 36.0±1.1 | 1.7±0.0 | 84.4±4.2 |
| ***After Sham*** | | | | | | | | |
| PP-Sham-NS | 3.9±0.1 | 2.6±0.1 | 62.3±1.2 | 21.6±1.0 | -17.9±0.9 | 31.7±2.6 | 1.8±0.0 | 88.7±7.1 |
| PP-Sham - Enpatoran | 3.9±0.1 | 2.6±0.1 | 64.2±1.1 | 21.3±1.7 | -18.9±1.0 | 33.3±2.4 | 1.8±0.0 | 85.0±2.9 |

n = 6 per group. Data are presented as mean ± SEM. Group differences were analyzed using the *t*-test. NS, normal saline; PP, prevention protocol; LVIDd, end-diastolic left ventricle internal diameter; LVIDs, end-systolic left ventricle internal diameter; LVEF, left ventricular ejection fraction; CO, cardiac output; LA, left atrium; E/e’ sr, mitral valve E wave velocity and reversed longitudinal strain rate ratio; GLS, Global longitudinal strain; GRS, Global radial strain.

**Table S4.** Detailed echocardiography data (baseline and Post-I/R) in WT and TLR7 KO male mice

| Groups | LVIDd (mm) | LVIDs (mm) | LVEF (%) | CO (mL/min) | GLS (%) | LA (mm) | E/e' sr (mm) |
|---|---|---|---|---|---|---|---|
| ***Baseline*** | | | | | | | |
| NS-WT | 4.0±0.1 | 2.6±0.1 | 63.7±1.1 | 23.5±1.4 | -18.7±0.8 | 1.8±0.0 | 83.1±4.3 |
| NS-TLR7 KO | 4.1±0.1 | 2.6±0.1 | 64.5±1.3 | 25.4±1.6 | -18.4±0.5 | 1.7±0.0 | 80.3±2.2 |
| Enpatoran-TLR7 KO | 4.0±0.1 | 2.6±0.1 | 64.5±1.5 | 25.1±0.8 | -18.3±0.8 | 1.6±0.0 | 86.0±5.8 |
| ***Post-I/R*** | | | | | | | |
| NS-WT | $4.6±0.1^{\#}$ | $4.0±0.1^{\#}$ | $26.8±2.6^{\#}$ | $12.4±1.2^{\#}$ | $-5.2±0.6^{\#}$ | $2.4±0.1^{\#}$ | $212.7±30.9^{\#}$ |
| NS-TLR7 KO | $4.2±0.1^{*}$ | $3.2±0.1^{\#*}$ | $49.1±1.1^{\#*}$ | $20.3±1.1^{\#*}$ | $-13.3±0.9^{\#*}$ | $2.0±0.1^{\#*}$ | $111.9±7.3^{*}$ |
| Enpatoran-TLR7 KO | $4.0±0.1^{+}$ | $2.9±0.1^{\#+}$ | $51.8±1.5^{\#+}$ | $18.8±1.4^{\#+}$ | $-11.3±0.5^{\#+}$ | $1.9±0.1^{\#+}$ | $108.3±4.9^{+}$ |

n = 7-9 mice per group. Data are presented as mean ± SEM. Group differences were analyzed using two-way analysis of variance (ANOVA). * $P < 0.05$, NS-WT vs. NS-TLR7 KO. $^{+}$ $P < 0.05$, NS-WT vs. Enpatoran-TLR7 KO. $^{\#}$ $P < 0.05$, Baseline vs. Post I/R. NS, normal saline; I/R, ischemia-reperfusion; KO, knockout; LVIDd, end-diastolic left ventricle internal diameter; LVIDs, end-systolic left ventricle internal diameter; LVEF, left ventricular ejection fraction; CO, cardiac output; LA, left atrium; E/e’ sr, mitral valve E wave velocity and reversed longitudinal strain rate ratio; GLS, Global longitudinal strain.

**Table S5.** Detailed echocardiography data (baseline and Post-I/R) in female WT and KO mice

| Groups | LVIDd (mm) | LVIDs (mm) | LVEF (%) | CO (mL/min) | GLS (%) | LA (mm) | E/e' sr (mm) |
|---|---|---|---|---|---|---|---|
| ***Baseline*** | | | | | | | |
| NS-WT | 3.7±0.1 | 2.4±0.1 | 65.2±1.1 | 20.8±1.3 | -20.3±0.9 | 1.6±0.0 | 70.5±5.0 |
| NS-TLR7 KO | 3.6±0.1 | 2.4±0.1 | 63.5±0.7 | 19.5±1.1 | -20.1±0.8 | 1.6±0.0 | 72.3±4.2 |
| Enpatoran -WT | 3.7±0.1 | 2.5±0.1 | 62.2±1.2 | 20.0±0.9 | -18.4±0.6 | 1.7±0.0 | 78.6±6.7 |
| ***Post-I/R*** | | | | | | | |
| NS-WT | 3.8±0.1 | $3.0±0.1^{\#}$ | $40.0±2.1^{\#}$ | $11.1±1.1^{\#}$ | $-7.4±1.3^{\#}$ | $2.2±0.1^{\#}$ | $147.9±24.8^{\#}$ |
| NS-TLR7 KO | 3.8±0.1 | $2.8±0.1^{\#}$ | $51.6±1.9^{\#*}$ | $15.8±1.4^{\#*}$ | $-13.9±1.1^{\#*}$ | $1.9±0.0^{\#*}$ | $79.6±8.6^{*}$ |
| Enpatoran-WT | 4.0±0.1 | $3.0±0.1^{\#}$ | $49.9±1.6^{\#+}$ | $17.9±1.1^{+}$ | $-13.4±1.5^{\#+}$ | $1.9±0.0^{\#+}$ | $80.5±12.6^{+}$ |

Data are presented as mean ± SEM. n = 8-9 mice per group. Group differences were analyzed using two-way analysis of variance (ANOVA). * $P < 0.05$, NS-WT vs. NS-TLR7 KO. $^{+}$ $P < 0.05$, NS-WT vs. Enpatoran-TLR7 KO. $^{\#}$ $P < 0.05$, Baseline vs. Post I/R in each group. NS, normal saline; I/R, ischemia-reperfusion; LVIDd, end-diastolic left ventricle internal diameter; LVIDs, end-systolic left ventricle internal diameter; LVEF, left ventricular ejection fraction; CO, cardiac output; LA, left atrium; E/e' sr, mitral valve E wave velocity and reversed longitudinal strain rate ratio; GLS, Global longitudinal strain.

**Table S6.** Quality assessment of nucleus preparations for snRNA-seq

| | NS-Sham-1 | NS-Sham-2 | NS-Sham-3 | NS-I/R-1 | NS-I/R-2 | NS-I/R-3 | Enpatoran-I/R-1 | Enpatoran-I/R-2 | Enpatoran-I/R-3 |
|---|---|---|---|---|---|---|---|---|---|
| Quality Control Pass | Pass | Pass | Pass | Pass | Pass | Pass | Pass | Pass | Pass |
| Number of Nuclei | 4,960 | 8,198 | 4,975 | 6,497 | 6,707 | 7,228 | 7,571 | 4,079 | 6,551 |
| Valid Barcodes | 97.2 | 97.3 | 97.3 | 97.5 | 97.3 | 97.1 | 97.4 | 97.2 | 97.2 |
| Valid UMI | 100.0 | 99.9 | 100.0 | 100.0 | 99.9 | 99.9 | 99.9 | 99.9 | 100.0 |
| Number of Reads | 638,739,752 | 628,538,413 | 508,547,281 | 719,151,817 | 775,096,046 | 566,217,167 | 627,140,919 | 649,593,790 | 657,908,024 |
| Mean Reads per Cell | 128,778 | 76,670 | 102,221 | 110,690 | 115,565 | 78,337 | 82,835 | 159,253 | 100,429 |
| Median Genes per Cell | 1,418 | 1,416 | 1,208 | 1,221 | 1,321 | 1,328 | 1,416 | 1,550 | 1,198 |
| Sequencing Saturation | 91.1 | 86.9 | 91.2 | 90.9 | 90.9 | 88.9 | 88.5 | 92.7 | 91.2 |
| Q30 Bases in Barcode | 97.0 | 97.0 | 97.0 | 96.9 | 97.1 | 97.0 | 96.7 | 97.0 | 97.1 |
| Q30 Bases in RNA Read | 94.4 | 94.0 | 94.8 | 93.4 | 93.9 | 94.4 | 93.2 | 93.2 | 94.3 |
| Q30 Bases in UMI | 96.5 | 96.5 | 96.5 | 96.4 | 96.6 | 96.4 | 96.2 | 96.5 | 96.5 |

| | | | | | | | | | |
|---|---|---|---|---|---|---|---|---|---|
| Reads Mapped to Genome | 91.1 | 91.2 | 92.3 | 88.7 | 89.0 | 90.2 | 91.9 | 89.6 | 90.5 |
| Reads Mapped Confidently to Genome | 88.0 | 88.4 | 89.3 | 82.5 | 85.5 | 87.2 | 89.2 | 86.8 | 87.5 |
| Reads Mapped Confidently to Intergenic Regions | 7.8 | 7.2 | 7.1 | 8.9 | 8.0 | 6.6 | 7.0 | 6.2 | 7.1 |
| Reads Mapped Confidently to Intronic Regions | 41.9 | 51.4 | 42.6 | 42.7 | 45.6 | 45.8 | 46.8 | 47.4 | 44.4 |
| Reads Mapped Confidently to Exonic Regions | 38.2 | 29.7 | 39.6 | 31.0 | 31.9 | 34.9 | 35.4 | 33.2 | 36.0 |
| Reads Mapped Confidently to Transcriptome | 70.5 | 71.7 | 72.0 | 64.7 | 67.9 | 69.7 | 73.3 | 69.9 | 70.5 |
| Reads Mapped Antisense to Gene | 9.3 | 9.1 | 9.9 | 8.6 | 9.3 | 10.6 | 8.6 | 10.4 | 9.6 |
| Fraction Reads in Cells | 65.0 | 73.5 | 65.8 | 66.6 | 70.1 | 70.5 | 72.9 | 69.5 | 69.1 |
| Total Genes Detected | 24,786 | 25,312 | 24,282 | 25,691 | 25,878 | 25,087 | 25,202 | 24,244 | 24,750 |
| Median UMI Counts per Cell | 2,800 | 2,732 | 2,239 | 2,179 | 2,397 | 2,346 | 2,650 | 3,033 | 2,145 |

NS, normal saline; I/R, ischemia-reperfusion.

**Table S7.** Nucleus number and percentage (cluster nuclei / total nuclei x 100%) of each cell cluster

| Cell cluster | NS-Sham | NS-I/R | Enpatoran-I/R |
|---|---|---|---|
| Average nuclei number for analysis | 5939 | 6692 | 5994 |
| Cardiomyocytes | 1623.3 | 1345.3 | 1555 |
| Endothelial cells | 2083.3 | 2123.7 | 2002.3 |
| Immune cells | 375.3 | 1126* | 625.7# |
| Neutrophils | 2.0 | 505.7* | 158.7# |
| Monocytes | 5.7 | 318.3* | 171.3# |
| Macrophages | 278.7 | 222.0 | 229.7 |
| B cells | 39.3 | 31.3 | 22.7 |
| T cells | 24.3 | 34.7 | 19.3 |
| cDCs | 8.7 | 10.7 | 20.0 |
| Natural killer T cells | 16.7 | 3.3 | 4.0 |
| Fibroblasts | 1142.7 | 1385 | 1119 |
| Pericytes | 507.7 | 486.3 | 448.7 |
| Mesothelial cells | 40.3 | 41.7 | 31 |
| Neuronal cells | 32.7 | 35.7 | 31.7 |
| Smooth muscle cells | 134 | 148.3 | 180.7 |

n = 3 mice per group. NS, normal saline; I/R, ischemia-reperfusion; EC, endothelial cell. * $P < 0.05$, NS-Sham vs. NS-I/R. # $P < 0.05$, NS-I/R vs. Enpa-I/R.

**Table S8.** KEGG pathway analysis – enriched transcriptomic signatures in EC8, monocytes, and neutrophils

| | Enrichment Pathway | P value | Involved Genes |
|---|---|---|---|
| ***Monocyte*** | | | |
| **NS-I/R vs. NS-Sham (up-regulated)** | | | |
| 1 | Hematopoietic cell lineage | 0.001 | **Itgam**/Il4ra/Cd33/Il1r1/Cd38/Itga5/Il1r2 |
| 2 | Leishmania infection | 0.005 | **Itgam**/Tlr4/Fcgr3/Fcgr2b/Nfkbia |
| 3 | Cytokine cytokine receptor Interaction | 0.001 | Ccl2/Il4ra/Ccl6/Ccr1/Cxcl2/Cxcl3/Il18rap/Il1r1/Ccr5/Ccl9/Ccr2/Pf4/Il1r2/Tnfrsf11a/Il6st |
| 4 | Chemokine signaling pathway | 0.001 | Plcb1/Ccl2/Ccl6/Ccr1/Cxcl2/Cxcl3/Nfkbia/Ccr5/Ccl9/Ccr2/Pf4 |
| 5 | Endocytosis | 0.031 | Dab2/Iqsec1/Eea1/Ap2a2/Ccr5/Igf1r/Rab5c |
| 6 | Nod like receptor signaling pathway | 0.003 | Ccl2/Cxcl2/Cxcl3/Nfkbia/Pstpip1 |
| **Enpatoran-I/R vs. NS-I/R (down-regulated)** | | | |
| 1 | Hematopoietic cell lineage | 0.001 | **Itgam**/Il1r1/Cd33/Il1r2/Cd38/Cd9/Il4ra |
| 2 | Leishmania infection | 0.001 | **Itgam**/Tlr4/Nfkbia/Fcgr3/Irak1/Fcgr2b/Ncf2/Jak2 |
| 3 | Cytokine cytokine receptor interaction | 0.014 | Il18rap/Il1r1/Cxcl3/Il1r2/Cxcl2/Il6st/Il10ra/Ccr1/Il4ra/Acvr2a |
| 4 | Chemokine signaling pathway | 0.017 | Cxcl3/Cxcl2/Nfkbia/Plcb1/Ccr1/Braf/Map2k1/Jak2 |
| 5 | Toll-like receptor signaling pathway | 0.009 | Tlr4/Nfkbia/Cd80/Irak1/Tbk1/Map2k1 |
| 6 | Adherens junction | 0.048 | Tcf7l2/Ptpn1/Igf1r/Crebbp |
| ***Neutrophil*** | | | |
| **NS-I/R vs. NS-Sham (up-regulated)** | | | |

| | | | |
|---|---|---|---|
| 1 | Cytokine-cytokine receptor interaction | 0.029 | Il18rap/Il1r2 |
| 2 | Porphyrin and chlorophyll metabolism | 0.044 | Fth1 |
| **Enpatoran-I/R vs. NS-I/R (down-regulated)** | | | |
| 1 | Regulation of actin cytoskeleton | 0.005 | **Itgam**/Enah/Rock1/Iqgap2/Diaph1/Myh9/Msn |
| 2 | Leishmania infection | 0.005 | Fcgr2b/**Itgam**/Jak2/Fcgr3 |
| 3 | B cell receptor signaling pathway | 0.007 | Fcgr2b/Pik3ap1/Inpp5d/Nfat5 |
| 4 | Adherens junction | 0.039 | Ptpn1/Ptprj/Igf1r |
| 5 | Viral myocarditis | 0.034 | Cd80/Myh9/Dmd |
| ***Endothelial cell cluster 4*** | | | |
| **NS-I/R vs. NS-Sham (up-regulated)** | | | |
| 1 | Leukocyte transendothelial migration | 0.001 | Actn1/Actn4/Esam/Pecam1/Msn/Actb/Pxn/Rapgef4/Vcl/Rapgef3/Itgb1/Rac1/Actg1/Mapk12/Cdc42/F11r |
| 2 | Regulation of actin cytoskeleton | 0.001 | Actn1/Itga5/Rras2/Myh9/Actn4/Msn/Actb/Arpc1b/Pxn/Vcl/Nckap1/Itgb1/Rac1/Ppp1cb/Actg1/Cyfip1/Arhgef7/Enah/Crk/Iqgap1/Pak2/Arpc2/Cdc42/Pdgfd/Arhgef1 |
| 3 | Focal adhesion | 0.001 | Actn1/Col4a2/Col4a1/Itga5/Flnb/Lama5/Actn4/Actb/Parvb/Pxn/Xiap/Vcl/Itgb1/Rac1/Ppp1cb/Actg1/Flna/Crk/Vwf/Pak2/Cdc42/Pdgfd |
| 4 | Tight junction | 0.001 | Actn1/Pard3/Rras2/Epb41l2/Myh9/Actn4/Actb/Ybx3/Sptan1/Actg1/Ppp2ca/Vapa/Ppp2r2a/Cdc42/F11r |
| 5 | Endocytosis | 0.007 | Pard3/Ehd4/Cblb/Dab2/Asap1/Rab31/Ap2a2/Eea1/Cltc/Rabep1/Cdc42/Acap2/Smurf2/Rab11a |
| 6 | Adherens junction | 0.001 | Actn1/Pard3/Actn4/Actb/Tgfbr2/Vcl/Rac1/Actg1/Iqgap1/Cdc42 |
| 7 | Spliceosome | 0.017 | Snrnp70/Ddx39b/Tcerg1/Rbm25/Prpf38b/Srsf10/Hnrnpm/Tra2b/Ddx46/Hnrnpk |

| | | | |
|---|---|---|---|
| 8 | Epithelial cell signaling in helicobacter pylori infection | 0.001 | Nfkbia/Atp6v1h/Rac1/Mapk12/Ikbkb/Cdc42/F11r/Adam10/Nfkb1 |
| 9 | FcγR mediated phagocytosis | 0.009 | Plpp1/Arpc1b/Gab2/Rac1/Inpp5d/Asap1/Crk/Arpc2/Cdc42 |
| 10 | Apoptosis | 0.011 | Bcl2l1/Nfkbia/Xiap/Ikbkb/Irak3/Irak2/Nfkb1/Cflar |
| 11 | Dilated cardiomyopathy | 0.018 | Itga5/Tpm3/Actb/Dmd/Itgb1/Actg1/Tpm4/Adcy4 |
| 12 | Arrhythmogenic right ventricular cardiomyopathy arvc | 0.020 | Actn1/Itga5/Actn4/Actb/Dmd/Itgb1/Actg1 |
| **Enpatoran-I/R vs. NS-I/R (down-regulated)** | | | |
| 1 | Leukocyte transendothelial migration | 0.001 | Actn1/Mapk12/Actn4/Cdh5/Rac1/Pecam1/Rapgef4/Rapgef3/Esam |
| 2 | Spliceosome | 0.001 | Rbm25/Ddx39b/Snrnp70/Prpf38b/Hnrnpm/Prpf40a/Hnrnpc/Hnrnpk/Ddx46 |
| 3 | Tight junction | 0.001 | Pard3/Actn1/Epb41l2/Sptan1/Rras2/Actn4/Ybx3/Ppp2ca/Vapa |
| 4 | Regulation of actin cytoskeleton | 0.019 | Actn1/Pdgfd/Rras2/Itga5/Actn4/Rac1/Arpc1b/Arhgef7/Cyfip1 |
| 5 | Wnt signaling pathway | 0.007 | Plcb4/Prickle2/Fzd4/Ppp2ca/Tbl1xr1/Tcf7l2/Rac1/Senp2 |
| 6 | Jak stat signaling pathway | 0.021 | Il4ra/Bcl2l1/Spred1/Socs2/Socs5/Jak2/Osmr |
| 7 | Adherens junction | 0.013 | Pard3/Actn1/Actn4/Tcf7l2/Rac1 |
| 8 | Small cell lung cancer | 0.021 | Col4a1/Col4a2/Bcl2l1/Ikbkb/Max |
| 9 | Rig i like receptor signaling pathway | 0.034 | Ddx3x/Mapk12/Tbk1/Ikbkb |
| 10 | Epithelial cell signaling in helicobacter pylori infection | 0.043 | Mapk12/Atp6v1h/Ikbkb/Rac1 |
| ***Endothelial cell cluster 5*** | | | |

| | | | |
|---|---|---|---|
| **NS-I/R vs. NS-Sham (up-regulated)** | | | |
| 1 | Leukocyte transendothelial migration | 0.001 | Actn1/Msn/Actb/**Icam1**/Pxn/Afdn/Rock2/Rac1/Actg1/Cdh5/Itgb1/Ptk2/Esam/Rap1b/Rapgef3 |
| 2 | Regulation of actin cytoskeleton | 0.001 | Actn1/Itga5/Msn/Actb/Pxn/Itgav/Rock2/Arpc1b/Rras2/Wasf2/Map2k1/Rac1/Arhgef1/Baiap2/Actg1/Arpc2/Ppp1cb/Myh9/Itgb1/Ptk2/Pdgfd/Limk2/Abi2/Diaph1/Kras |
| 3 | Focal adhesion | 0.001 | Col4a1/Actn1/Col4a2/Itga5/Lama5/Parvb/Actb/Pxn/Itgav/Rock2/Rapgef1/Flnb/Vwf/Flna/Map2k1/Rac1/Actg1/Ppp1cb/Itgb1/Ptk2/Pdgfd/Rap1b/Capn2/Diaph1 |
| 4 | Tight junction | 0.001 | Actn1/Epb41l2/Ybx3/Actb/Afdn/Ppp2ca/Rras2/Cttn/Actg1/Myh9/Pard3/Tjp2/Kras/Vapa |
| 5 | Chemokine signaling pathway | 0.016 | Nfkbia/Pxn/Rock2/Plcb4/Map2k1/Rac1/Stat3/Gnb1/Grk5/Ptk2/Pard3/Rap1b/Kras |
| 6 | FcγR mediated phagocytosis | 0.001 | Plpp1/Gab2/Plpp3/Inpp5d/Arpc1b/Wasf2/Map2k1/Rac1/Arpc2/Dnm2/Limk2 |
| 7 | Adherens junction | 0.001 | Actn1/Actb/Afdn/Wasf2/Rac1/Baiap2/Actg1/Ptprj/Pard3/Tgfbr1 |
| 8 | Viral myocarditis | 0.016 | Actb/**Icam1**/Eif4g1/Rac1/Actg1/Eif4g3/Myh9 |
| 9 | Tgf beta signaling pathway | 0.014 | Id1/Tgfb1/Rock2/Ppp2ca/Smurf2/Smad1/Smurf1/Tgfbr1 |
| **Enpatoran-I/R vs. NS-I/R (down-regulated)** | | | |
| 1 | Leukocyte transendothelial migration | 0.2095 | Actn1/Afdn/Msn |
| 2 | Focal adhesion | 0.0014 | Col4a1/Itga5/Pdgfd/Col4a2/Actn1/Mapk1/Lama5/Map2k1/Flna |
| 3 | Regulation of actin cytoskeleton | 0.0022 | Itga5/Pdgfd/Actn1/Rras2/Mapk1/Baiap2/Cyfip1/Map2k1/Msn |
| 4 | Tight junction | 0.0021 | Ybx3/Actn1/Rras2/Afdn/Epb41l2/Amotl1/Pard3 |
| 5 | Adherens junction | 0.0032 | Actn1/Mapk1/Afdn/Baiap2/Pard3 |

| | | | |
|---|---|---|---|
| 6 | FcγR mediated phagocytosis | 0.0098 | Plpp1/Mapk1/Plpp3/Map2k1/Inpp5d |
| 7 | Hematopoietic cell lineage | 0.0253 | Il4ra/Itga5/Cd9/Il1r1 |
| 8 | Gap junction | 0.0296 | Pdgfd/Mapk1/Plcb4/Map2k1 |
| 9 | Toll like receptor signaling pathway | 0.0448 | Lbp/Mapk1/Tbk1/Map2k1 |
| ***Endothelial cell cluster 8*** | | | |
| **NS-I/R vs. NS-Sham (up-regulated)** | | | |
| 1 | Leukocyte transendothelial migration | 0.001 | **Itgam**/Actn1/Mmp9/Actg1/Actb/Vasp/Pik3r5/Ncf2/Pecam1/Ncf1/Pxn/Ptk2b/Rapgef3/Cyba/Vcl/Vav1 |
| 2 | Regulation of actin cytoskeleton | 0.001 | **Itgam**/Actn1/Myh9/Iqgap1/Itga5/Diaph1/Enah/Actg1/Actb/Pik3r5/Pxn/Vcl/Vav1 |
| 3 | Hematopoietic cell lineage | 0.001 | **Itgam**/Il1r2/Csf3r/Il4ra/Itga5/Cd44/Cd33/Cd9/Csf2ra |
| 4 | Leishmania infection | 0.001 | **Itgam**/Jak2/Nfkbia/Tlr4/Ncf2/Ncf1/Cyba/Fcgr2b |
| 5 | Cytokine-cytokine receptor interaction | 0.012 | Il18rap/Il1r2/Csf3r/Il4ra/Cxcl2/Ccr1/Il17ra/Il1rap/Csf2ra/Ccl6/Plekho2 |
| 6 | Focal adhesion | 0.001 | Actn1/Col4a1/Col4a2/Flna/Itga5/Diaph1/Actg1/Actb/Vasp/Pik3r5/Flnb/Pxn/Igf1r/Vcl/Vav1 |
| 7 | Chemokine signaling pathway | 0.001 | Fgr/Jak2/Cxcl2/Ccr1/Nfkbia/Stat3/Pik3r5/Pard3/Ncf1/Pxn/Ptk2b/Grk2/Ccl6/Vav1 |
| 8 | Adherens junction | 0.001 | Actn1/Iqgap1/Ptpn1/Actg1/Actb/Pard3/Igf1r/Vcl |
| 9 | JAK–STAT signaling pathway | 0.001 | Csf3r/Il4ra/Jak2/Bcl2l1/Pim1/Socs3/Stat3/Pik3r5/Csf2ra/Cblb |
| 10 | FcγR mediated phagocytosis | 0.004 | Vasp/Pik3r5/Ncf1/Asap1/Plpp1/Fcgr2b/Vav1 |

| | | | |
|---|---|---|---|
| **Enpatoran-I/R vs. NS-I/R (down-regulated)** | | | |
| 1 | Leukocyte transendothelial migration | 0.001 | **Itgam**/Actn1/Vcl/Esam/Mmp9/Cdh5/Ncf1/Vasp |
| 2 | Regulation of actin cytoskeleton | 0.045 | **Itgam**/Myh9/Enah/Iqgap1/Actn1/Vcl/Diaph1 |
| 3 | Hematopoietic cell lineage | 0.008 | **Itgam**/Cd33/Il1r2/Il4ra/Cd9 |
| 4 | Leishmania infection | 0.003 | **Itgam**/Jak2/Fcgr3/Tlr4/Ncf1 |
| 5 | Focal adhesion | 0.011 | Col4a2/Igf1r/Actn1/Vcl/Col4a1/Flnb/Diaph1/Vasp |
| 6 | Adherens junction | 0.001 | Igf1r/Ptpn1/Iqgap1/Actn1/Vcl/Pard3 |

NS, normal saline; I/R, ischemia-reperfusion

**Table S9.** Detailed echocardiography data (baseline and Post-I/R) in male mice treated with ITGAM and control antibody

| | ECHO Parameters | | | | | | |
|---|---|---|---|---|---|---|---|
| Groups | LVIDd (mm) | LVIDs (mm) | LVEF (%) | CO (mL/min) | GLS (%) | LA (mm) | E/e' sr (mm) |
| ***Baseline*** | | | | | | | |
| ITGAM antibody | 3.7±0.1 | 2.4±0.1 | 64.0±1.1 | 20.7±1.4 | -21.5±1.0 | 1.7±0.0 | 62.5±2.7 |
| Isotype control | 3.7±0.1 | 2.4±0.1 | 64.3±1.2 | 19.9±1.2 | -20.4±1.0 | 1.7±0.0 | 72.9±3.6 |
| ***Post-I/R*** | | | | | | | |
| ITGAM antibody | 3.9±0.1 | 3.0±0.1$^{\#}$ | 50.4±0.8$^{\#}$ | 18.2±0.9 | -13.2±0.9$^{\#}$ | 1.9±0.0$^{\#}$ | 82.0±12.7 |
| Isotype control | 4.1±0.1$^{\#}$ | 3.4±0.1$^{\#*}$ | 32.6±2.2$^{\#*}$ | 12.2±1.4$^{\#*}$ | -6.2±0.4$^{\#*}$ | 2.1±0.0$^{\#*}$ | 185.4±19.4$^{\#*}$ |

n = 9 mice per group. Data are presented as mean ± SEM. Group differences were analyzed using two-way analysis of variance (ANOVA). * $P < 0.05$, ITGAM-neutralizing antibody vs. Isotype control. $^{\#}$ $P < 0.05$, Baseline vs. Post I/R. I/R, ischemia-reperfusion; LVIDd, end-diastolic left ventricle internal diameter; LVIDs, end-systolic left ventricle internal diameter; LVEF, left ventricular ejection fraction; CO, cardiac output; LA, left atrium; E/e' sr, mitral valve E wave velocity and reversed longitudinal strain rate ratio; GLS, Global longitudinal strain.

**Table S10.** Global marker genes (separate file)
