## Supplemental Figures for "TLR7 inhibition limits cardiac ischemic injury by disrupting ITGAM-dependent immune–endothelial interaction"

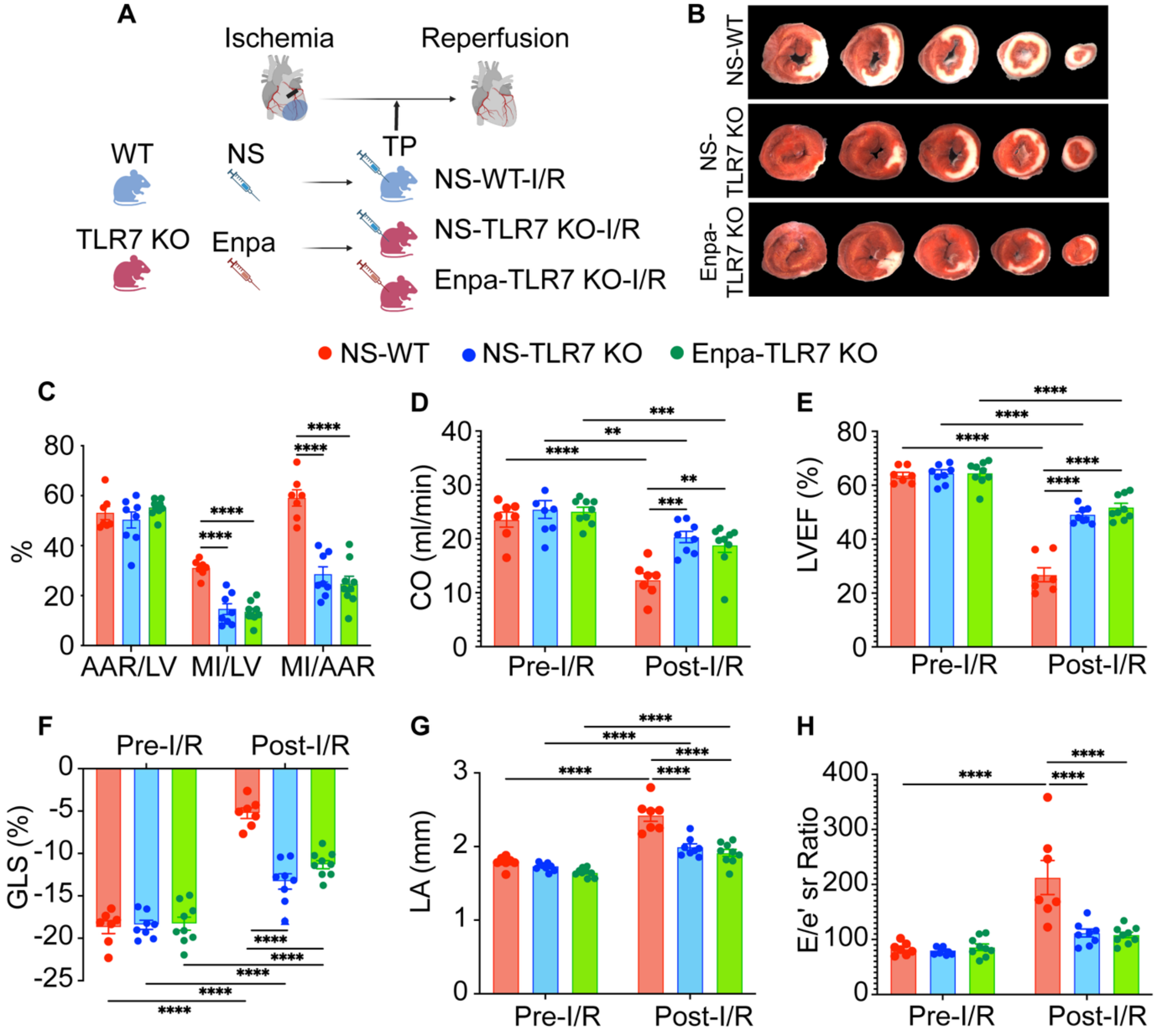


**Figure S1. Enpatoran-induced cardioprotection in ischemic MI is TLR7-dependent. A,** Study diagram for enpatoran administration (Treatment Protocol) in WT and TLR7 KO male mice subjected to myocardial I/R (n = 7-9 mice per group). **B**, Representative images of MI after triphenyl tetrazolium chloride (TTC) staining. **C,** AAR/LV, MI/LV, and MI/AAR 24h after I/R. **D-E**, Cardiac output (CO) and left ventricular ejection fraction (LVEF). **F,** Speckle tracking measurements with global longitudinal strain (GLS). **G,** Echocardiographic data depict LA diameter. **H**, Mitral valve E wave velocity, and reversed longitudinal strain rate ratio (E/e' sr). Data shown are means ± SEM. Two-way ANOVA was used to compare groups for statistical significance. $^*P < 0.05$, $^{**}P < 0.01$, $^{***}P < 0.001$, $^{****}P < 0.0001$. NS, normal saline; Enpa, Enpatoran; WT, wild-type; TLR, toll-like receptor; KO, knockout; I/R, ischemia-reperfusion; AAR, area-at-risk; MI, myocardial infarction; LV, left ventricle; LA, left atrium; E/e' sr, mitral valve E wave velocity and reversed longitudinal strain rate ratio.

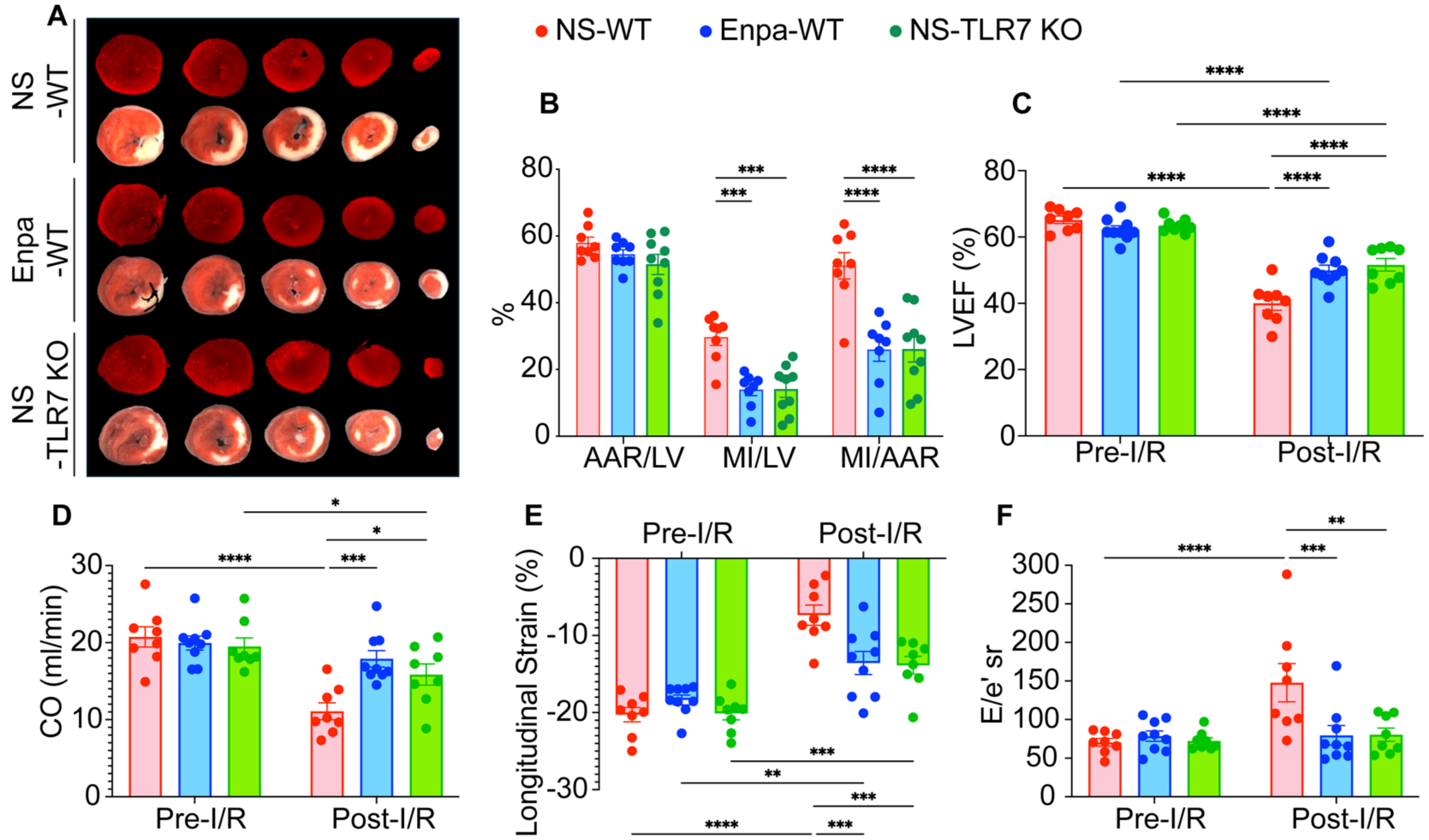


**Figure S2. TLR7 KO and Enpatoran treatment reduce MI size and preserve cardiac function after I/R in female mice. A,** Representative images of MI after triphenyl tetrazolium chloride (TTC) staining (n = 8-9 mice per group). **B,** AAR/LV, MI/LV, and MI/AAR 24h after I/R. **C-D**, Cardiac output (CO) and left ventricular ejection fraction (LVEF). **E**, Echocardiography speckle tracking imaging showing longitudinal strain. **F**, Echocardiographic data depict E wave to reversed longitudinal strain rate ratio (E/e' sr). Data shown are means ± SEM. ANOVA was used to assess group differences for statistical significance. *P < 0.05, **P < 0.01, ***P < 0.001, ****P < 0.0001 among groups. NS, normal saline; Enpa, Enpatoran; WT, wild-type; KO, knockout; I/R, ischemia-reperfusion; AAR, area-at-risk; MI, myocardial infarction; LV, left ventricle. LVEF, left ventricular ejection fraction; E/e' sr, mitral valve E wave velocity and reversed longitudinal strain rate ratio.

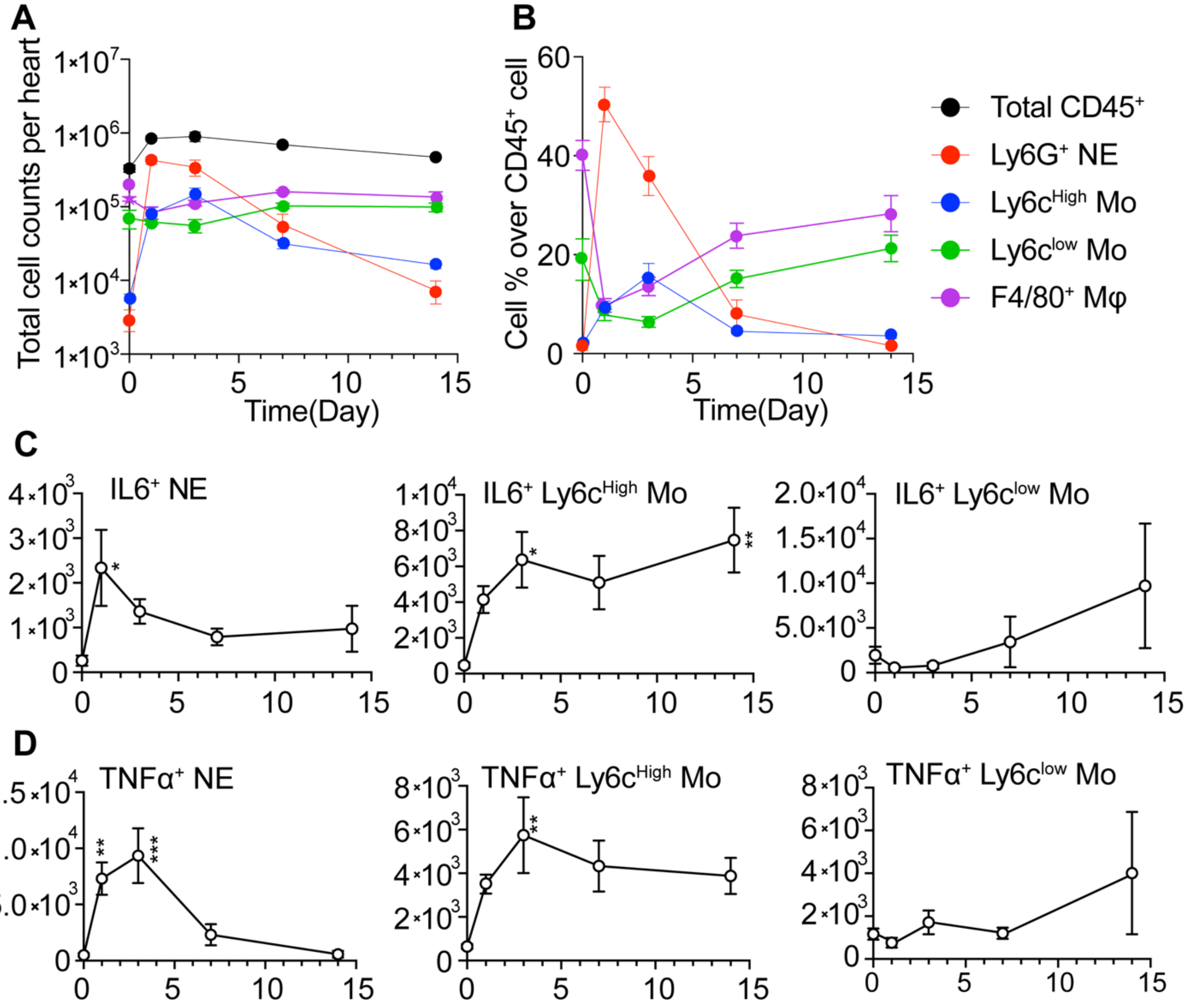


**Figure S3. Temporal dynamics of cardiac immune cell infiltration and cytokine-expressing cells after I/R injury. A-B,** Time-course analysis of leukocyte infiltration in mouse hearts following I/R injury. **C-D,** Temporal pattern of cytokine-expressing (IL6 or TNFα) immune cells in the heart after I/R. Symbols in the legend represent the following immune cell populations: black circle (Total $CD45^+$), red circle ($Ly6G^+$ neutrophils), blue circle ($Ly6C^{High}$ monocytes), green circle ($Ly6C^{low}$ monocytes), and purple circle ($F4/80^+$ macrophages). n=5-6 mice per time point. Data are shown as mean ± SEM. One-way ANOVA with post-hoc Dunnett's test was used to assess statistical differences across time points relative to baseline. *P < 0.05, **P < 0.01, ***P < 0.001 compared with time 0. I/R, ischemia-reperfusion; NE, neutrophil; Mo, monocyte; Mφ, macrophage; SEM, standard error of the mean.

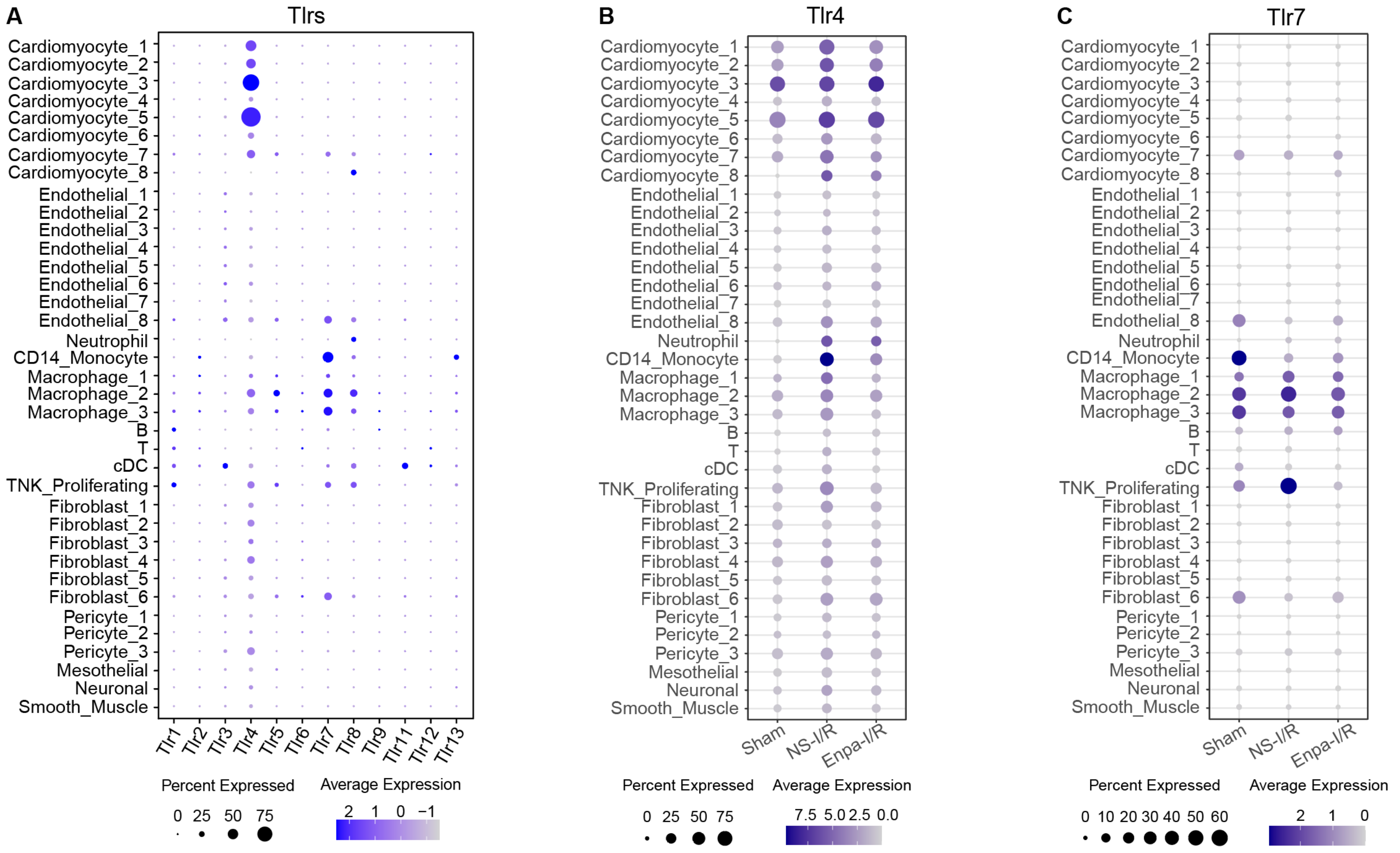


**Figure S4. Cardiac cell type-specific TLR family gene expression and dynamic changes in response to I/R and enpatoran. A,** Dot plot showing the expression patterns of TLR family genes across major cardiac cell types under Sham conditions. **B-C,** Dot plots illustrating the expression of *Tlr4* (**B**) and *Tlr7* (**C**) across various cardiac cell types under Sham, NS-I/R, and Enpa-I/R conditions. The color scale represents normalized average expression intensity. The dot size indicates the percentage of cells expressing each gene. *Tlr*, toll-like receptor; NS, normal saline; Enpa, enpatoran.

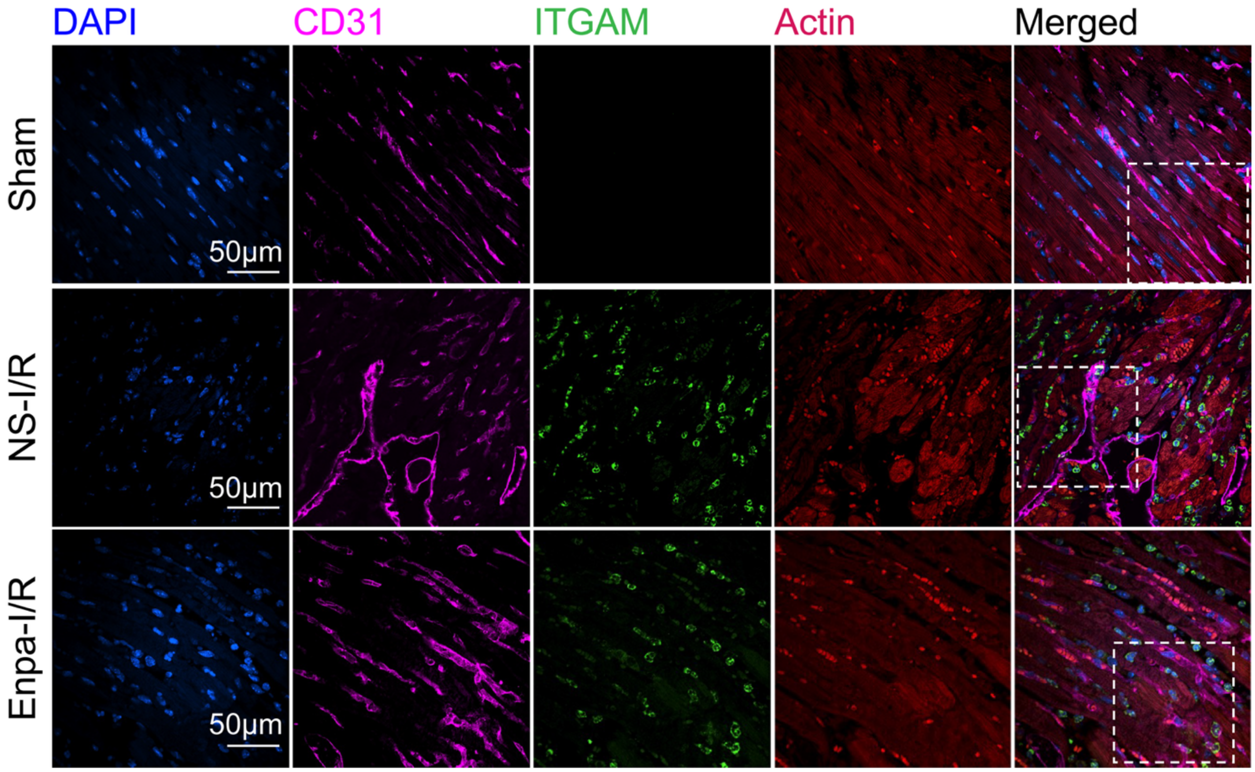


**Figure S5. ITGAM expression in endothelial cells.** Fluorescent imaging of DAPI (Blue), CD31 (magenta), ITGAM (green), and sarcomeric actin (red) on heart sections (60X). Dashed boxes indicate the regions shown at higher magnification in Fig. 3D - EC panel. NS, normal saline; Enpa, Enpatoran; I/R, ischemia-reperfusion; EC, endothelial cell; DAPI, 4′,6-diamidino-2-phenylindole.

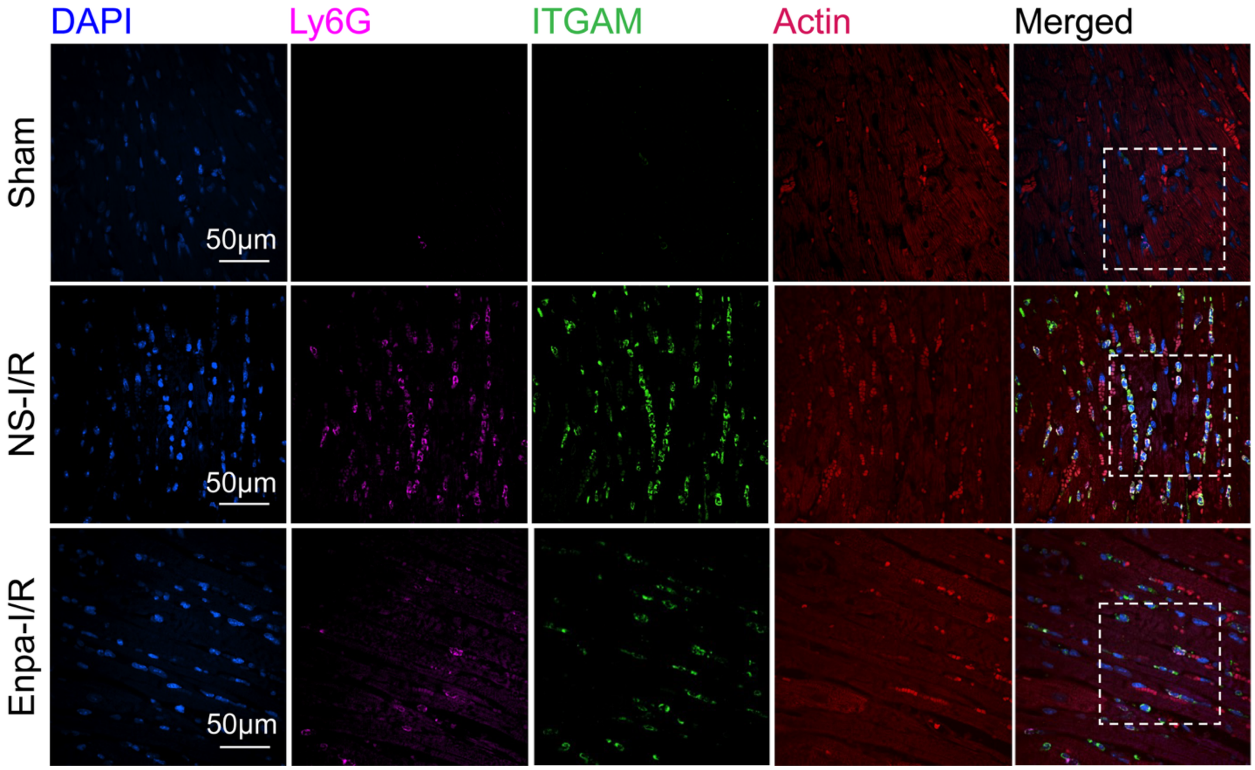


**Figure S6. ITGAM expression in neutrophils.** Fluorescent imaging of DAPI (Blue), Ly6G (magenta), ITGAM (green), and sarcomeric actin (red) on heart sections (60X). Dashed boxes indicate the regions shown at higher magnification in Fig. 3D - NE panel. NS, normal saline; Enpa, Enpatoran; I/R, ischemia-reperfusion; NE, neutrophil; DAPI, 4′,6-diamidino-2-phenylindole.

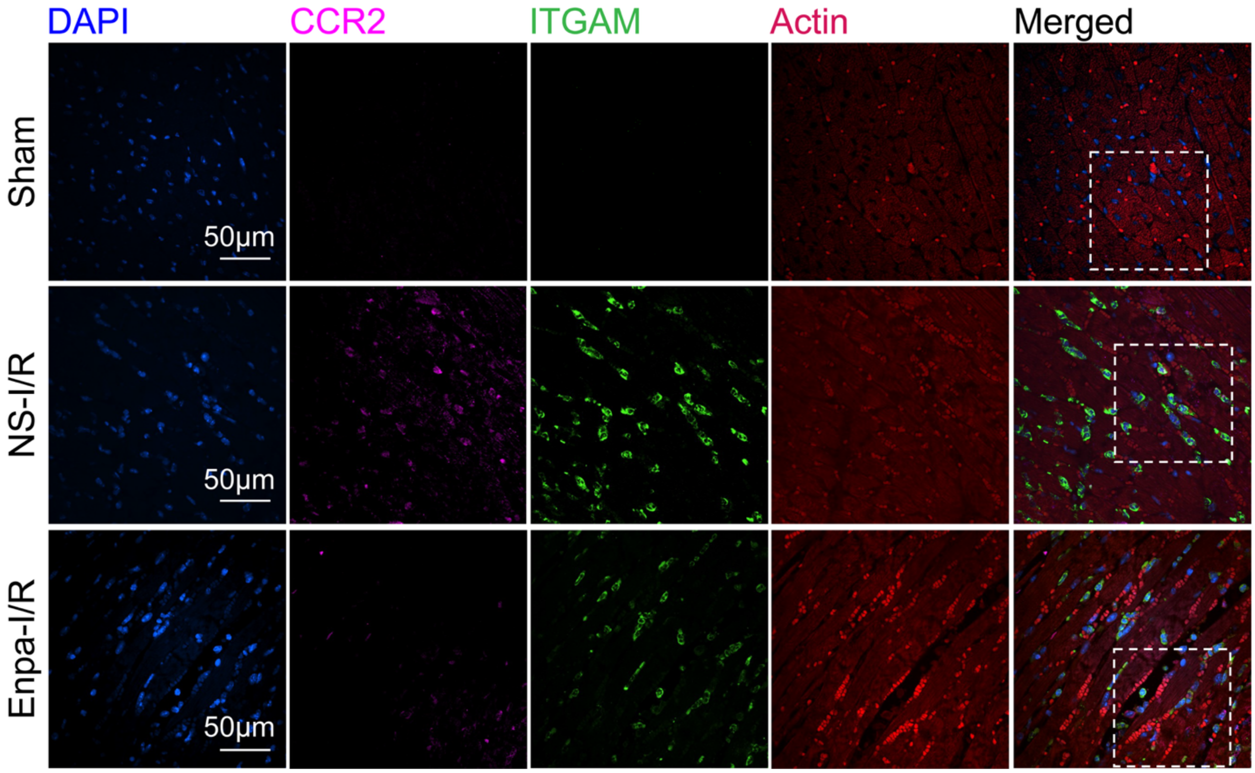


**Figure S7. ITGAM expression in monocytes.** Fluorescent imaging of DAPI (Blue), CCR2 (magenta), ITGAM (green), and sarcomeric actin (red) on heart sections (60X). Dashed boxes indicate the regions shown at higher magnification in Fig. 3D - Mo panel. NS, normal saline; Enpa, Enpatoran; I/R, ischemia-reperfusion; Mo, monocyte; DAPI, 4′,6-diamidino-2-phenylindole.

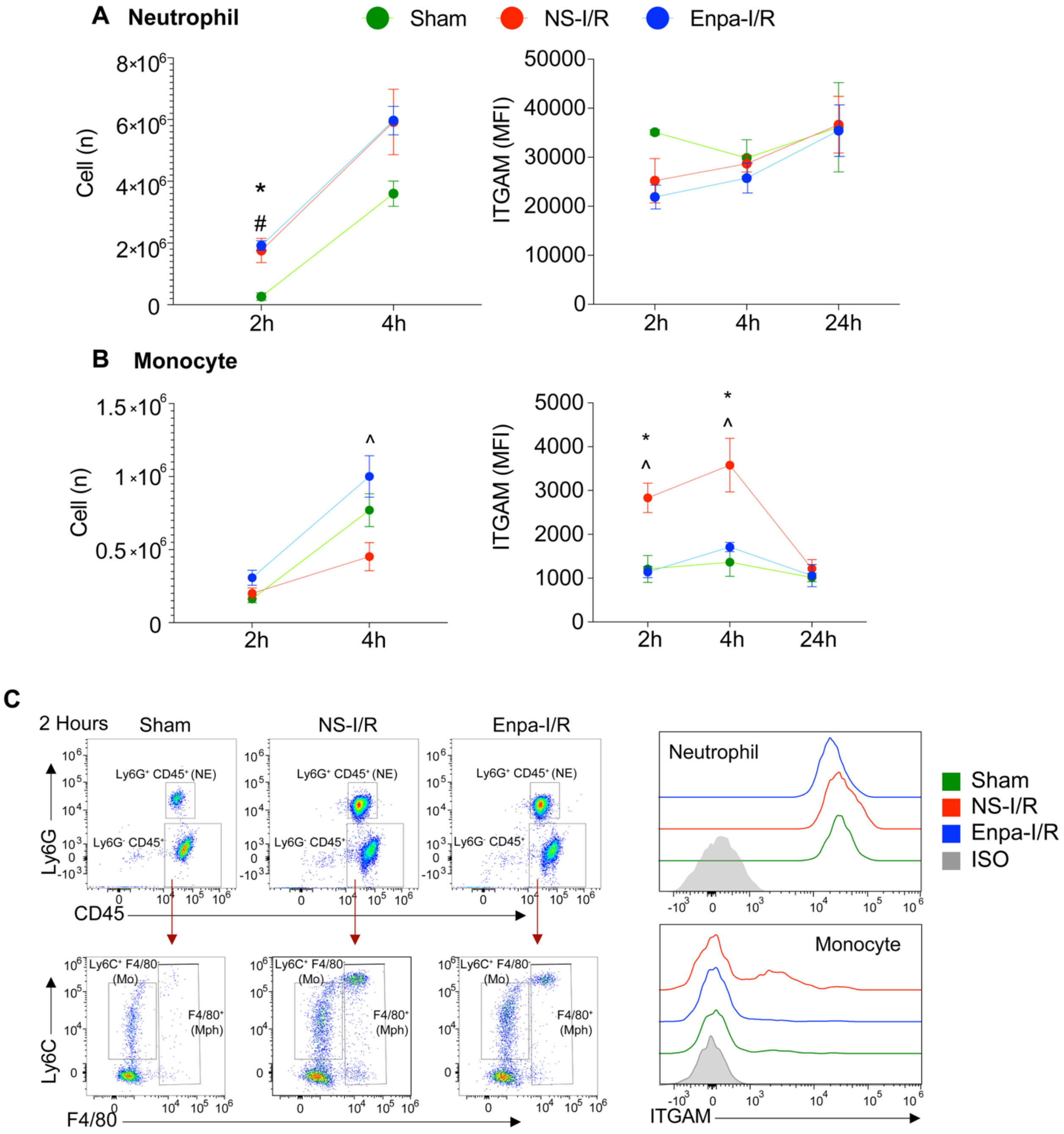


**Figure S8. Blood neutrophil and monocyte dynamics and ITGAM expression following myocardial I/R injury. A-B**, Flow cytometric quantification of circulating neutrophils (**A**) and monocytes (**B**) and ITGAM expression at 2h, 4h, and 24h after I/R injury, with Enpa or NS treatment. n=3-6 mice per time point. **C**, Representative gating strategy and histograms showing ITGAM expression in neutrophils and monocytes at 2h after I/R. Data are presented as mean ± SEM. One-way ANOVA was used for statistical comparisons among groups. *: Sham vs. NS-I/R, $P < 0.05$; #: Sham vs. Enpa-I/R, $P < 0.05$; ^: Enpa-I/R vs. NS-I/R, $P < 0.05$. NS, normal saline; Enpa, Enpatoran; I/R, ischemia-reperfusion; NE, neutrophil; Mo, monocyte; MFI, mean fluorescence intensity.

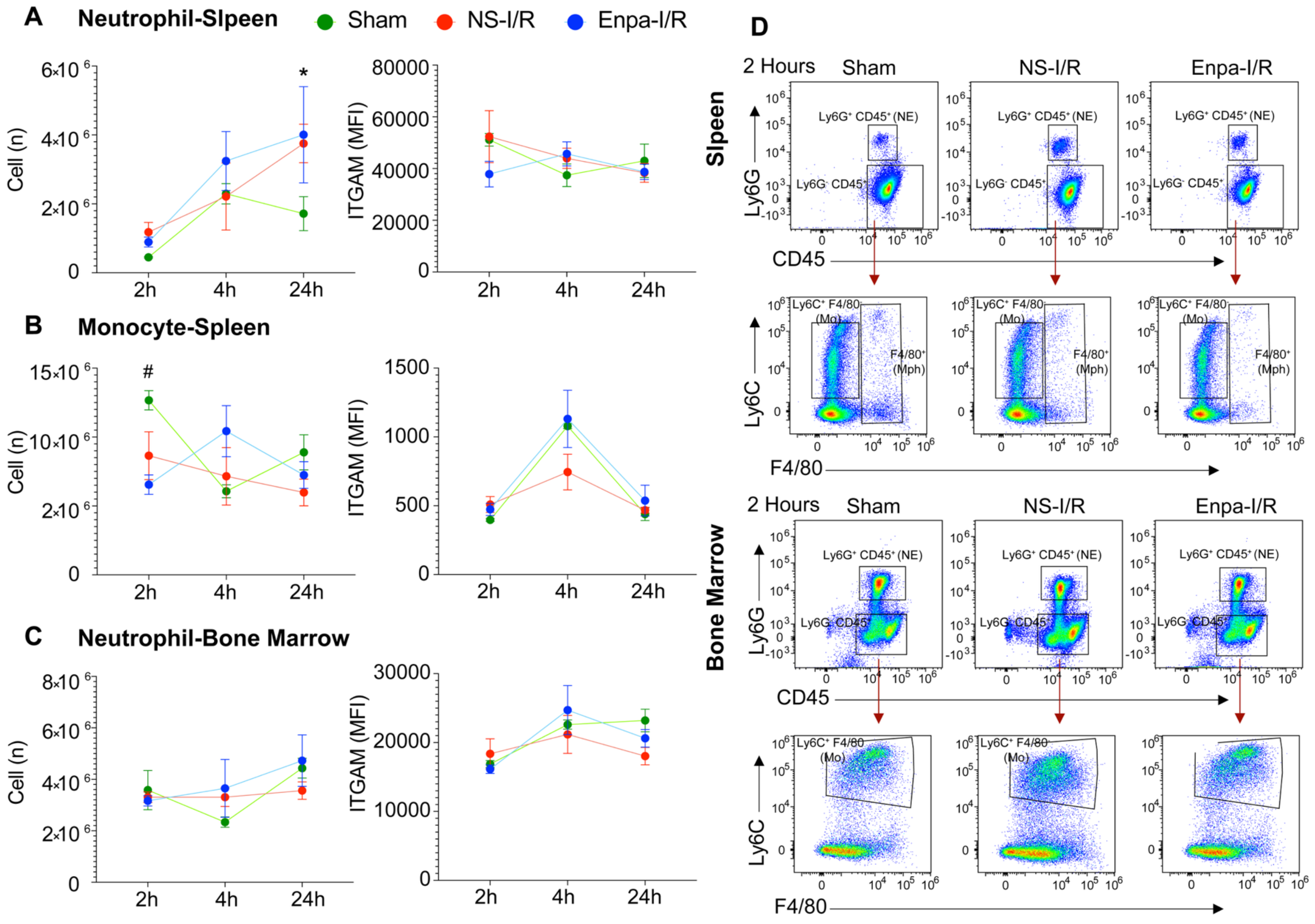

A
Neutrophil-Slpeen
Sham
NS-I/R
Enpa-I/R
Cell (n)
ITGAM (MFI)
2h
4h
24h
B
Monocyte-Spleen
C
Neutrophil-Bone Marrow
D
2 Hours
Slpeen
Bone Marrow
Ly6G⁺ CD45⁺ (NE)
Ly6G⁻ CD45⁺
CD45
Ly6G
Ly6C⁺ F4/80⁻ (Mo)
F4/80⁺ (Mph)
Ly6C
F4/80

**Figure S9. Spleen and bone marrow neutrophil and monocyte dynamics and ITGAM expression following I/R injury. A-C**, Flow cytometric quantification of spleen and bone marrow neutrophils (**A and C**) and spleen monocytes (**B**), and ITGAM expression at 2h, 4h, and 24h after ischemia-reperfusion (I/R) injury, with or without Enpa treatment. n=3-6 mice per time point. **D**, Representative gating strategy and histograms showing ITGAM expression in neutrophils and monocytes at 2h after I/R. NS, normal saline; Enpa, Enpatoran; I/R, ischemia-reperfusion; NE, Neutrophil; Mo, Monocyte. Data are presented as means ± SEM. One-way ANOVA was used for statistical comparisons among groups. *: Sham vs. NS-I/R, $P < 0.05$; #: Sham vs. Enpa-I/R, $P < 0.05$; ^: Enpa-I/R vs. NS-I/R, $P < 0.05$. NS, normal saline; Enpa, Enpatoran; I/R, ischemia-reperfusion; NE, neutrophil; Mo, monocyte; MFI, mean fluorescence intensity.

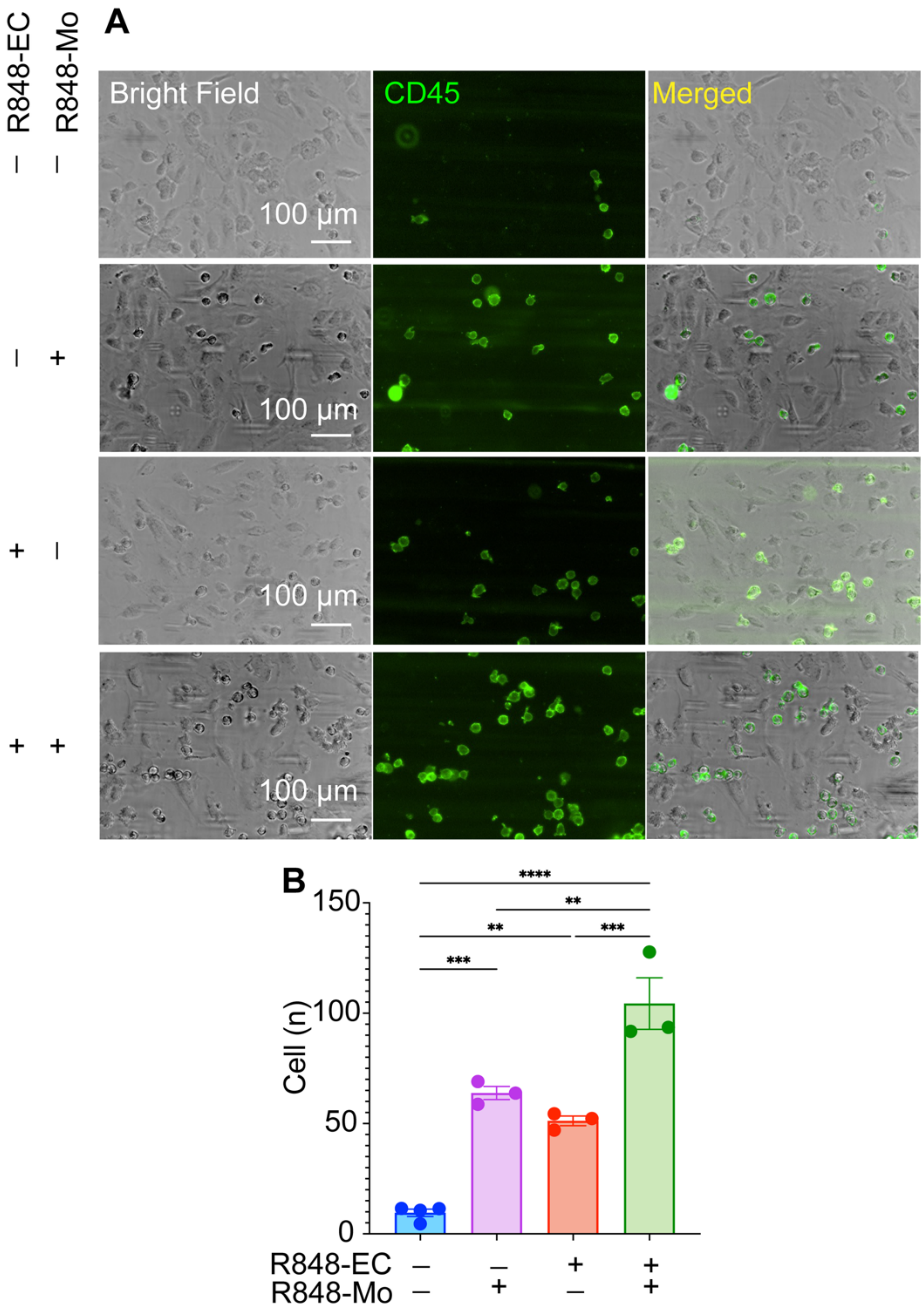


**Figure S10. Microfluidic adhesion assay of THP-1 adhesion to HCAEC.** Representative photomicrographs (**A**) and quantification (**B**) showing FITC anti-human CD45 antibody–labeled THP-1 cells (green) adhering to human coronary artery endothelial cells (HCAECs) within microfluidic channels. For each EC treatment condition, 3-4 corresponding channels were infused with THP-1 cells prepared under the corresponding treatment condition. Quantification of adherent monocytes was performed across 3-4 channels in one experiment. $CD45^+$ immune cells are shown in green. Adherent $CD45^+$ cells appear as discrete green puncta, while continuous green streaks reflect moving monocytes under flow conditions. nc, non-treated control; R, R848 (1µg/mL); Mo, monocytes; EC, endothelial cell.

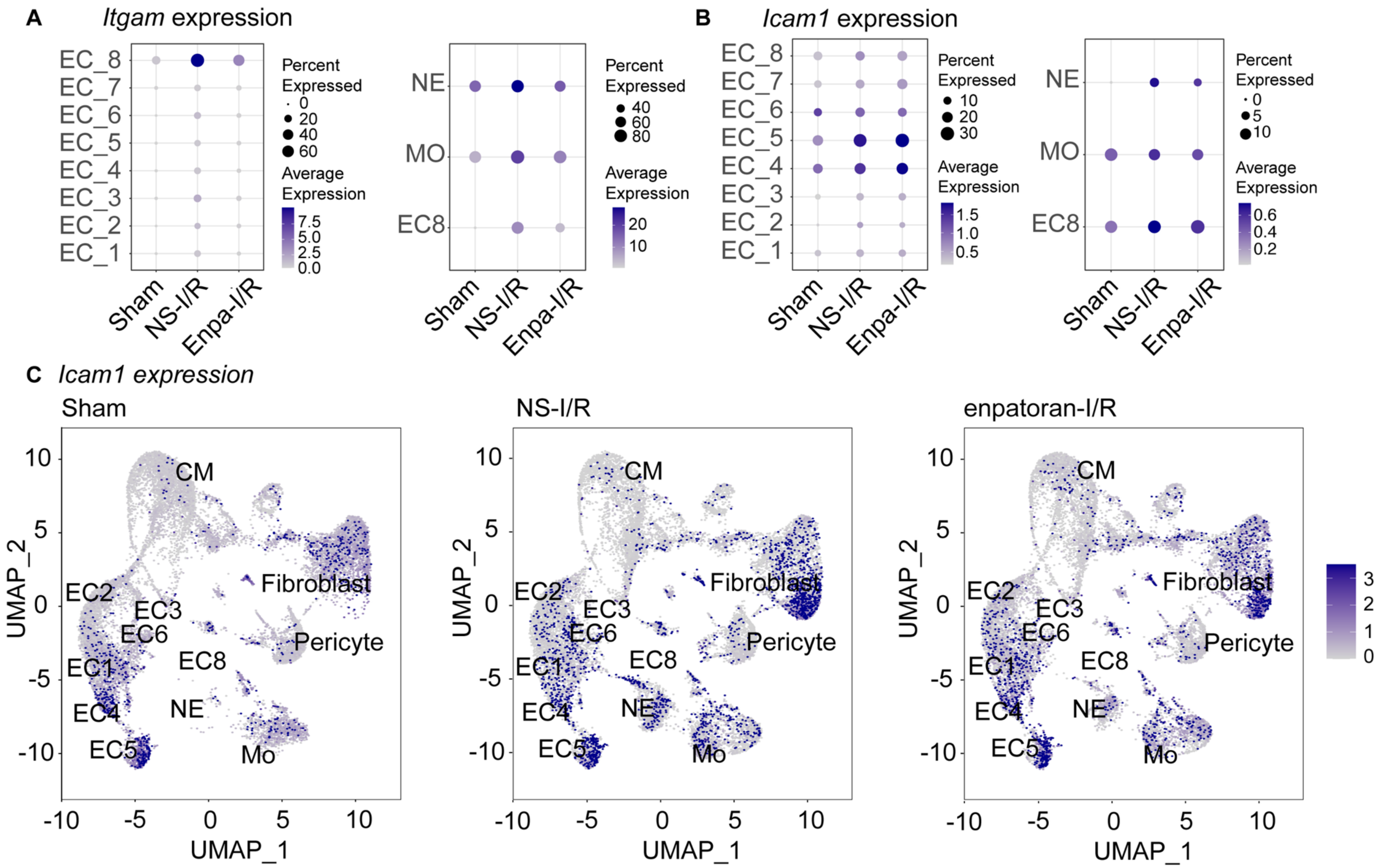

A
Itgam expression
EC_8
EC_7
EC_6
EC_5
EC_4
EC_3
EC_2
EC_1
Percent Expressed
0
20
40
60
Average Expression
7.5
5.0
2.5
0.0
Sham
NS-I/R
Enpa-I/R
NE
MO
EC8
Percent Expressed
40
60
80
Average Expression
20
10
B
Icam1 expression
Percent Expressed
10
20
30
Average Expression
1.5
1.0
0.5
Percent Expressed
0
5
10
Average Expression
0.6
0.4
0.2
C
Icam1 expression
Sham
NS-I/R
enpatoran-I/R
CM
Fibroblast
Pericyte
EC2
EC3
EC6
EC1
EC8
EC4
NE
EC5
Mo
UMAP_1
UMAP_2

**Figure S11. *Itgam* and *Icam1* expression level. A and B,** Dot plots showing *Itgam* and *Icam1* expression levels and the proportion of *Itgam*-expressing cells across endothelial subtypes (EC1–EC8), neutrophils (NE), and monocytes (Mo). **C,** UMAP plots showing the spatial distribution and *Icam1* expression in snRNA-seq data from Sham, NS-I/R, and Enpa-I/R mouse hearts. The color intensity represents normalized expression levels of *Icam1* across cardiac cell clusters. *Icam1* expression was increased in NS-I/R hearts compared with Sham and was reduced with Enpa treatment. I/R, ischemia-reperfusion; NS, normal saline; Enpa, Enpatoran; EC, endothelial Cell; Mo, monocyte; NE, neutrophils; UMAP, uniform manifold approximation and projection.

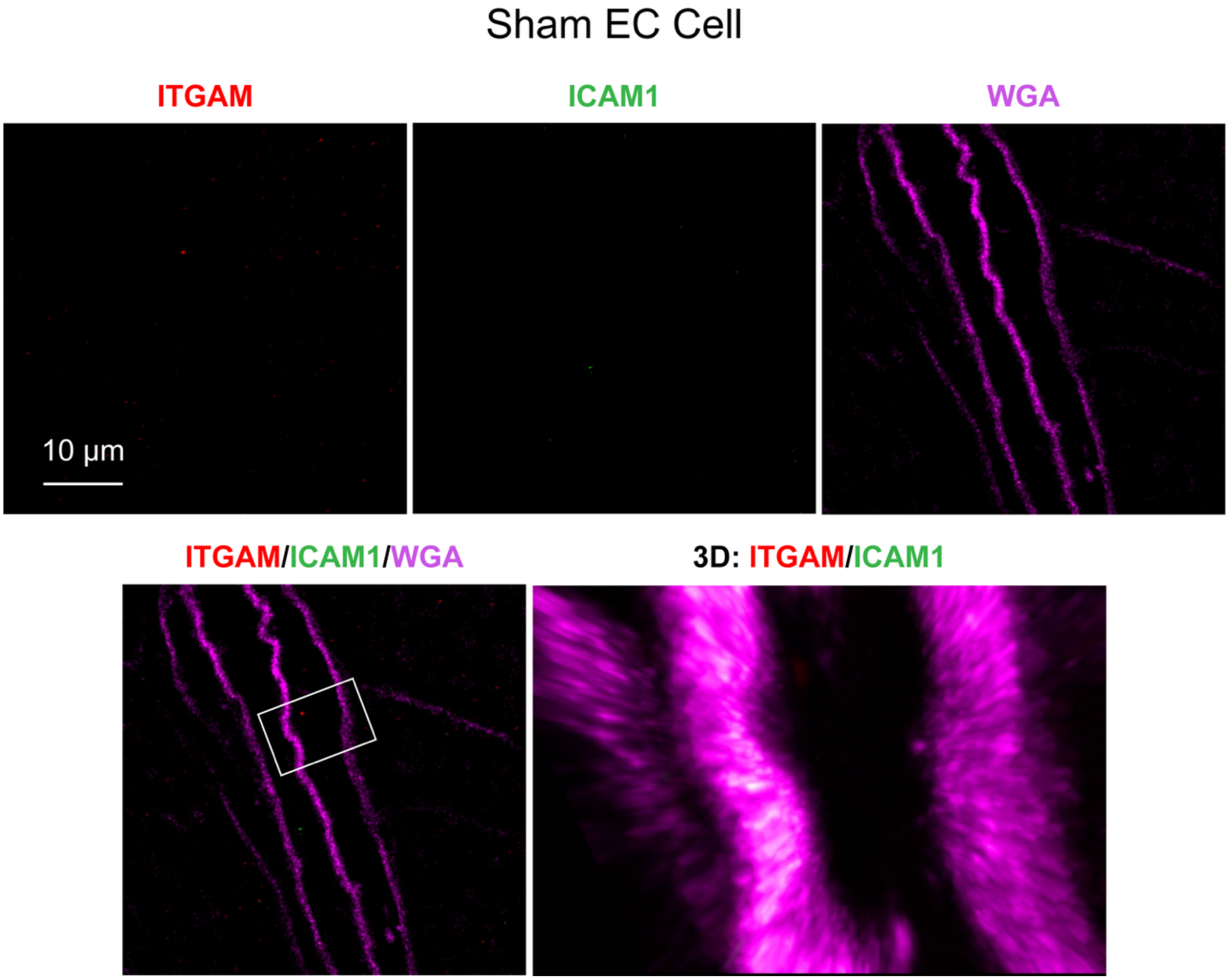


**Figure S12. STED super-resolution imaging of endothelial cells in the normal heart.** Cell membranes were imaged with Wheat Germ Agglutinin (WGA, magenta). There is a low level of ITGAM and ICAM-1 expression in the normal heart.

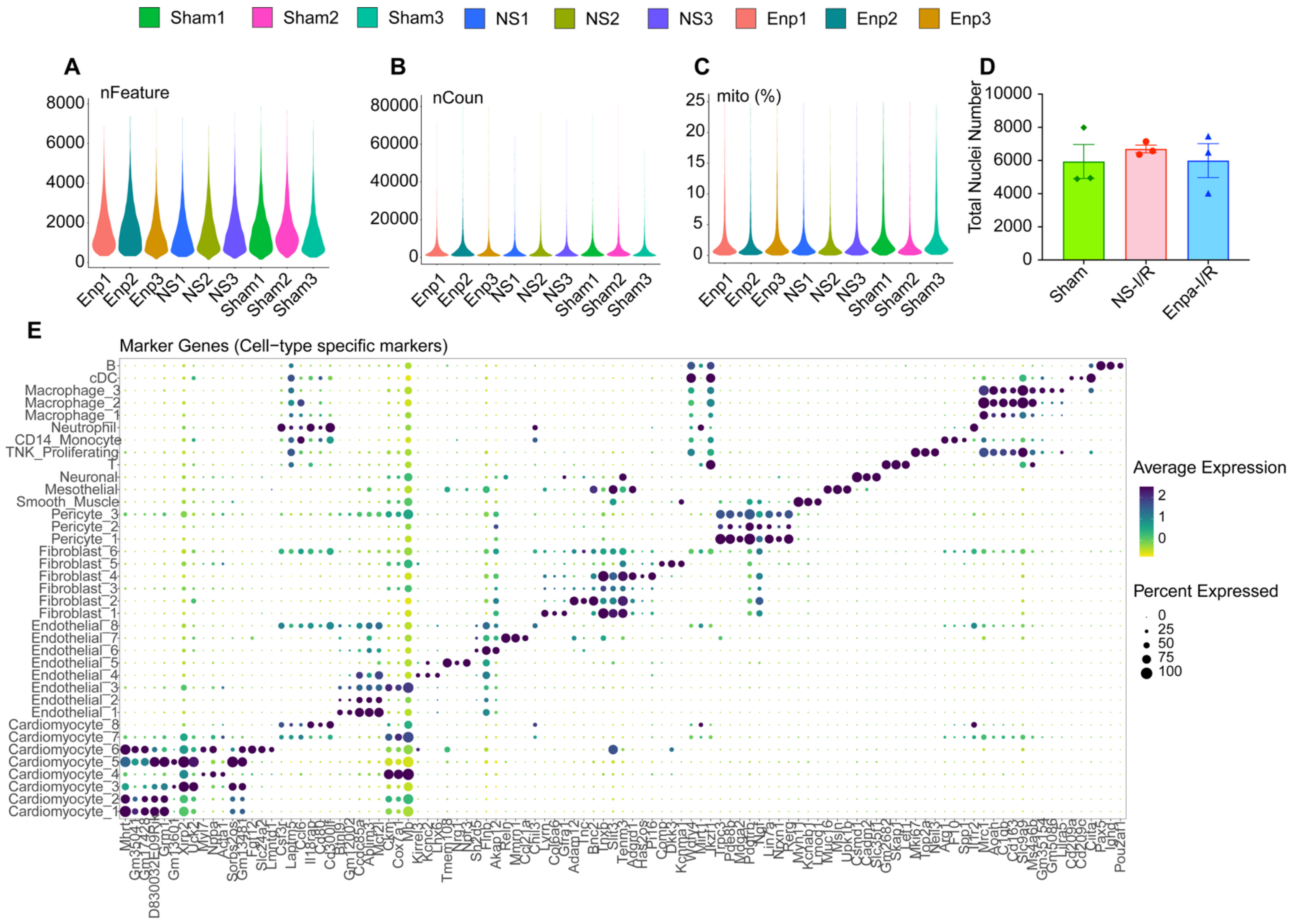

Sham1 Sham2 Sham3 NS1 NS2 NS3 Enp1 Enp2 Enp3
A
nFeature
B
nCoun
C
mito (%)
D
Total Nuclei Number
Sham NS-I/R Enpa-I/R
E
Marker Genes (Cell−type specific markers)
Average Expression
Percent Expressed

**Figure S13. Detailed snRNA-seq analysis data. A-C,** Quality control metrics for snRNA-seq across all samples, showing the number of detected genes per nucleus (*nFeature*, **A**), total RNA counts (*nCount*, **B**), and mitochondrial gene percentage (*mito%*, **C**); **D,** Quantification of the total number of nuclei analyzed per experimental condition (NS-Sham, NS-I/R, and Enpa-I/R); **E,** Cell cluster identification and validation based on cell-type-specific marker genes. The dot plot demonstrates the expression of canonical markers across distinct clusters, confirming clear separation. NS, normal saline; Enpa, Enpatoran; I/R, ischemia-reperfusion.
